# Clonal dynamics of haematopoiesis across the human lifespan

**DOI:** 10.1101/2021.08.16.456475

**Authors:** Emily Mitchell, Michael Spencer Chapman, Nicholas Williams, Kevin Dawson, Nicole Mende, Emily F Calderbank, Hyunchul Jung, Thomas Mitchell, Tim Coorens, David Spencer, Heather Machado, Henry Lee-Six, Megan Davies, Daniel Hayler, Margarete Fabre, Krishnaa Mahbubani, Fede Abascal, Alex Cagan, George Vassiliou, Joanna Baxter, Inigo Martincorena, Michael R Stratton, David Kent, Krishna Chatterjee, Kourosh Saeb Parsy, Anthony R Green, Jyoti Nangalia, Elisa Laurenti, Peter J Campbell

## Abstract

Age-related change in human haematopoiesis causes reduced regenerative capacity^1^, cytopenias^2^, immune dysfunction^3^ and increased risk of blood cancer. The cellular alterations that underpin the abruptness of this functional decline after the age of 70 years remain elusive. We sequenced 3579 genomes from single-cell-derived colonies of haematopoietic stem cell/multipotent progenitors (HSC/MPPs) across 10 haematologically normal subjects aged 0-81 years. HSC/MPPs accumulated 17 mutations/year after birth and lost 30bp/year of telomere length. Haematopoiesis in adults aged <65 was massively polyclonal, with high indices of clonal diversity and a stable population of 20,000–200,000 HSC/MPPs contributing evenly to blood production. In contrast, haematopoiesis in individuals aged >75 showed profoundly decreased clonal diversity. In each elderly subject, 30-60% of haematopoiesis was accounted for by 12-18 independent clones, each contributing 1-34% of blood production. Most clones had begun their expansion before age 40, but only 22% had known driver mutations. Genome-wide selection analysis estimated that 1/34 to 1/12 non-synonymous mutations were drivers, occurring at a constant rate throughout life, affecting a wider pool of genes than identified in blood cancers. Loss of Y chromosome conferred selective benefits on HSC/MPPs in males. Simulations from a simple model of haematopoiesis, with constant HSC population size and constant acquisition of driver mutations conferring moderate fitness benefits, entirely explained the abrupt change in clonal structure in the elderly. Rapidly decreasing clonal diversity is a universal feature of haematopoiesis in aged humans, underpinned by pervasive positive selection acting on many more genes than currently identified.

## Introduction

The age-related mortality curve for modern humans is an extreme outlier among species across the tree of life, with an abrupt increase in standardised mortality rates after the average lifespan^4^, leading to surprisingly low variance in age at death^5^. Behavioural, medical and environmental interventions have driven down early and extrinsic causes of mortality, thereby unmasking a substantial burden of intrinsic, age-associated disease. Studies of ageing at the cellular level have demonstrated that accumulation of molecular damage across the lifespan is gradual and lifelong, including telomere attrition^6–8^, somatic mutation^9–11^, epigenetic change^12^ and oxidative or replicative stress^13,14^. It remains unresolved how such gradual accumulation of molecular damage can translate into an abrupt increase in mortality after the age of 70 years.

The haematopoietic system is an interesting organ for studying human ageing. It manifests several age-associated phenotypes including anaemia; loss of regenerative capacity, especially in the face of insults such as infection, chemotherapy or blood loss; and increased risk of blood cancer. One aspect of age-related change in human haematopoietic stem cells (HSCs) that has been an area of intense study is ‘clonal haematopoiesis’^15^. This is defined by single-cell-derived expansions in blood, most commonly associated with mutations in genes recognised as causal for myeloid neoplasms, so-called driver mutations. Clonal haematopoiesis increases with age, reaching 10-20% prevalence^16–20^ or even higher^21^ after 70 years. Data are now emerging that many elderly individuals have evidence of clonal expansions even in the absence of known driver mutations^22–24^. Cellular changes of ageing have primarily been studied in mice^25^. While the numbers of phenotypic HSCs increase in aged mice, aged HSCs produce fewer mature progeny both *in vitro* and in serial transplantation compared to young HSCs^26–28^.

HSCs accumulate somatic mutations linearly with age, with each cell acquiring a new mutation every 2-3 weeks on average throughout life^10,29^. Whole-genome sequencing of colonies grown from single cells enables comprehensive identification of these somatic mutations and reconstruction of lineage relationships among cells^30^ (**Fig. 1a**). We used this approach to study the clonal dynamics of human haematopoiesis across the lifespan, sequencing whole genomes of 3579 single-cell-derived colonies from 10 healthy individuals.

**Fig. 1.**
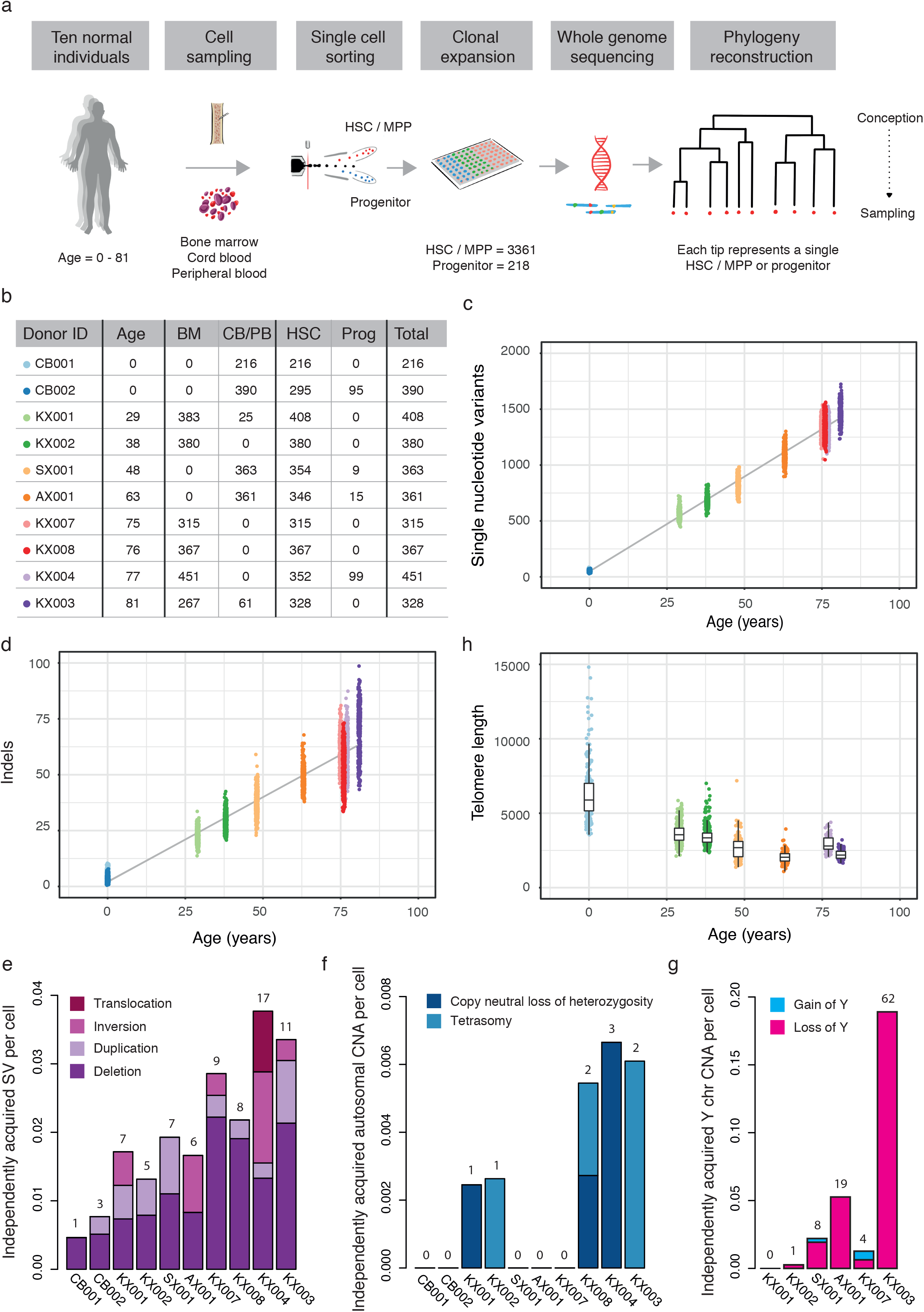
Mutational burden in normal HSC/MPPs. **a**, Experimental approach. **b**, Table showing the age and numbers of colonies sequenced from each tissue and cell type for each donor in the study. **c**, Burden of single nucleotide variants across the donor cohort. The points represent individual HSC/MPP colonies (n = 3361) and are coloured by donor. The grey line represents a regression of age on mutation burden, with 95% CI shaded. **d**, Burden of small indels across the donor cohort. The points and line are depicted as in c. **e**, Barplot of the number of independently acquired structural variants per colony sequenced in each donor. The absolute number of SVs is at the top of each bar. **f**, Barplot of the number of independently acquired autosomal copy number aberrations per colony sequenced in each donor. The absolute number of CNAs is at the top of each bar. **g**, Barplot of the number of independently acquired Y chromosome copy number aberrations sequenced in each male donor. The absolute number of CNAs is at the top of each bar. **h**, Telomere length across the donor cohort, including only those samples sequenced on the HiSeq X10 platform. Each point represents a single HSC/MPP colony. The boxes overlaid indicate the median and interquartile range and the whiskers denote the range. Two outlying points for CB001 are not shown (telomere lengths 16,037bp and 21,155bp).

### Whole genome sequencing of HSC-derived colonies

We obtained samples from 10 individuals, with no known haematological disease, aged between 0 and 81 years (**Table S1**). One subject had inflammatory bowel disease treated with azathioprine (KX002, 38-year male) and one had selenoprotein deficiency^31^, a genetic disorder not known to impact HSC dynamics (SX001, 48 year male). The source of stem cells was cord blood (CB) for the two neonates, and bone marrow and/or peripheral blood for adult donors (**Fig. 1b**). Bone marrow samples were obtained peri-mortem, allowing sampling of large volumes (50-80mL) from multiple vertebrae.

For all individuals, single phenotypic haematopoietic stem cell/multipotent progenitors (HSC/MPPs: Lin-, CD34+, CD38-, CD45RA-) were flow-sorted^32^ and cultured (**Extended Fig. 1a**). Overall, 42-89% of sorted HSC/MPPs produced colonies, meaning that the colonies were a representative sample of the HSC/MPP population in each individual (**Extended Fig. 1b**). For four individuals, haematopoietic progenitor cells (HPCs) (Lin-, CD34+, CD38+) were also isolated for comparison of mutation burden between HSCs and HPCs.

We performed whole-genome sequencing at an average sequencing depth of 14x on 224-453 colonies per individual (**Fig. 1b**). We excluded 17 colonies with low coverage, 34 technical duplicates and 7 colonies derived from more than a single cell (**Extended Fig. 2a-c**). The final dataset comprised whole genomes from 3579 colonies, of which 3361 were single HSC/MPP-derived and 218 were single HPC-derived. Raw mutation burdens were corrected for sequencing depth using asymptotic regression (**Extended Fig. 2d**). Single base substitution spectra were consistent with published results^29,30,33^ (**Extended Fig. 2e-f**). Phylogenetic trees for each adult individual were constructed from the patterns of shared and unique somatic mutations (**Extended Fig. 3a-d**). Benchmarking of the phylogenies included an assessment of their internal consistency, stability across phylogenetic inference algorithms, and robustness to bootstrapping approaches (**Supplementary Fig. 3**). All code for the variant filtering, signatures, phylogenetic reconstruction, benchmarking and downstream analyses are available (**Supplementary Code**).

### Mutation burden and telomere lengths

Consistent with published data^29,34^, point mutations accumulated linearly in HSC/MPPs throughout life, at a rate of 16.8 substitutions/cell/year (CI_95%_=16.5-17.1; **Fig. 1c**) and 0.71 indels/cell/year (CI_95%_=0.65-0.77; **Fig. 1d**). We found no significant difference in mutation burden between HSC/MPPs and HPCs (**Extended Fig. 4a**). Structural variants were rare, with only 1-17 events observed in each individual, mostly deletions, correlating with age (**Fig. 1e; Extended Fig. 4b; Table S3**). Autosomal copy number aberrations were rare at all ages and comprised either copy-neutral loss-of-heterozygosity events or tetrasomies (**Fig. 1f**). In contrast, Y chromosome copy number changes were frequent in males, increasing with age as previously shown^35^ (**Fig. 1g**).

We estimated telomere lengths for the 1505 HSC/MPP colonies from 7 individuals sequenced on Hiseq X10^36^ (**Fig. 1h**). As previously reported^8,37,38^, telomere lengths decreased steadily with age, at an average attrition rate of 30.8 bp/year in adult life (CI_95%_=13.2-48.4), which is close to published estimates of 39 bp/year from bulk granulocytes^8^. By sequencing single-cell-derived colonies, we can estimate the variance and distribution in telomere lengths among cells with a resolution not possible for bulk populations. In cord blood and adults aged <65, a small proportion of HSC/MPPs had unexpectedly long telomeres, as assessed using several criteria for outliers, a proportion that reduced in frequency with age (**Extended Fig. 4c**). Given that telomeres shorten at cell division, these outlier cells have presumably undergone fewer historic cell divisions. A rare population of dormant HSCs dividing infrequently has been described in the mouse^39–41^ and our telomere data would be consistent with an analogous population in humans, especially children and young adults.

### Qualitative change in HSC population structure after 70 years of age

The phylogenetic trees generated here depict the lineage relationships among ancestors of the stem and progenitor cells sequenced. Given the consistent, linear rate of mutation accumulation across the lifespan, we can scale the raw phylogenetic trees (**Extended Fig. 3d**) to chronological time to study clonal dynamics of HSC/MPPs across the lifespan (**Figs 2-3**). Branch-points in the tree, so-called ‘coalescences’, define historic phylogenetic relationships between stem cells. The chance of capturing coalescent events in a phylogeny of randomly sampled stem cells depends on both the population size and time between symmetric self-renewal divisions. In taxonomy, a ‘clade’ is defined as the group of organisms descended from a single common ancestor – in the context of somatic cells, this represents a clone, and its size can be estimated from the fraction of colonies derived from that ancestor. Here, we define an ‘expanded clade’ as a post-natal ancestral lineage whose descendants contributed >1% of colonies at the time of sampling.

**Fig. 2.**
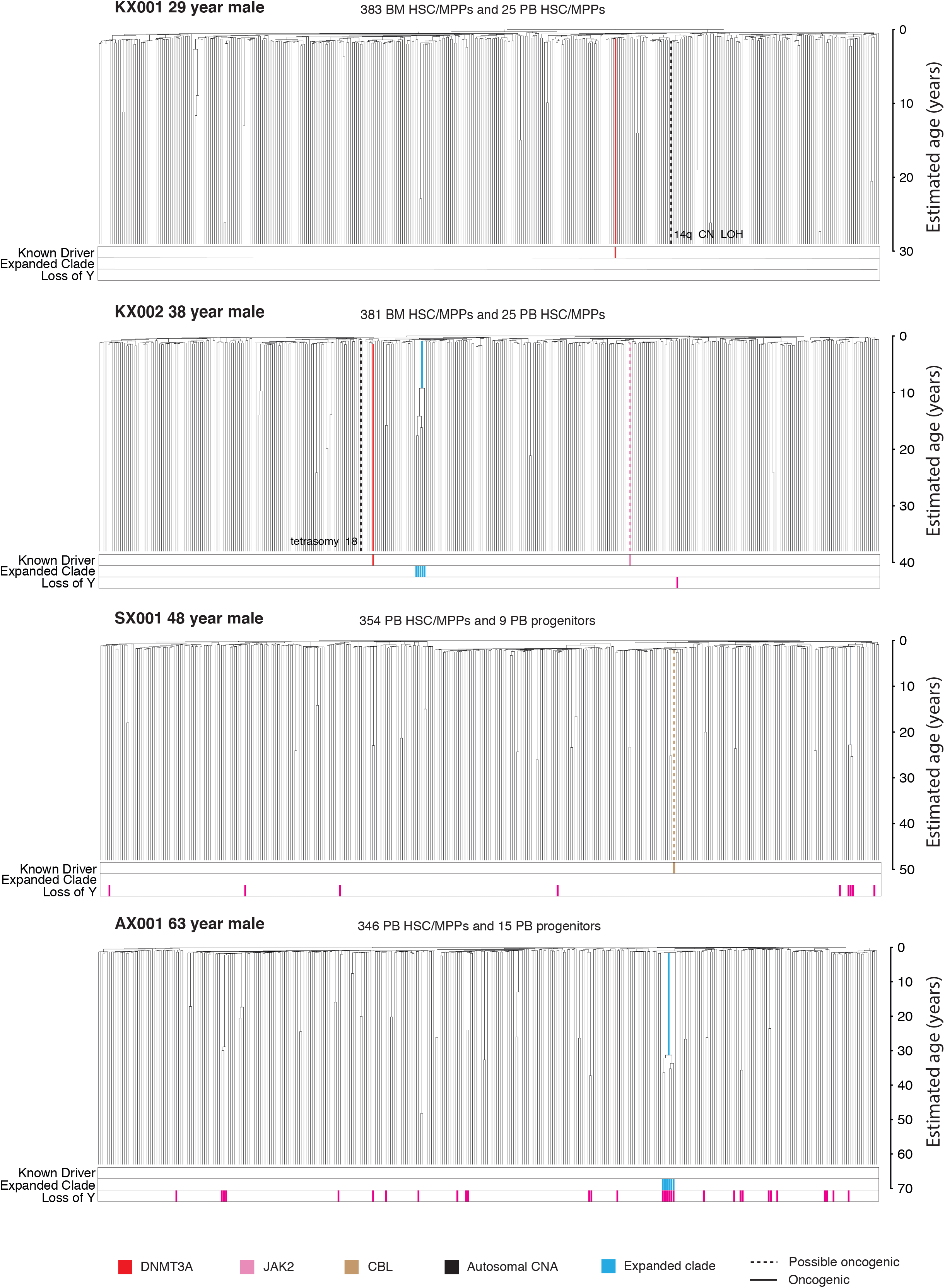
HSPC phylogenies for the four youngest adult donors. Each tip on a phylogeny represents a single HSPC, with the respective numbers of colonies of each cell and tissue types recorded at the top. Branches with known oncogenic drivers (solid line) and possible oncogenic drivers (dashed line) in one of our top 17 clonal haematopoiesis genes (**Table S4**) are highlighted and coloured by gene. Essential splice site mutations annotated by *dNdScv* in these genes are included. Branches with autosomal CNAs are highlighted with a black dashed line. A heatmap at the bottom of each phylogeny highlights cells from ‘known driver’ clades in red, ‘expanded clades’ (defined as those with a clonal fraction > 1%) in blue and cells with loss of Y in pink. The y axis is estimated age in years which allows timing of the coalescent (or branching events) in the tree.

**Fig. 3.**
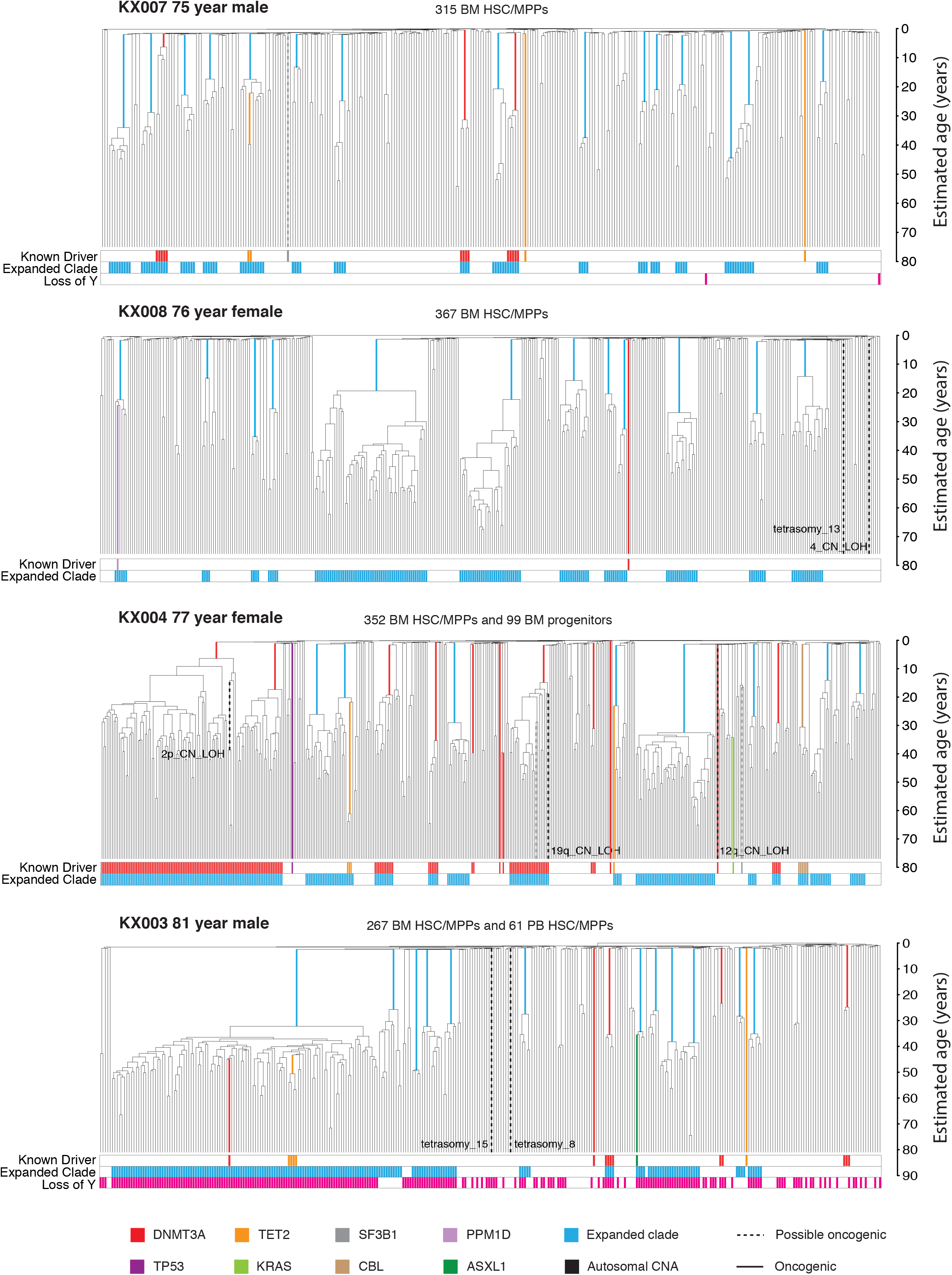
HSPC phylogenies for the four elderly adult donors. Each tip on a phylogeny represents a single HSPC, with the respective numbers of colonies of each cell and tissue types recorded at the top. Branches with known oncogenic drivers (solid line) and possible oncogenic drivers (dashed line) in one of our top 17 clonal haematopoiesis genes (**Table S4**) are highlighted and coloured by gene. Essential splice site mutations annotated by dNdScv in these genes are included. Branches with autosomal CNAs are highlighted with a black dashed line. A heatmap at the bottom of each phylogeny highlights cells from ‘known driver’ clades in red, ‘expanded clades’ (defined as those with a clonal fraction > 1%) in blue and cells with loss of Y in pink. The y axis is estimated age in years which allows timing of the coalescent (or branching events) in the tree.

Phylogenetic trees of the 4 adults aged <65 showed that healthy young and middle-aged haematopoiesis is highly polyclonal (**Fig. 2**). Despite sequencing 361-408 colonies per individual, we found at most a single expanded clade in each sample, and the two we did observe contributed <2% of all haematopoiesis in those individuals. Known driver point mutations were sparse, with 2 in *DNMT3A* and 1 possible oncogenic variant each in *CBL* and *JAK2*, none of which occurred in the two expanded clades.

Phylogenetic trees for the 4 adults aged >70 were qualitatively different (**Fig. 3**), with an oligoclonal pattern of haematopoiesis. In each elderly individual, we found 12-18 independent clones established between birth and the age of 40 that each contributed between 1% and 34% of colonies sequenced, most in the 1-3% range. Collectively, these clones summed to a significant proportion of all blood production in our elderly research subjects – between 32% and 61% of all colonies sequenced derived from the expanded clades.

Most of the oligoclonal blood production in the elderly individuals could not be accounted for by known driver mutations. Although mutations in *DNMT3A, TET2* and *CBL* did occur in some expansions, mutations in the top 17 myeloid driver genes could only explain 10/58 expanded clades with a clonal fraction >1%. Only 3 additional mutations were identified if we extended to a wider set of 92 genes implicated in myeloid neoplasms^42^ (**Table S5**), still leaving 45 clonal expansions unexplained.

### Trajectory of HSC/MPP population size in young individuals

The frequency of branch-points in phylogenetic trees in a neutrally evolving, well-mixed population of somatic cells is primarily determined by the product of population size and time between symmetric self-renewal cell divisions (*Nτ*) – both smaller populations and more frequent symmetric divisions increase the density of coalescences. In young adults, where clonal selection has had minimal impact on the phylogenetic tree structure, we can exploit this property to estimate the lifelong trajectory of population size dynamics^43^ (**Fig. 4a**). There is a rapid increase in estimated *Nτ in utero*, reaching a plateau during infancy and early childhood. This is evidenced by a high density of coalescences in the first ∼50 mutations of molecular time (**Fig. 2**), noting that the cord blood colonies we sequenced had a mean of 55 mutations.

**Fig. 4.**
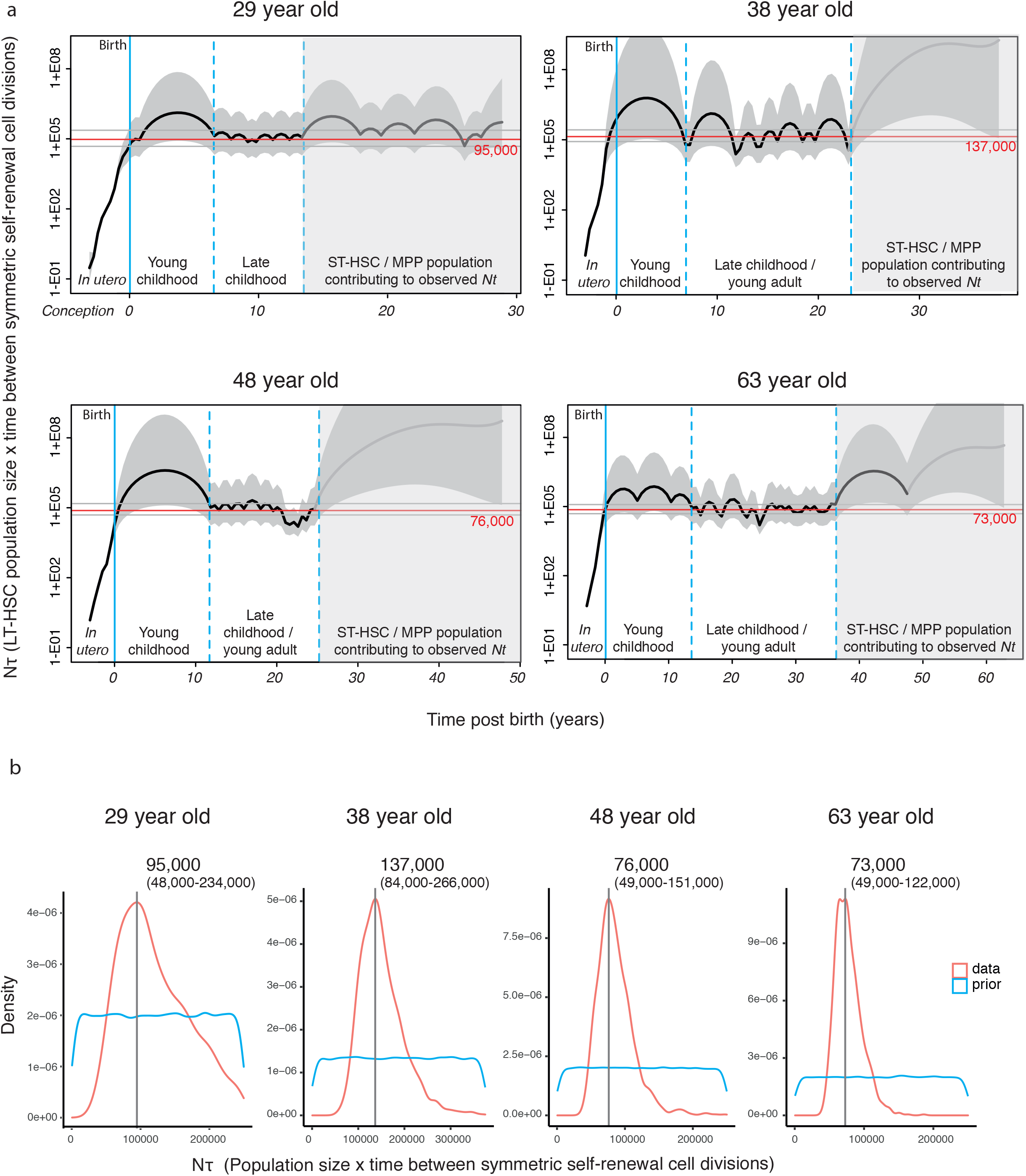
Estimating N*τ* in the human LT-HSC compartment. **a**, *Phylodyn* plots illustrating the trajectory of *Nτ* for human LT-HSCs in the four adult donors aged <65. The black line represents the trajectory of LT-HSC *Nτ*, with the shaded grey area on either side representing the 95% credibility interval. The solid blue line is the time of birth. The dashed blue lines enclose the region of time in each individual where the trajectory is at the late childhood/young adult level. The shaded region of the plots represents the period of time prior to sampling over which it is likely that ST-HSC/MPPs are contributing to the observed *Nτ*. The trajectory line is shaded dark grey in the time period where coalescent events are occurring and the trajectory likely represents the combined LT-HSC and ST-HSC/MPP *Nτ*. The trajectory line is shaded light grey where there is a complete absence of coalescent events and the estimates are highly inaccurate. The red line shows the Bayesian (maximum posterior density) estimate of *Nτ*. **b**, Results from approximate Bayesian inference of population size over the first (non-shaded) part of life for each individual. The blue line represents the prior density of *Nτ* and the red line represents the posterior density. The vertical grey line is the peak *Nτ* for each donor. The peak *Nτ* with 95% confidence limits is written at the top of each plot.

Using both phylodynamic and Approximate Bayesian Computation modelling, we estimate that steady-state haematopoiesis plateaus at an *Nτ* of about 100,000 HSC-years (**Fig. 4a-b**). This was consistent across all 4 young adults, with individual estimates and 95% confidence intervals in the range 50,000–250,000, consistent with published estimates^30,44^. In order to estimate *N* (the HSC population size), we require an estimate of *τ*, the time between symmetric HSC self-renewal divisions. Previous estimates of *τ* for HSCs range between 0.6 and 6 years^8,30,45^. Telomeres in HSCs shorten at mitosis by 30-100bp per cell division^46^, and therefore provide an independent means to estimate the number of historic cell divisions an HSC has undergone. We estimated that HSC/MPP telomeres shorten at a rate of 30 bp/year (**Fig. 1h**) – this bounds the number of symmetric cell divisions as at most 1-2 divisions per year. Two recent studies in mice in mice have shown that symmetric divisions predominate within the HSC pool, accounting for 80-100% of all HSC divisions^47,48^. Taken together, then, these data are most consistent with adult haematopoiesis being maintained by a population of 20,000-200,000 long-term HSCs.

In the young individuals, and indeed the four elderly individuals, we noted a sparsity of branch-points towards the tips of the phylogenetic trees compared to earlier ages – this consistently emerged about 10-15 years of molecular time before sampling, irrespective of the age of the research subject. We hypothesise this pattern arises from a large, short-term HSC/MPP compartment in humans that contributes to haematopoiesis for 10-15 years. Due to its short-term contribution, the effects of a short-term HSC compartment would only be evident at the tips of the phylogenetic tree, with the earlier branches of the lineage tree more faithfully reflecting dynamics in the long-term HSC compartment. A substantially larger population size for these short-term HSC/MPPs would explain why the density of coalescences decreases towards the tips of the trees (**Extended Fig. 5a-c**). The mouse ST-HSC/MPP compartment is known to sustain steady-state haematopoiesis without significant input from the long-term HSC compartment for many months^49–51^, a similar proportion of a mouse’s lifespan as 10-15 years is for a human.

### HSC dynamics in the elderly

We observed a qualitative change in the population structure of HSCs in the elderly, with an abrupt increase in the frequency of expanded clones and the total fraction of haematopoiesis they generate (**Fig. 5a**). Measures of clonal diversity, such as the Shannon Index, showed a precipitous decline in diversity after the age of 70 (**Figure 5b**). Although this change was only evident in the elderly, the branch-points in the phylogenetic tree that underpinned the clonal expansions occurred much earlier in life, between childhood and middle-age. There are several possible explanations for these observations, including age-related changes in population size, changes in spatial patterning of HSCs and cell-autonomous variation in selective fitness.

**Fig. 5.**
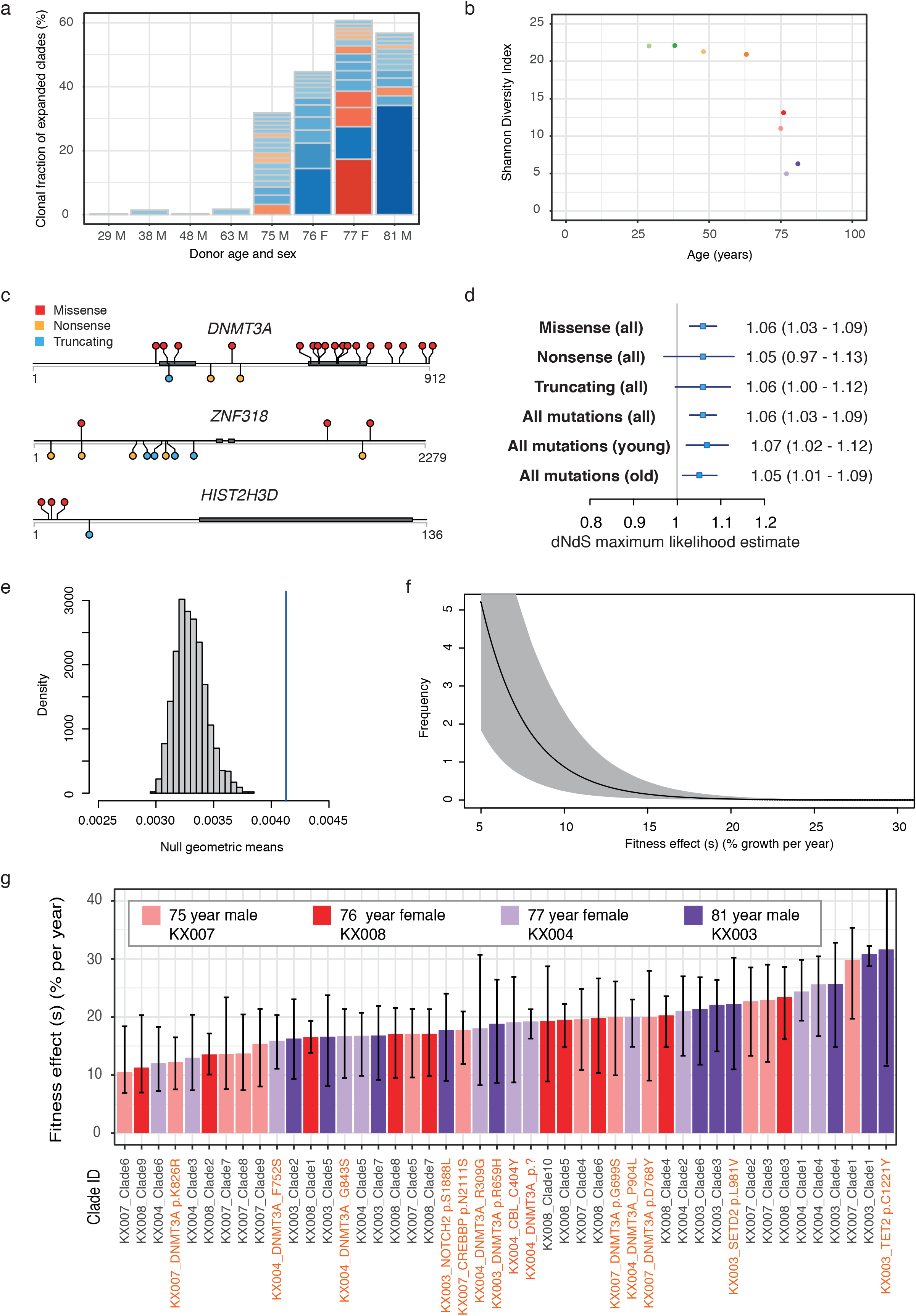
Widespread positive selection in the HSC/MPP compartment of normal individuals. **a**, Stacked barplot to show number and size of clones with a clonal fraction >1% per individual. Shades of blue denote clones with no known driver and shades of red denote those with a known myeloid gene driver. **b**, Shannon diversity index calculated for each phylogeny. The Shannon diversity index is a mathematical measure of species diversity that takes into account both the abundance and evenness of the species present. Here the number of species is the number of lineages present at 100 mutations of molecular time (equivalent to the first few years after birth) and abundance is the number of samples within the clade derived from that lineage. **c**, Lolliplot plots to show the sites of variants in the dataset in the three genes under significant positive selection according to dN/dS. Thick grey bars denote locations of conserved protein domains. **d**, dN/dS maximum likelihood estimates for missense, nonsense, truncating and all mutations in the complete dataset (n = 25,888 coding mutations) and for all mutations in the young (individuals aged < 65 year) and old (individuals aged > 75 years) datasets analysed separately. The boxes show the estimate with whiskers showing the 95% CI. The numbers to the left give the numeric values for the estimates with 95%CI in brackets. **e**, Results of a randomisation / Monte Carlo test to define the null expected distribution of clade size for cells with loss of Y. This null distribution of geometric means from 2000 simulations is shown (histogram) together with the observed geometric mean of clades with Y loss (vertical blue line). The observed value significantly outlies the expected distribution showing that clades with Y loss are significantly larger than would be expected by chance. **f**, Distribution of fitness effects for the ‘driver’ mutations entering the HSC population, as determined by ABC modelling approach. The point estimate for the shape and rate parameters of the gamma distribution were shape = 0.73 and rate = 33 (these are the univariate marginal maximum posterior density estimates; **Supplementary Fig. 12**). **g**, Fitness effects within the HSC/MPP compartment are estimated for clades with causal driver mutations containing 4 or more HSC/MPP colonies (percent additional growth per year). Fitness effects are also estimated for expanded clades containing 5 or more HSC/MPP colonies (percent additional growth per year). Clade numbers are illustrated on the phylogenies in **Extended Fig. 9c**.

Using Approximate Bayesian Computation^52,53^, we explored whether changes in population size, most likely a population bottleneck occurring in mid-life, could explain the observed trees (**Extended Fig. 6a**). However, although these simulations were able to accurately recapitulate the phylogenies observed in the individuals aged <65, even the 1% most closely-matching phylogenies from these simulations poorly replicated those observed in the elderly (**Extended Fig. 7a-b**). Although we could generate phylogenies with the same number of mid-life coalescences, we were unable to emulate the observed clade size distribution. Essentially, our observed trees show marked asymmetry among clones, with a few large clades but also numerous ‘singleton’ branches. In contrast, the simulated trees generated by variable population sizes showed a more even distribution of coalescences among clades.

Theoretically, altered spatial patterning of HSCs could also explain the changes in phylogeny – if, for example, HSCs circulated less frequently with ageing, the bone marrow population would be less well mixed and therefore show increased density of coalescences in a bone marrow sample obtained from just a few vertebrae, as studied here. To address this, we sequenced both marrow and peripheral blood HSCs from the 81-year old individual. The peripheral blood colonies recaptured most of the expanded clades evident in marrow and at similar clonal fractions, suggesting that spatial segregation is not sufficient to explain the changes with ageing (**Extended Fig. 6b**).

### dN/dS evidence for pervasive positive selection

The asymmetry of population structure in the elderly suggests that clone-specific factors could lead to differential expansion among HSCs. Positively selected driver mutations are one, but not the only, possible cause of clone-specific variation in expansion rates – epigenetic change and critical telomere shortening, for example, could also contribute. Indeed, it is striking that previously known driver mutations were present in only a minority (22%) of expanded clades. Furthermore, it is unclear whether driver mutations acquired randomly throughout life could lead to such a qualitative change in population structure after the age of 70. To address these questions, we used two approaches: one based on genetic analysis of selection and one based on Approximate Bayesian Computation models of drivers in haematopoiesis.

For the genetic analysis, we considered synonymous mutations as selectively neutral, and used their rate to quantify whether non-synonymous mutations occurred at equivalent or elevated rates (dN/dS ratio, with dN/dS>1.0 denoting a tilt towards positive selection) – this was undertaken gene-by-gene and across all coding genes collectively^54^. Estimating dN/dS gene-by-gene identified three genes under positive selection after correction for multiple hypothesis testing: *DNMT3A* (q= 2.7×10^−11^), *ZNF318* (q= 1.2×10^−6^) and *HIST2H3B* (q= 0.086); (**Fig. 5c**). *DNMT3A* is a well-known myeloid cancer gene that carried 23 mutations in our dataset, of which 13 were in expanded clades. We could time the occurrence of these 13 mutations from the phylogenetic trees to a range of ages: 2 were acquired before the age of 10, 3 before the age of 20, and the remainder before the age of 40. *ZNF318*, a zinc finger transcription factor previously identified in two studies of age-related clonal haematopoiesis^22,55^, carried predominantly truncating and nonsense mutations. *HIST2H3D* showed a cluster of missense mutations in the N terminal region, neighbouring amino acids whose post-translational modifications play a critical role in transcriptional regulation^56^ (**Table S7**). Interestingly, screening 534 AML genomes^57–59^ for variants in *ZNF318* and *HIST2H3D* identified only one possible oncogenic mutation in *ZNF318* (**Table S8**), showing that while these variants are under selection in HSC/MPPs they do not necessarily contribute to the development of malignancy. We found 7 likely driver mutations in *TET2*, and 1 each in *ASXL1, CBL* and 5 other known myeloid cancer genes.

The genome-wide estimate of dN/dS was significantly elevated at 1.06 (CI_95%_=1.03-1.09; **Fig. 5d**), equating to 1 in 18 (1/34-1/12) non-synonymous coding mutations in the dataset being drivers (defined as any mutation under positive selection). Estimated dN/dS ratios were virtually identical in young and old individuals (1.06 and 1.05 respectively), suggesting that the *fraction* of mutations under positive selection does not change with age. Given that the overall mutation rate is constant across the lifespan, driver mutations therefore enter the HSC pool at a constant rate throughout life, and the expected *number* of driver mutations per cell increases linearly with age.

Converting dN/dS ratios to estimated numbers of driver mutations revealed that each adult studied typically had >100 driver mutations among the colonies sequenced (**Extended Fig. 6d**). These numbers are considerably higher than the number of non-synonymous mutations we identified in known myeloid cancer genes. This implies that mutations under positive selection in normal HSCs affect a much wider set of genes than usually assessed, as suggested by other studies^23^, and corroborates our observation that many clonal expansions in the elderly occur in the absence of mutations in known myeloid cancer genes.

We also noted that loss of the Y chromosome was a frequent occurrence in the older males in our cohort. Loss-of-Y has been well described in bulk blood samples from ageing men, correlating with all-cause mortality^35^, but whether this plays a functional role or is merely a marker of general chromosomal instability is unclear. Our clone-specific data enabled us to assess its distribution across the lineage tree. Strikingly, in many of our older males, the Y chromosome was lost multiple times in independent clones – the oldest male in our study, aged 81, had independently lost the Y chromosome at least 62 times across the phylogeny. Furthermore, loss-of-Y was significantly correlated with clonal expansion (p<0.001; **Fig. 5e**) – several of the expanded clades in our dataset had loss-of-Y, often in the absence of known point mutation drivers, even in the younger males (**Fig. 2-3**). This suggests that loss-of-Y is under positive selection in HSCs, corroborating mouse models showing *KDM6C* on the Y chromosome suppresses leukaemogenesis^60^.

### Modelling positive selection in HSCs

The genetic analysis suggests that positive selection is pervasive within HSCs from both young and old, but whether positive selection can completely explain the observed phylogenies remains unclear. To address this, we used Approximate Bayesian Computation (ABC)^52,53^ modelling to infer HSC dynamics across the lifespan, with three aims: (1) to assess whether clone-specific positive selection can explain apparently abrupt loss in clonal diversity after the age of 70; (2) to estimate the rate at which driver mutations enter the HSC pool across life; and (3) to estimate the distribution of fitness effects, defined as the excess average growth rate per year of a clone with the driver over that of wild-type HSCs (*s*=0 indicates neutrality; **Extended Fig. 8a-d; Supplementary Methods**). Initial simulations showed that mutations with a fitness effect *s*<5% are unlikely to expand to >1% of HSCs over a human lifespan (**Extended Fig. 8d**), so were not further considered.

The modelling showed that our observed trees could be closely emulated (**Extended Fig. 9a**), and that they carry sufficient information to provide meaningful refinement of the credible distributions for occurrence rate and fitness effects of driver mutations (**Fig. 5f, Extended Fig. 9b, Supplementary Fig. 12**). Observed phylogenies are most consistent with driver mutations entering the HSC compartment at a rate of ≥2.0×10^−3^/HSC/year. This rate is broadly comparable with the estimate obtained from the genetic analysis. Non-synonymous mutations accumulated in HSC/MPPs at a rate of 0.12/HSC/year (CI_95%_=0.11-0.13), with dN/dS estimates suggesting that 1/34 to 1/12 non-synonymous mutations were drivers. This computes to a driver rate of 3.6-10.0×10^−3^/HSC/year estimated from the genetic analysis, an estimate which would include drivers with *s*<5% present in sequenced colonies.

For fitness effects, the distribution most consistent with the data had a preponderance of moderate-effect drivers, with *s* in the range 5-10%, but a heavy tail of rare drivers conferring greater selective advantage (*s*>10%). For the 46 largest observed clones, we could directly estimate their fitness effects from the patterns of coalescences within their clade, resulting in estimates of *s* in the range of 10-30% (**Fig. 5g, Extended Fig. 9c**). This is consistent with the heavy tail of the credible distribution from simulation models, and shows that clones without known drivers can evolve comparable selective advantages to classic driver mutations.

The ABC modelling generates simulations of HSC clonal dynamics across a lifetime, which enables us to track how phylogenetic trees for a given avatar would appear when sampled at different ages using the study design deployed here. In these avatars, we found that there was typically a sharp change towards oligoclonal haematopoiesis in the tree’s appearance after the age of 60 (**Extended Fig. 10**), thus demonstrating that the observed trees can indeed be entirely accounted for by steady accumulation of positively selected mutations within the HSC compartment.

## Discussion

Our data provide a pleasingly parsimonious model of how lifelong, gradual accumulation of molecular damage can result in an abrupt, qualitative change in tissue composition towards the end of life. HSCs accumulate somatic mutations at a steady rate throughout life with minimal cell-to-cell and person-to-person variation. We find the fraction of somatic mutations under positive selection is constant across the lifespan, meaning that driver mutations also enter the HSC compartment at a steady, linear rate across the lifespan. These drivers mostly confer moderate fitness benefits, so the clonal expansions they trigger, while exponential, only gather momentum slowly, over decades. Simulations under this simplistic model of haematopoiesis – constant HSC population size, constant rate of driver mutations, slow but inexorable exponential expansion – entirely recapitulate the dramatic and apparently inevitable shift towards oligoclonality observed after the age of 70. Human haematopoiesis is certainly more complex than this model, with its various compartments of stem and progenitor cells, lineage biases, epigenetic change and interconnected microenvironment. However, our data clearly demonstrate that complex age-related dynamics can emerge from the convolution of predictable, lifelong processes.

Our findings provide a framework to understand how known lifestyle and disease risk factors causally contribute to the development of clonal haematopoiesis and myeloid malignancy. Factors that increase HSC turnover (such as chronic inflammation^61^, smoking^62^, infection^63^, autoimmune disease^64^) or strengthen the selective advantage of clones with driver mutations (such as chemotherapy^42^) have been shown to induce earlier expansion of clones with driver mutations in known myeloid cancer genes. One prediction of our model is that these factors should also bring forward the time of transition to oligoclonal haematopoiesis.

Other testable corollaries emerge. First, the large population size of HSCs and steady accumulation of driver mutations suggest that loss of clonal diversity in aged haematopoiesis will be universal in the elderly. Second, although inevitable, which driver mutations accrue, and when in life they occur, will strongly shape the size and behaviour of the few clones that come to dominate haematopoiesis, thus generating considerable person-to-person variation. We already know this variation predicts future risk of blood cancer^65,66^; it will be fascinating to determine whether it also predicts other phenotypes of ageing haematopoiesis such as anaemia and reduced resilience to chemotherapy and infection^67^. Third, if the various driver mutations share common downstream phenotypes, we may find that oligoclonality underpins emergent features of ageing blood, such as recurrent DNA methylation patterns^12^, biases in lineage commitment^68^, and global reduction in differentiation capacity^69,70^.

Importantly, oligoclonality also occurs in the elderly haematopoietic systems of other mammalian species, including mice^70^ and macaques^71^. In addition, virtually all human organ systems studied to date, including skin^72,73^, bronchus^74^, endometrium^75,76^ and oesophagus^77,78^, show a similar age-related increase in the density of clones with driver mutations. With such ubiquity of driver mutations, selected purely for their competitive advantage within the stem cell compartment, and with the wholesale rewiring of cellular pathways they induce, it is feasible that they may contribute to phenotypes of human ageing beyond the risk of cancer.

## Supporting information

Mathematical basis for Approximate Bayesian Computation

## ACKNOWLEDGEMENTS

This work was supported by the WBH Foundation. Investigators at the Sanger Institute are supported by a core grant from the Wellcome Trust. P.J.C. was a Wellcome Trust Senior Clinical Fellow (WT088340MA) until 2020. N.F.∅. is the recipient of a Danish Lundbeck Fellowship (2016-17) and M.S.S. is the recipient of a BBSRC CASE Industrial PhD Studentship. Work in the D.G.K. laboratory is supported by a Bloodwise Bennett Fellowship (15008), a European Research Council Starting Grant (ERC-2016-STG–715371) and a European Hematology Association Non-Clinical Advanced Research Fellowship. Work in the A.R.G. laboratory is supported by the Wellcome Trust, Bloodwise, Cancer Research UK, the Kay Kendall Leukaemia Fund, and the Leukemia and Lymphoma Society of America. Work in E.L. laboratory is supported by a Wellcome Trust Sir Henry Dale Fellowship, BBSRC and a European Haematology Association Non-Clinical Advanced Research Fellowship. The E.L. and A.R.G. laboratories are supported by a core support grant from the Wellcome Trust and Medical Research Council to the Cambridge Stem Cell Institute. K.M. is supported by the Chan-Zuckerberg Initiative. K.C. is supported by a Wellcome Investigator award (210755/Z/18/Z). We acknowledge further assistance from the National Institute for Health Research Cambridge Biomedical Research Centre and the Cambridge Experimental Cancer Medicine Centre. We are grateful to the donors, donor families and the Cambridge Biorepository for Translational Medicine for the gift of their tissue.

## EXTENDED FIGURE LEGENDS

**Extended Data Fig. 1**| **Flow-sorting strategy for single HSC/MPP and HPC cells. a**, Sorting of single human HSC/MPP and HPCs from cord blood, peripheral blood and bone marrow. Cells were stained with the panel of antibodies in **Table S1** then single HSC/MPP or HPCs were index sorted according to the strategy depicted into individual wells of 96 well plates. **b**, Colony forming efficiency per individual of all single HSPCs sorted.

**Extended Data Fig. 2**|**Quality assurance of mutation calls. a**, Histogram of VAFs for a typical sample in the dataset, showing a tight distribution around 50%, as expected for an uncontaminated clonal sample derived from a single cell. The variants with VAFs < 0.2 represent *in vitro* acquired mutations and sequencing artefacts and were removed using a VAF-based filtering strategy with a cut off of 0.2 (red line). **b**, VAF distribution of variants after filtering steps had been applied. The red line shows the peak VAF and the dashed grey line shows the threshold peak VAF for excluding samples as being non-clonal / contaminated. **c**, Histogram of VAFs for a colony that was seeded by 2 cells showing a median VAF around 25%. Colonies showing evidence of non-clonality in this way were excluded from downstream analysis using a peak VAF cut off of 0.4. **d**, Left-hand plot shows the relationship between raw mutation counts per colony for one individual post filtering and sequencing depth. The black line depicts an asymptotic regression line fitted to the raw data. Right-hand plot shows the adjusted mutation burdens per colony after asymptotic regression correction. **e**, Trinucleotide context mutation spectra of private (top plot) and shared variants (bottom plot) for one individual. The spectra are extremely similar, showing the variant filtering strategy used is robust and prevents excess artefacts in the private variant set. **f**, Trinucleotide mutation spectrums for each individual created from all variants post filtering. The results are consistent between the two cord blood donors and all the adult donors.

**Extended Data Fig. 3**|**Approach to phylogeny construction. a**, Raw phylogeny for KX003 (81-year male) derived directly from *MPBoot*. The input to *MPBoot* is a genotype matrix of all variant calls shared by more than 1 colony from an individual. **b**, Phylogeny with edge lengths proportional to the number of mutations assigned to the branch using original count data and the *tree_mut* package. **c**, Phylogeny with raw mutation count branch lengths adjusted for sequencing depth of the sample using sensitivity for germline variant calling. **d**, Phylogeny with adjusted branch lengths converted to ultrametric form (equal branch lengths). One axis shows mutation number, the other axis shows the equivalent estimated age in years, which is possible due to the linear accumulation of mutations in HSPCs with time. All tips end at age 81, the age at the time of sampling.

**Extended Data Fig. 4**|**Mutational burden. a**, Regression of number of single nucleotide variants (SNVs) in HSCs (red line) compared to HPCs (blue line). Grey shading indicates the 95% CI. The estimated difference in burden, together with the t-value is above the plot. The t-value of 1.54 demonstrates non-significance of the difference. **b**, Phylogenies depicted for the individuals with clonally expanded structural variants (SVs). The bar at the bottom highlights cells with one of the three classes of structural variant. The exact variant breakpoints can be found in **Table S3. c**, Plot showing the percentage of HSC/MPP cells that have outlying telomere lengths per individual. Outliers were identified using the Interquartile Range criterion. There were no outliers with shorter than expected telomeres in any individuals, such that this data only reflects the percentage of cells with longer than expected telomeres. The blue line shows a regression of percentage outlying telomere lengths with age. This shows a significant negative correlation (t-value and p-value shown).

**Extended Data Fig. 5**|**Interpretation of young adult HSPC phylogenies. a**, Trajectories of *Nτ* used as input to *rsimpop* for the simulations to create phylogenies in b. Note the Y axis depicting *Nτ* is on a log scale. **b**, Phylogenies created by randomly sampling 380 cells from the final full simulated population of between 100,000 cells (Phylogeny 1) and 1,000,000 cells (Phylogeny 4). Phylogenies 1 to 3 are derived from simulations of the HSC population in a 30-year-old, while phylogeny 4 is derived from a simulation of the HSC population in an 80-year-old. Each simulation has an initial *Nτ* of 100. In all cases *Nτ* is the same as the population size (*N*) as the generation time (*τ*) in all simulations is fixed at 1. The blue boxes indicate the period of time in which the population size is increased. The *phylodyn* trajectories to the right of each simulated phylogeny use the pattern of coalescent events to recover the input trajectories for *Nτ*. The blue line marks the time of change in *Nτ*. In all cases the initial part of the trajectory is able to correctly estimate *Nτ* at 100,000. However, in Phylogeny 3 where there is a complete absence of coalescent events once the population size is increased, *phylodyn* loses resolution and wildly overestimates the value of *Nτ*. **c**, Real trees with red boxes highlighting the last 10-20 years prior to sampling, where the relative number of coalescent events is decreased (meaning the estimated *Nτ* is larger).

**Extended Data Fig. 6**|**Interpretation of elderly adult HSPC phylogenies. a**, Phylogenies created by randomly sampling 380 cells from the final full simulated population. As with the previous simulations, *Nτ* ∼ population size because the time between symmetric self-renewal divisions is set at 1 year. In both simulated phyologenies the final population is 100,000 cells in size, but Phylogeny 2 has been created from a population that underwent a bottleneck in size to and *Nτ* of 10,000 cells between the ages of 35 and 50. This period of time over which the population size was reduced can be visualised in the blue box which highlights the increased density of coalescent events in this time block. The phylodyn trajectories are able to accurately recover information on changes in *Nτ* of the HSC population over time. **b**, Real HSC/MPP phylogeny for KX003 (81-year-male) with PB HSC/MPP terminal branches coloured red (BM HSC/MPP branches remain black). The CF of the largest clade is shown for PB and BM cells. **c**, Real HSPC phylogeny for KX004 (77-year-female) with BM HPC terminal branches coloured blue (BM HSC/MPP branches remain black). The CF of two clades is shown for HSC/MPPs and progenitor cells. **d**, Estimated number of driver mutations in the different datasets. The boxes show the estimate with whiskers showing the 95% CI. The numbers to the left give the numeric values for the estimates with 95%CI in brackets. ‘n’ is the number of cells included in each dataset.

**Extended Data Fig. 7**|**Modelling HSC populations incorporating only changes in N*τ*, without positive selection. a**, Overview of modelling approach used to estimate *Nτ* alone in the young adult individuals and to investigate whether changes in *Nτ* could explain the observed clade size distribution in the elderly adult individuals. These simulations were run using a neutral model (that is, no acquisition of driver mutations), with *Nτ* being the only parameter to change over time. For the young adult individuals *Nτ* was estimated for two time-blocks (time before and after population increase due to ST-HSC/MPP contribution). For the elderly adult individuals *Nτ* was estimated for three time-blocks as the *phylodyn* plots predicted a population ‘bottleneck’ (**Supplementary Fig. 11**) was the most parsimonious way to recreate the observed change in coalescence density over life. **b**, Plots showing the posterior predictive distribution of the difference between the simulated chi-squared discrepancy and the observed chi-squared discrepancy, for each donor individual under a neutral model incorporating change in population size. For each donor, the posterior predictive distribution of the difference between predictive (simulated) and observed chi-squared discrepancy is represented as a histogram based on a Monte Carlo sample of 1,000,000 simulated phylogenies, drawn from the posterior predictive distribution. The proportion of simulated phylogenies which lie to the right of zero (red line) is a Monte Carlo estimate of the posterior predictive p-value (the probability that the predictive chi-squared discrepancy exceeds the observed chi-squared discrepancy under the neutral model). In the case of the four young adult individuals, the proportion of simulated phylogenies which lie to the right of 0 (red line) is close to 0.5, indicating that the simple neutral models (incorporating changes in Nt over life) predict trees that have similar clade size distributions to our observed trees. In contrast, for the four elderly adults, the proportion of simulated phylogenies which lie to the right of 0 is very small (Less than 0.05), demonstrating that the neutral models are, on their own, unable to recreate trees with similar clade size distributions to those observed.

**Extended Data Fig. 8**| **Modelling of HSC populations incorporating positive selection. a**, Overview of modelling approach used to estimate the shape and rate of the gamma distribution of selection coefficients from which ‘driver mutations’ are drawn, and the number of driver mutations drawn from this distribution (using a selection coefficient threshold of > 0.05) that are entering the HSC population per year. For these simulations *Nτ* was fixed at 100,000 and therefore only summary statistics for the first 3 timepoints were used to assess how well a given simulation for an individual resembled the observed tree. **b**, Plot showing maximum posterior density estimates of the rate and shape parameters of the gamma distribution for selection coefficients (pink line) obtained using Approximate Bayesian computation. Blue/green lines show how altering the rate and shape parameters affect the gamma distribution. **c**, Plot showing how changing the shape of the gamma distribution of selection coefficients (each line has a different shape) alters the probability of a driver gene fixing in the population. Reducing the shape below 0.1 does not affect the probability of driver gene fixation and therefore was the lower limit of the shape prior. **d**, Plot showing how the probability of detecting a clone with CF 2.5% changes over time for different selection coefficients. There is only a probability of 0.1 of being able to identify a driver mutation with a selection coefficient of 0.05 that entered the population at birth. We therefore used a lower threshold of 0.05 for the driver mutation selection coefficients

**Extended Data Fig. 9**|**Driver modelling results and expanded clade annotation. a**, Plots showing the posterior predictive distribution of the difference between the predictive (simulated) chi-squared discrepancy and the observed chi-squared discrepancy, for each donor individual under the simple positive selection model. For the definition of the chi-squared discrepancy, and details of how the posterior predictive p-values are estimated, see Supplementary information “Posterior predictive model checking (PPC) methods which can be applied to Approximate Bayesian Computations (ABC)”, Sections 1, 2 and 5. In these plots, the chi-squared discrepancy is computed from summary statistics evaluated at the first 3 (out of 4 equally spaced) timepoints on the phylogeny obtained from the specified donor (**Extended Fig. 8**). For each donor, the posterior predictive distribution of the difference between predictive (simulated) and observed chi-squared discrepancy is represented as a histogram based on a Monte Carlo sample of at least 100,000 simulated phylogenies, drawn from the posterior predictive distribution. The proportion of simulated phylogenies which lie to the right of zero (red line) is a Monte Carlo estimate of the posterior predictive p-value (the probability that the predictive chi-squared discrepancy exceeds the observed chi-squared discrepancy under the positive selection model). Those p-values written in grey text are based on chi-squared discrepancies computed from summary statistics evaluated at the first 2 (out of 4 equally spaced) timepoints. Notice that all these p-values are all above the 0.05 threshold, indicating that observed phylogenies (up to the second time point) are compatible with the simple positive selection model. Those p-values written in blue text are based on chi-squared discrepancies computed from summary statistics evaluated at the first 3 (out of 4 equally spaced) timepoints. Notice that all but two observed phylogenies (up to the third time point) are compatible with this positive selection model. These p-values indicate that, once the third time point is included, the phylogenies of two of the younger individuals (38 year-old and 48 year-old) are no longer compatible with the positive selection model. Notice that these two donors also exhibit the most striking increase in population size from the middle part of the population trajectory onwards (**Fig. 4a**). When all four timepoints are included, the phylogenies of 5 out of 8 donors have become incompatible with the positive selection model (data not shown). Only the phylogenies from the donors of ages 77, 76 and 29, remain compatible with the positive selection model. This suggests that the current positive selection model does not adequately account for the population processes towards the time of sampling. **b**, Posterior distribution of number of ‘driver’ mutations with s>5% entering HSC population of 100,000 cells per year. Black line shows peak estimate. **c**, Phylogenies of the four adults aged > 75 labelled with driver mutations and clade ID annotations as used in Fig. 5g.

**Extended Data Fig. 10**|**Positive selection over life**. Four consecutively simulated phylogenies of 380 cells sampled from a population of 100,000 cells that has been maintained at a constant *Nτ* over life, with incorporation of positively selected ‘driver mutations’. The driver mutations have a fitness effect > 5% (drawn from a gamma distribution with shape = 0.73 and rate = 33) and enter the population at a rate of 200 per year. These are the maximum posterior density estimates of the rate and shape parameters obtained using the ABC method. The inclusion of these driver mutations is able to recapitulate a similar clade size distribution to that observed in the real HSPC phylogenies of the observed individuals across the whole age range. However, including driver mutations does not fully recapitulate the observed lack of coalescent events in the last 10-15 years of life, showing that an increase in *Nτ* over this time is also required to fully recreate the patterns of coalescences in the real phylogenies. Driver mutations are marked with a symbol and their descendent clades are coloured. In all cases *Nτ* is the same as the population size (*N*) as the generation time (*τ*) in all simulations is fixed at 1 year. The symbols / colours are not consistent for driver mutations between plots. The largest clades are therefore coloured in a consistent way beneath the plots to show how their size changes over time. Between the ages of 60 and 80 almost all clades expand. However, between 80 and 100 variable patterns of growth are observed, with some clades staying stable in size and others getting smaller. In all cases there are 1 or more ‘fittest’ clones that continue expanding.

