## Extended Figures 1-10 for "Clonal dynamics of haematopoiesis across the human lifespan"

a

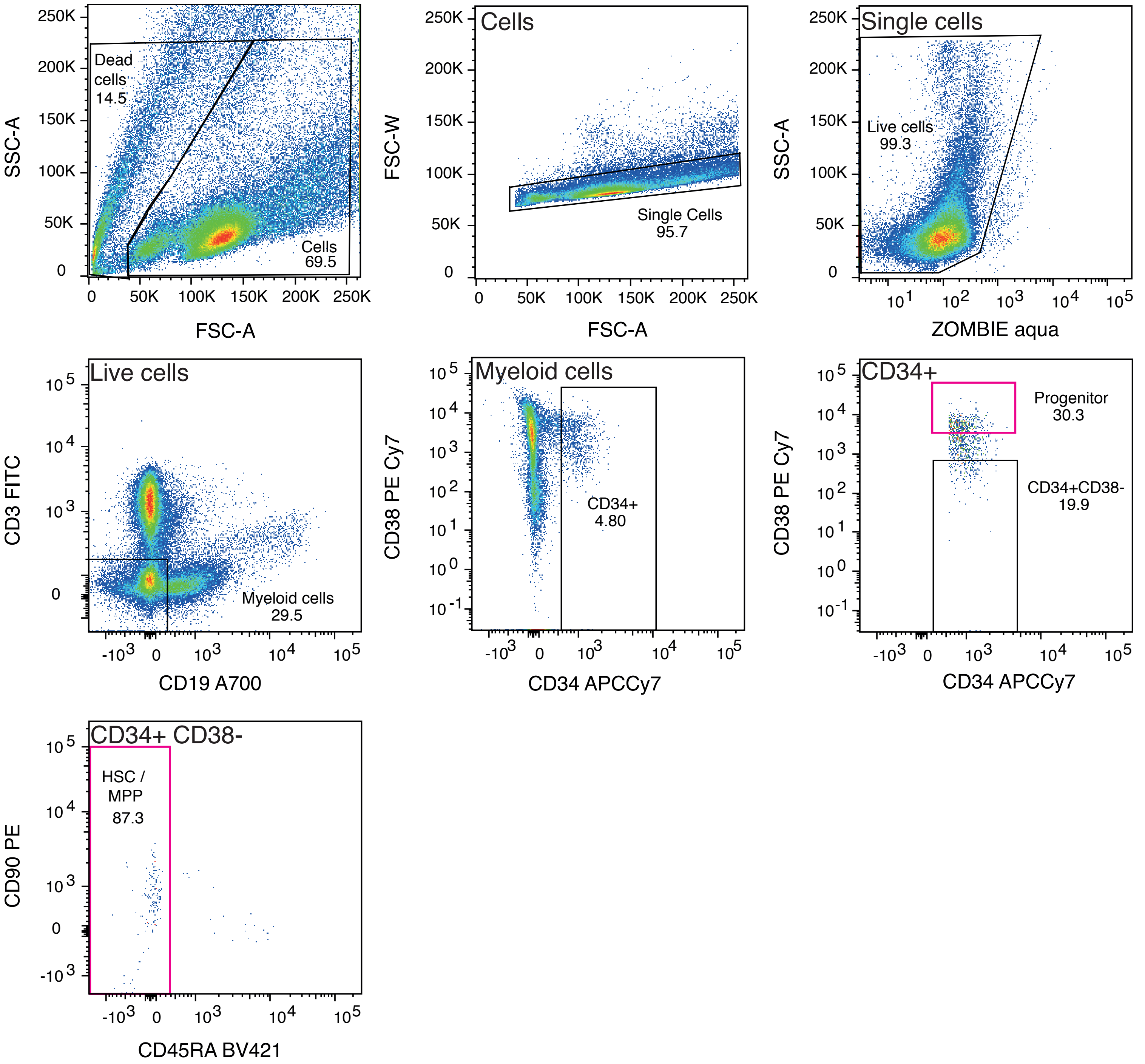


b

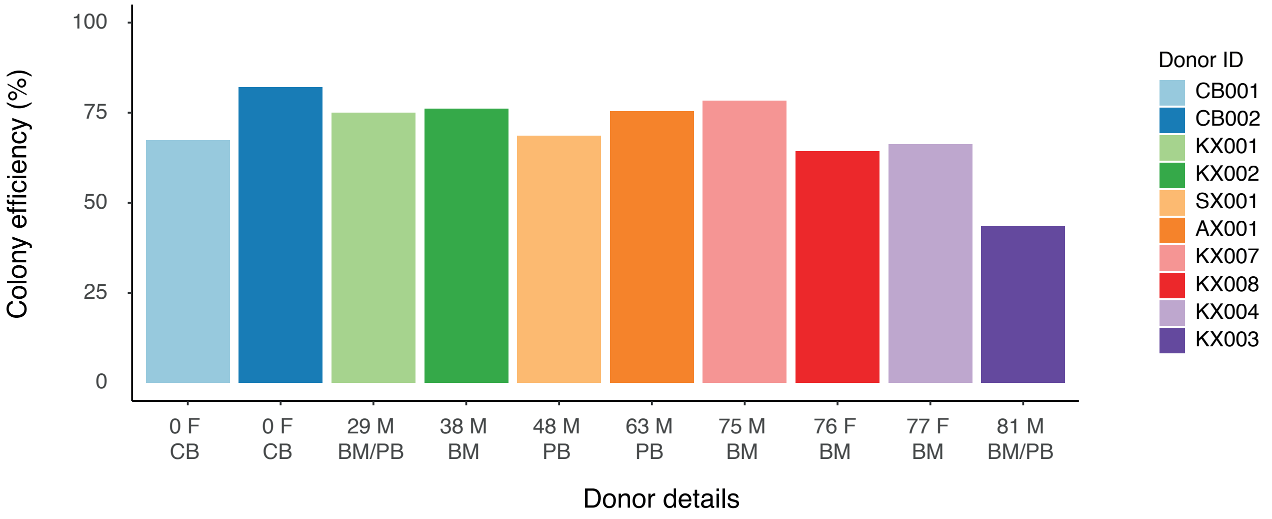


Extended Figure 1

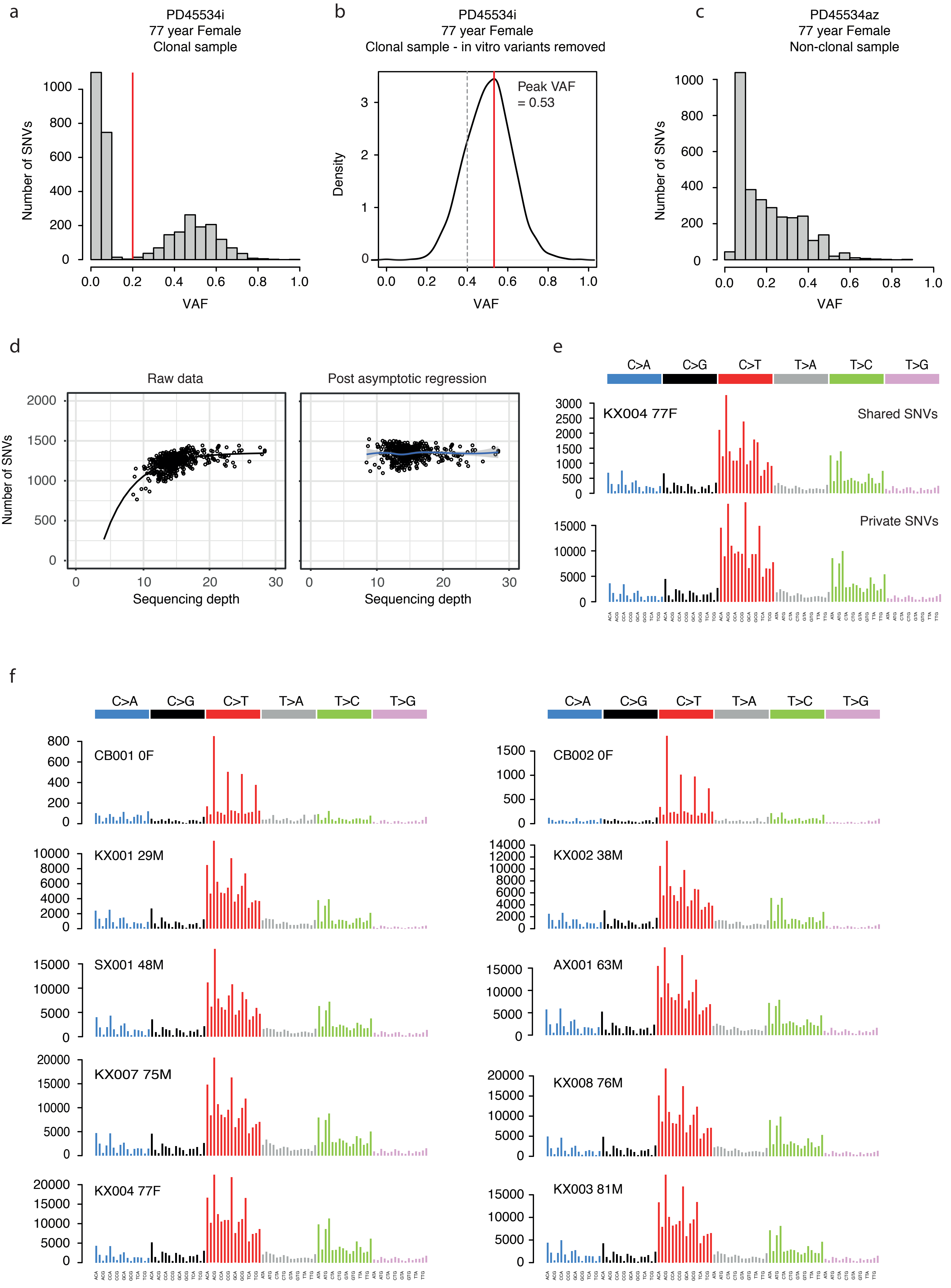


Extended Figure 2

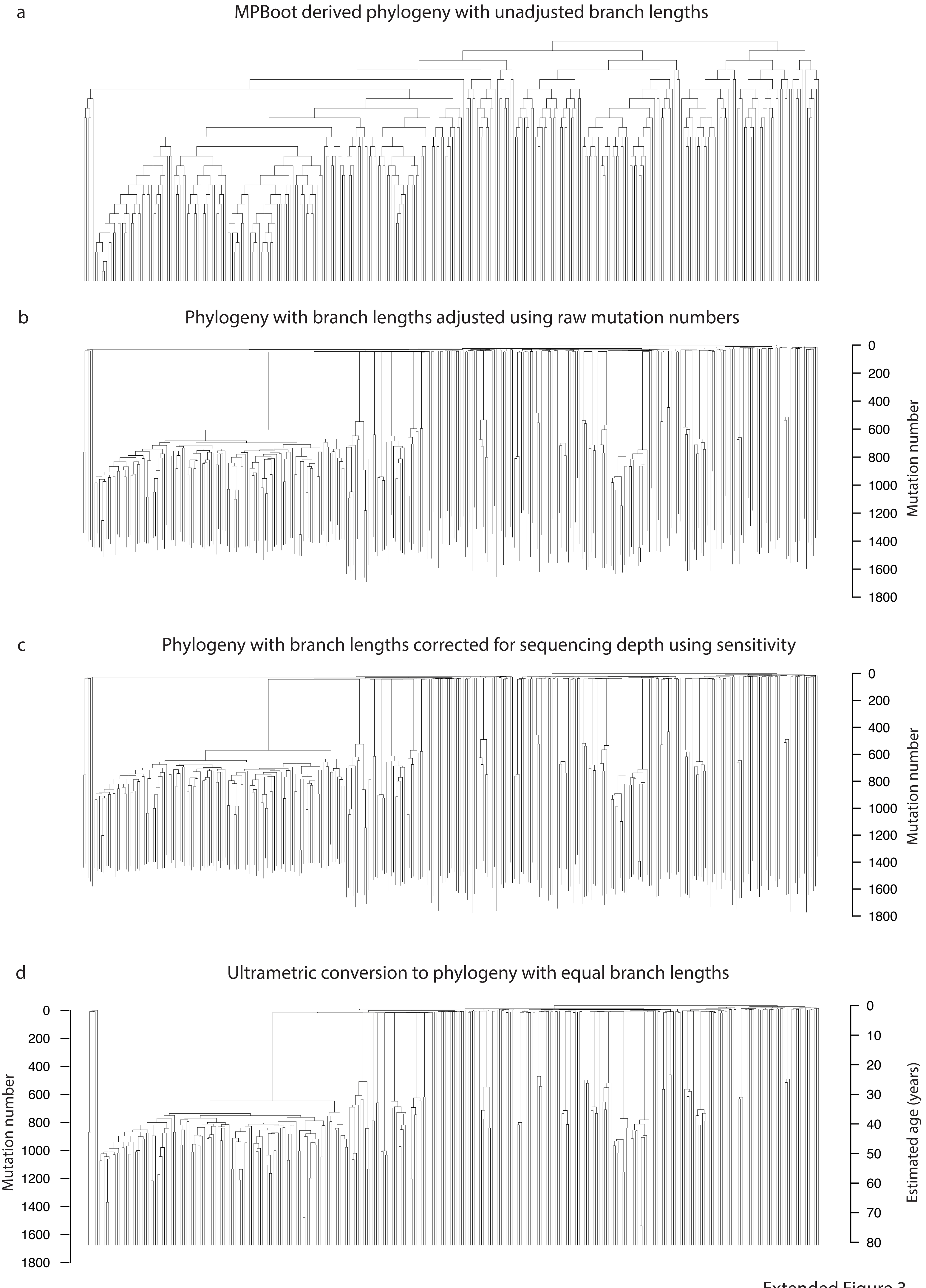


Extended Figure 3

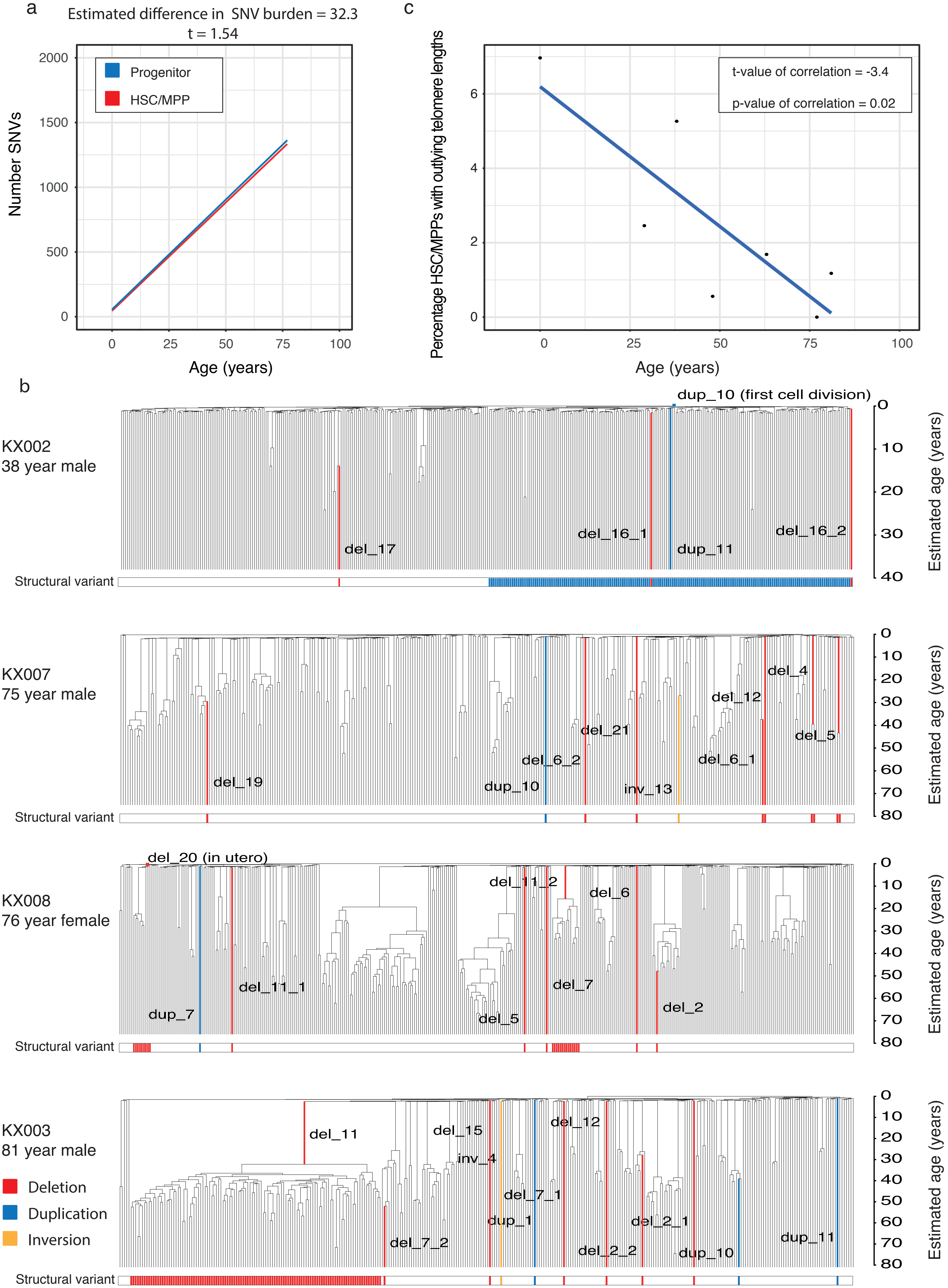


Extended Figure 4

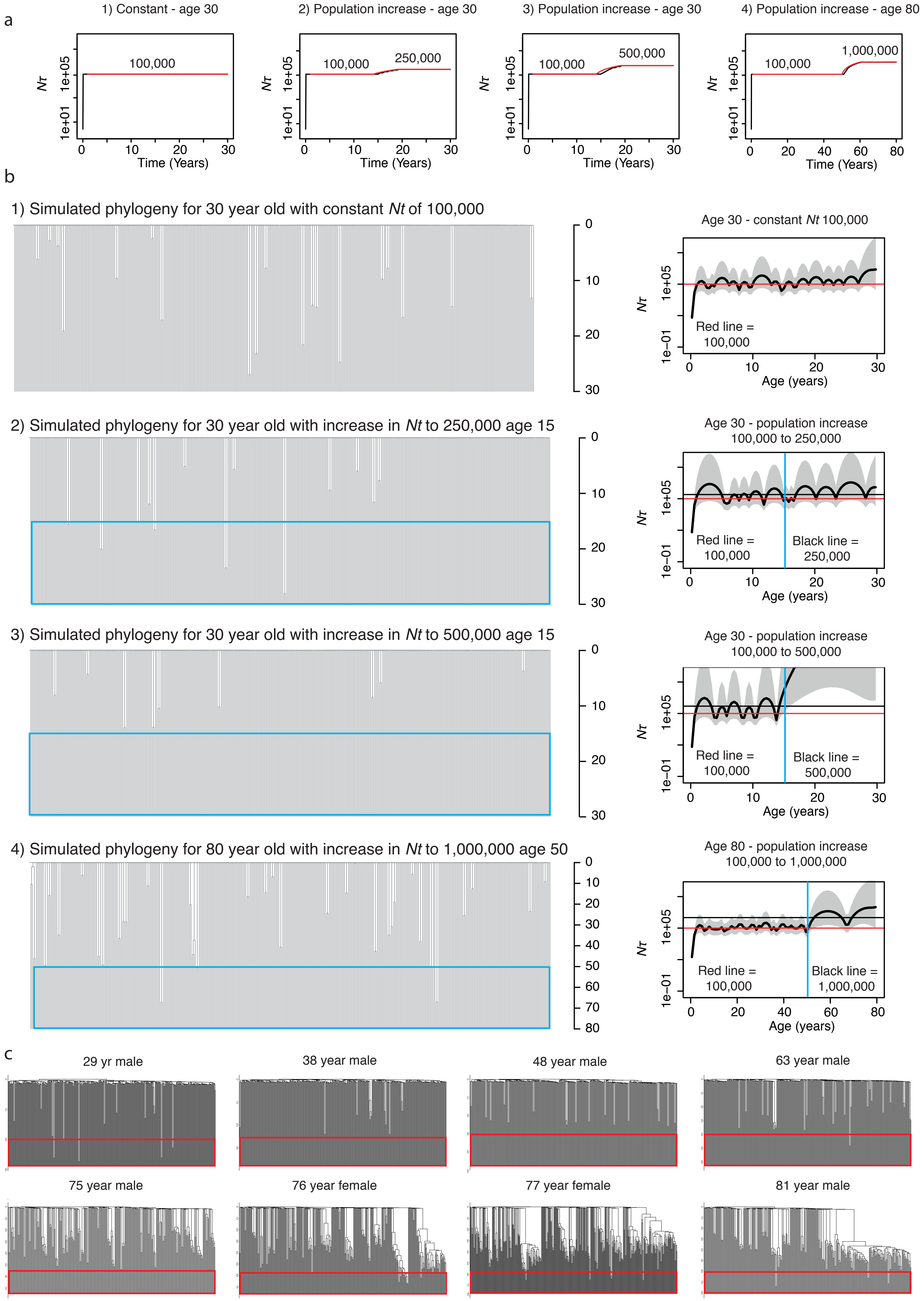


Extended Figure 5

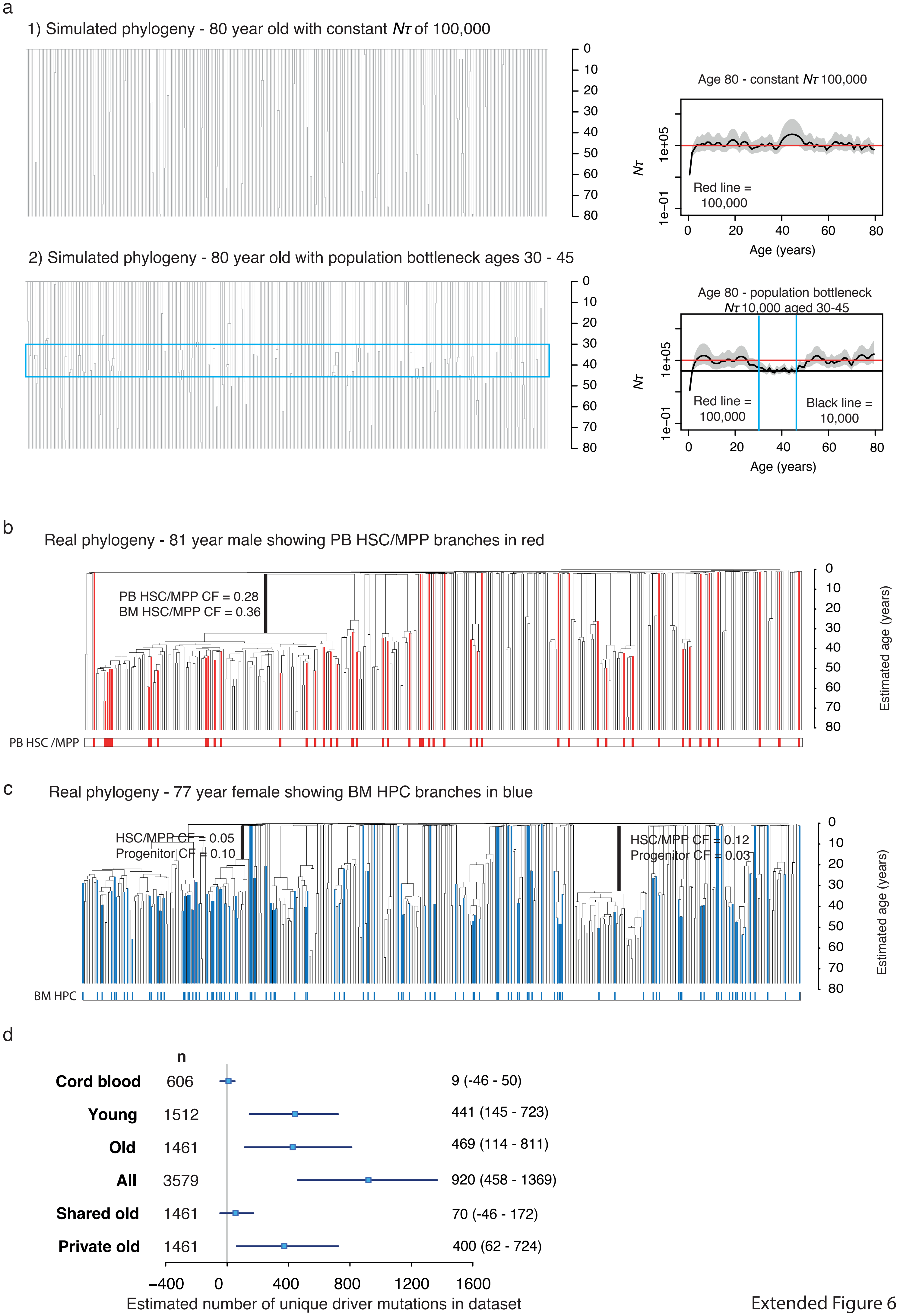


Extended Figure 6

a

### Bayesian inference

#### Model: Neutral model allowing varying HSC population size over life

- time between symmetric self renewing cell divisions fixed at a point estimate of 2 per year ($N/2 = N\tau$)

**Prior**

Independent prior densities on three parameters:

1. N in 1st 2-3 decades of life (all donors)
   - uniform prior on interval [a,b],
   a = 1000 b = 250,000, 500,000, 750,000
2. Midlife fold-change in *N* (elderly donors only)
   - uniform prior on interval [ 0.01,1.2]
3. Latelife fold-change in *N* (all donors)
   - uniform prior on interval [0.5 to 8]

**Summary statistics for ABC**

1. Time-weighted mean number of lineages
(calculated at 3 time-points: 0.25, 0.5, 0.75)

#### Approximate Bayesian Computation using ABC rejection method

Donor-specific ABC performed separately on data from each donor
Separate Monte Carlo samples generated from the (approximate) donor-specific posterior distribution (one for each donor).
For each donor-specific ABC:
- 100,000 parameter vectors sampled from prior
- From each parameter vector a simulated data set (phylogeny) is generated using (neutral) rsimpop
- From each simulated data set (phylogeny) a vector of summary statistics is computed
- Simulations ranked w.r.t. Euclidean distance between simulated and observed vectors of summary statistics
- The 1000 simulations with shortest Euclidean distances (to the observed data) are accepted (tolerance = 1000/100,000 = 0.01)

The parameter vectors for the accepted simulations represent a sample from an approximate posterior distribution.

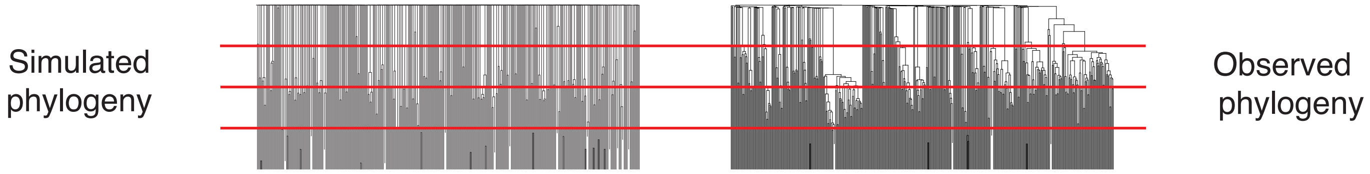


#### Posterior predictive checks (PPC)

Donor-specific (posterior predictive) p-value computed. For each donor we do the following:
- We have (from the ABC output) a sample of 1000 parameter vectors from the (approximate) donor-specific posterior distribution.
- For each parameter vector 1000 new (predicted) data sets (phylogenies) are simulated using (neutral) rsimpop;
- From each simulated data set (phylogeny) a vector of summary statistics is computed;
- From each (simulated) vector of summary statistics a (simulated) chi-squared discrepancy value is computed;

The proportion of simulations where the simulated chi-squared discrepancy value exceeds the observed chi-squared discrepancy value is an estimate of the (posterior predictive) p-value.

**Summary statistics for PPC**

(all calculated at 4 time-points: 0.2, 0.4, 0.6, 0.8)

1. Time-weighted mean number of lineages
2. Size of top 3 largest clades
3. Number of singleton samples (clade size = 1)

**Interpretation of posterior predictive p-values**

- If the (posterior predictive) p-value is close to zero, this is evidence against the proposed model as an explanation for the features of the data captured by the summary statistics.
- If the p-value is close to zero, then the observed data is an outlier compared to the data predicted under the proposed model.

b

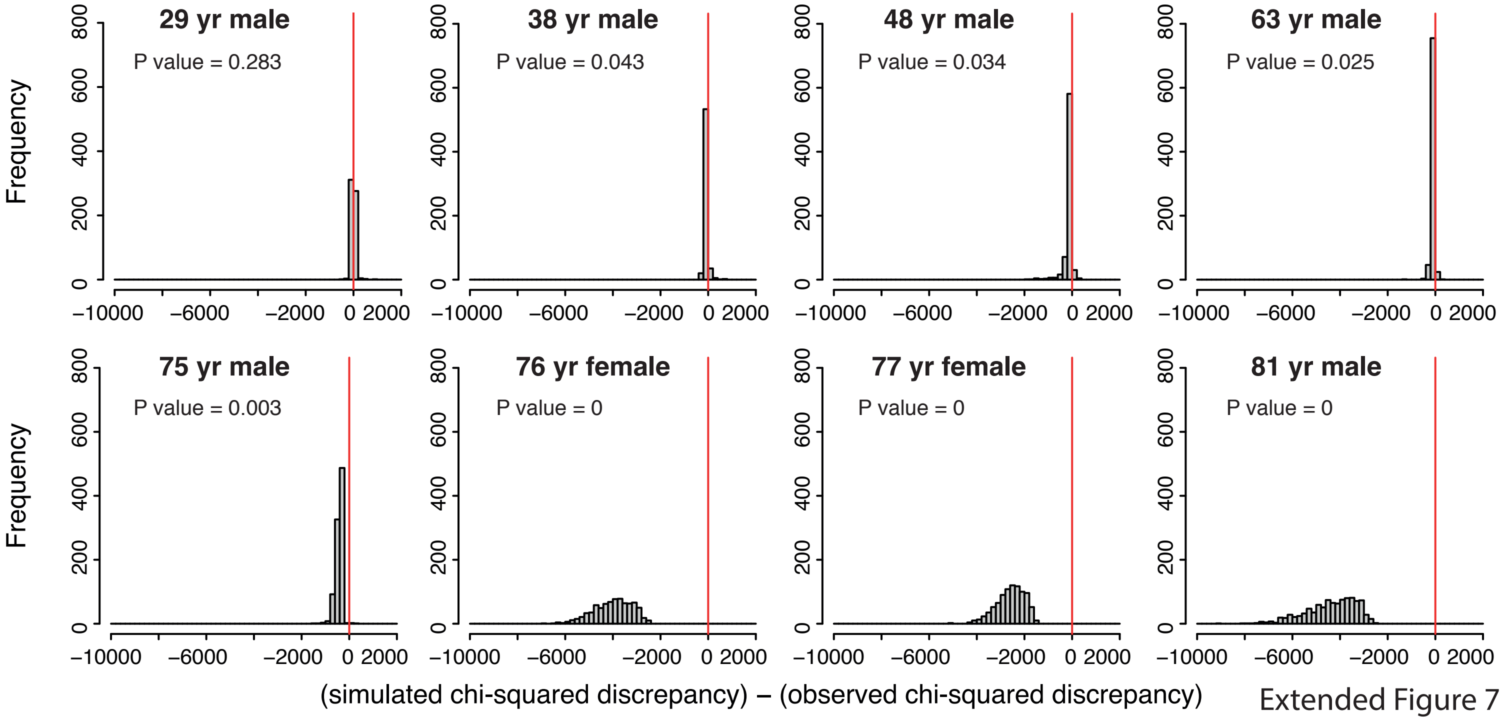


Extended Figure 7

a

### Bayesian inference

#### Model incorporating positive selection in the form of 'driver mutations'

- selection coefficient of 'driver mutation' drawn from gamma distribution;
- lower fitness threshold of selection coefficient set at 0.05 (equivalent to a fitness effect of 5% additional growth per year);
- Nτ fixed at a point estimate of 100,000 (population of N = 100,000 HSCs, dividing symmetrically once per year)

**Prior**

Independent prior densities on three parameters:

1. Shape of gamma distribution of selection coefficients
   - uniform prior on interval [0.1,2.0]
2. Rate of gamma distribution of selection coefficients
   - uniform prior on interval [5,120]
3. Number of drivers entering the population per year
   - uniform prior on interval [1,200]

**Summary statistics for ABC**

(all calculated at 3 time-points: 0.2, 0.4, 0.6 - apart from stastic 4.)

1. Time-weighted mean number of lineages
2. Size of top 3 largest clades
3. Number of singleton samples (clade size = 1)
4. Number of coalescent events

(calculated at 2 time-intervals: 0.2-0.4, 0.4-0.6)

#### Approximate Bayesian Computation using ABC regression method

Multiple-donor ABC performed on the combined data from the 4 oldest donors.
Single Monte Carlo sample from the (approximate) multiple-donor posterior distribution generated using 4 ABC regression steps.
For each ABC regression step:
- 100,000 parameter vectors sampled from 'prior'
- From each parameter vector a simulated data set (phylogeny) is generated using rsimpop (incorporating driver mutations)
- From each simulated data set (phylogeny) a vector of summary statistics is computed
- Simulations are ranked w.r.t. Euclidean distance between simulated and observed vectors of summary statistics
- The 2000 simulations with the shortest Euclidean distances (to observed data) are accepted (tolerance = 2000/100,000 = 0.02)
- For each parameter, the 2000 values (from the accepted parameter vectors) are adjusted by performing a separate ridge regression on the (re-scaled and logit-transformed) parameter values

The resulting 2000 parameter vectors of adjusted parameter values represent a sample from an approximate posterior distribution.
In the first ABC regression step, 100,000 parameter vectors are sampled from the original prior distribution.
At each subsequent ABC regression step, 100,000 parameter vectors are sampled from the posterior sample generated by the preceding ABC regression step.
The parameter vectors for the accepted simulations represent a sample from an approximate posterior distribution.
The 2000 parameter vectors of adjusted parameter values obtained from the final ABC regression step represent a sample from the (approximate) multiple-donor posterior distribution.

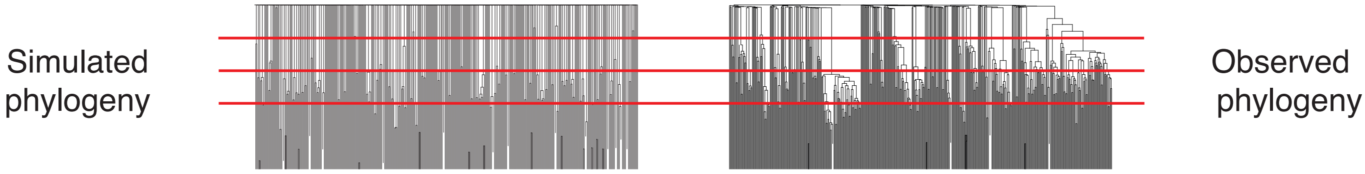


#### Posterior predictive checks (PPC)

Donor-specific (posterior predictive) p-value computed. For each donor we do the following:
- Sample a minimum of 200 parameter vectors from the sample (of 2000) from the multiple-donor posterior distribution
- For each parameter vector a min. of 500 new data sets (phylogenies) are simulated using rsimpop (incorporating driver mutations) resulting in a min. of 100,000 simulations
- From each (a min. of 100,000) simulated data set (phylogeny) a vector of summary statistics is computed
- From each (simulated) vector of summary statistics a (simulated) chi-squared discrepancy value is computed

The proportion of simulations where the simulated chi-squared discrepancy value exceeds the observed chi-squared discrepancy value is an estimate of the (posterior predictive) p-value.

**Summary statistics for PPC**

For 1st PPC: calculated at 2 time-points (0.2,0.4)
For 2nd PPC: calculated at 3 time-points (0.2, 0.4, 0.6)

1. Time-weighted mean number of lineages
2. Size of top 5 largest clades
3. Number of singleton samples (clade size = 1)
4. Number of coalescent events

**Interpretation of posterior predictive p-values**

- If the (posterior predictive) p-value is close to zero, this is evidence against the proposed model as an explanation for the features of the data captured by the summary statistics.
- If the p-value is close to zero, then the observed data is an outlier compared to the data predicted under the proposed model.

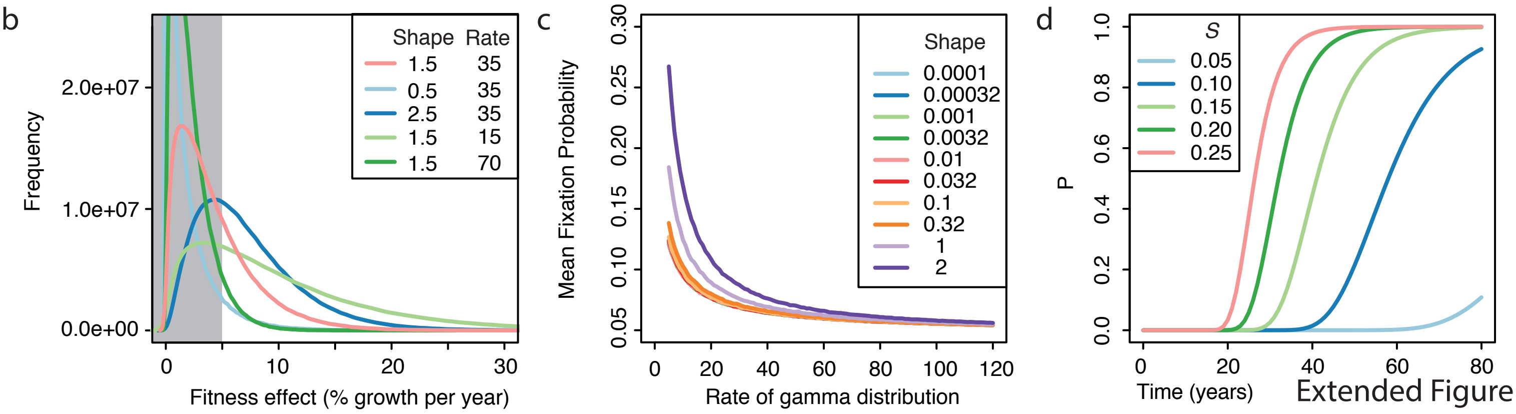


Extended Figure 8

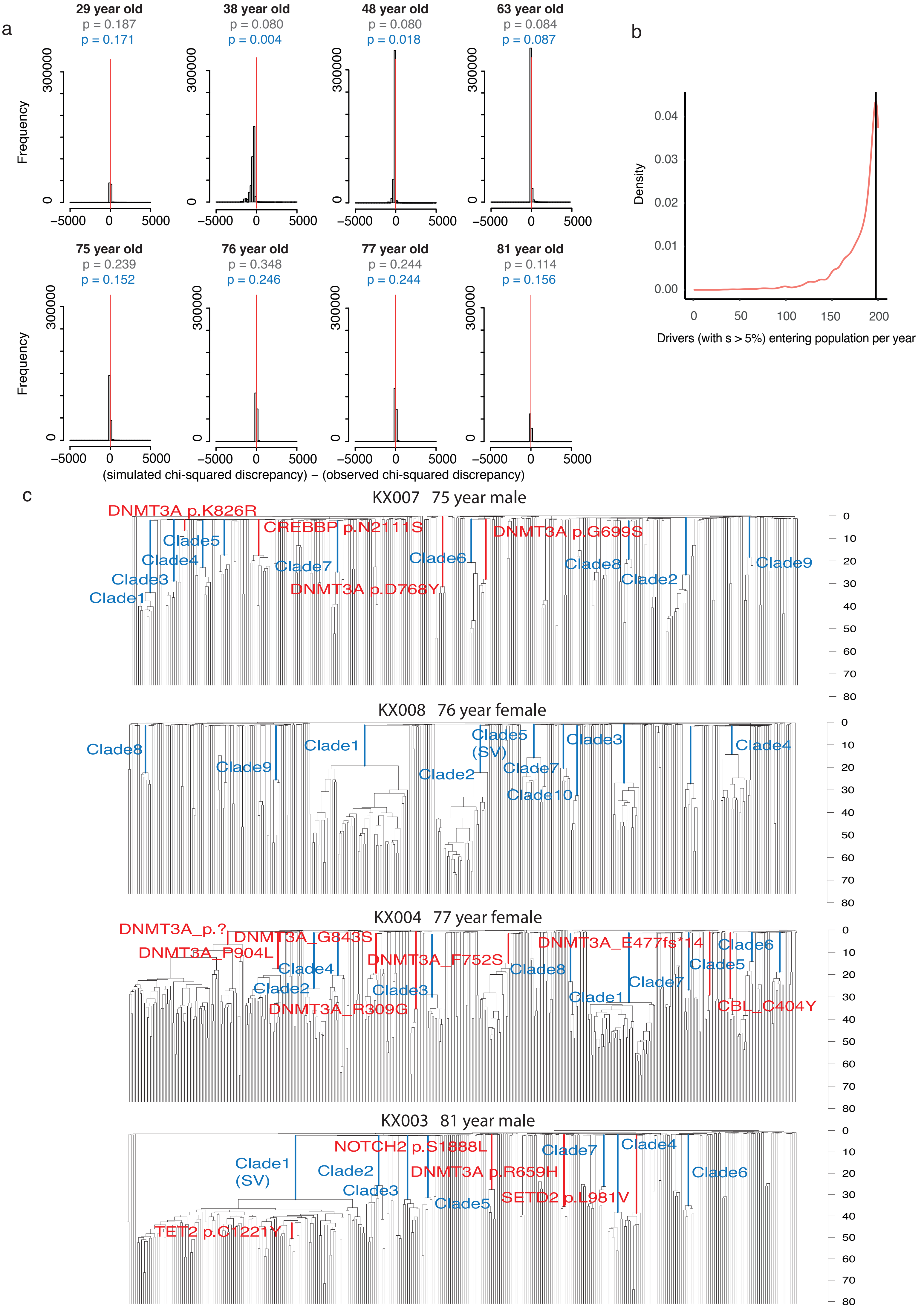


Extended Figure 9

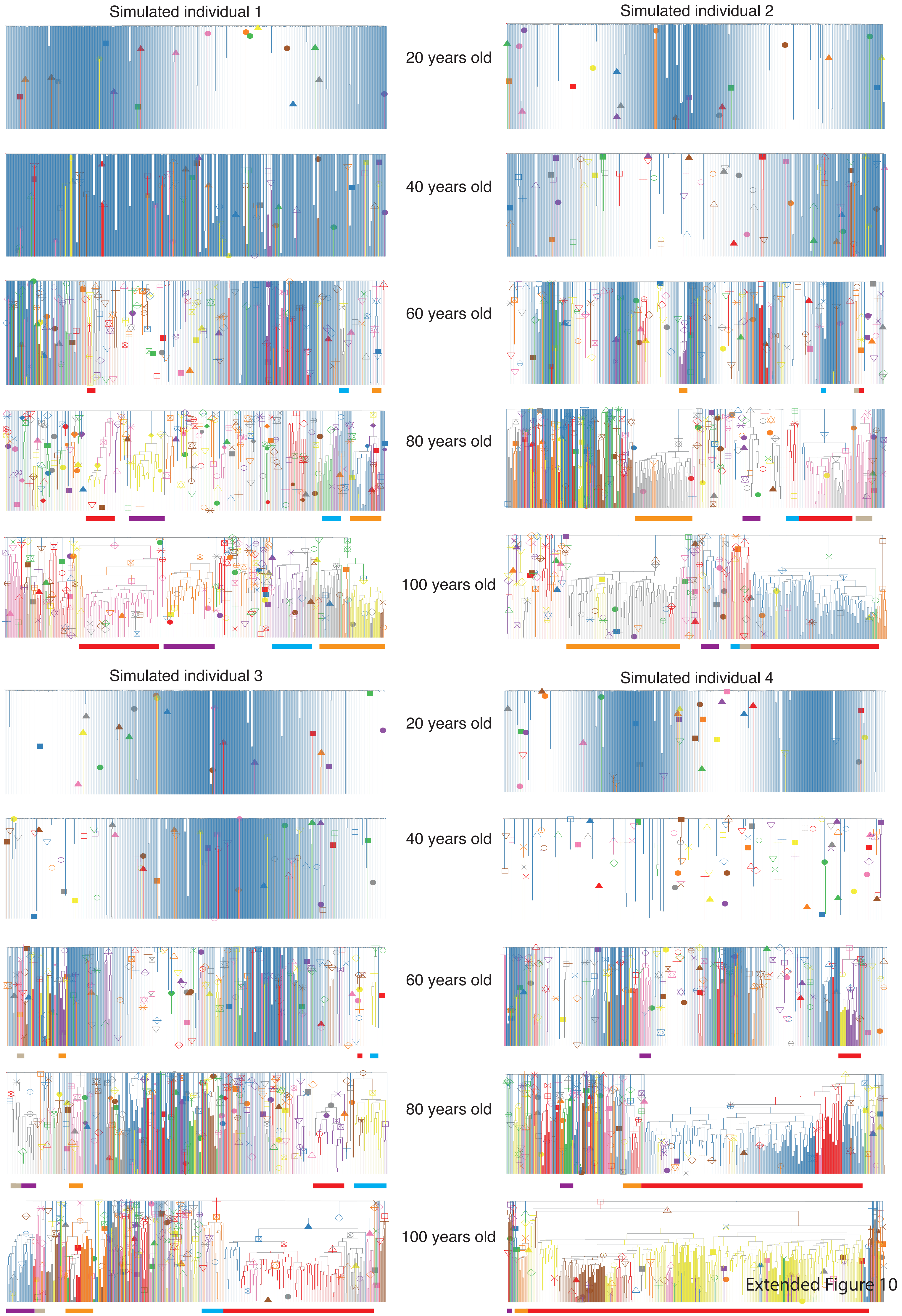


Extended Figure 10
