## Supplementary Methods for "Clonal dynamics of haematopoiesis across the human lifespan"

#### Data reporting

No statistical methods were used to predetermine sample size. The experiments were not randomized and the investigators were not blinded to allocation during experiments and outcome assessment.

#### Samples

In order to obtain representative data from across the whole human lifespan samples were obtained from three sources: **1)** Stem Cell Technologies provided frozen mononuclear cells (MNCs) from two cord blood samples that had been collected with informed consent, including for whole genome sequencing (catalog #70007). **2)** Cambridge Blood and Stem Cell Biobank (CBSB) provided fresh peripheral blood samples taken with informed consent from two patients at Addenbrooke’s Hospital (NHS Cambridgeshire 4 Research Ethics Committee reference 07/MRE05/44 for samples collected pre-November 2019 and Cambridge East Ethics Committee reference 18/EE/0199 for samples collected from November 2019 onwards. **3)** Cambridge Biorepository for Translational Medicine (CBTM) provided frozen bone marrow +/- peripheral blood MNCs taken with informed consent from six deceased organ donors. Samples were collected at the time of abdominal organ harvest (Cambridgeshire 4 Research Ethics Committee reference 15/EE/0152). Details of the individuals studied and the samples they provided are listed in **Fig. 1b**, with additional information in **Table S1**.

#### Isolation of MNCs from fresh peripheral blood samples

Mononuclear cells (MNCs) were isolated using lymphoprep^TM^ density gradient centrifugation (STEMCELL Technologies), after diluting whole blood 1:1 with PBS. The red blood cell and granulocyte fraction of the blood was then removed. The MNC fraction underwent red cell lysis using 1 incubation at 4°C for 15 mins with RBC lysis buffer (BioLegend). CD34 positive cell selection of peripheral blood and cord blood MNC samples was undertaken using the EasySep human whole blood CD34 positive selection kit (STEMCELL Technologies). The kit was used as per the manufacturer’s instructions, but with only a single round of magnetic selection. Bone marrow MNCs did not undergo CD34 positive selection prior to cell sorting.

#### Fluorescence activated cell sorting

MNC or CD34 enriched samples were centrifuged and resuspended in PBS/3%FBS containing an antibody panel consisting of: CD3/FITC, CD90/PE, CD49f/PECy5, CD38/PECy7, CD33/APC, CD19/A700, CD34/APCCy7, CD45RA/BV421 and Zombie/Aqua. Cells were stained (30 minutes at 4°C) in the dark before washing and resuspension in PBS/3%FBS for cell sorting. For all samples ‘HSC/MPP’ pool cells (Lin-, CD34+, CD38-, CD45RA-) were sorted using either a BD Aria III or BD Aria Fusion cell sorter (BD Biosciences) at the NIHR Cambridge BRC Cell Phenotyping hub. The gating strategy is illustrated in **Extended Fig. 1a**. In a subset of individuals, a small number of HPCs (Lin-, CD34+, CD38+) were also sorted. The antibody panel used is shown in **Table S2**.

#### Single-cell colony expansion *in vitro*

Single phenotypic ‘HSC/MPP’ or ‘HPC’ cells were index sorted, as above, into Nunc 96 flat-bottomed TC plates (Thermofisher), containing 100μl supplemented StemPro media (Stem Cell Technologies) but no murine cell feeder layer. The following supplements were added to promote proliferation and push differentiation toward granulocyte, monocyte, erythroid and NK cell types: StemPro Nutrients (0.035%, Stem Cell Technologies), L-Glutamine (1%, ThermoFisher), Penicillin-Streptomycin (1%, ThermoFisher) and cytokines (SCF, 100 ng/ml; FLT3, 20 ng/ml; TPO, 100 ng/ml; EPO 3 ng/ml; IL-6, 50 ng/ml; IL-3, 10 ng/ml; IL-11, 50 ng/ml; GM-CSF, 20 ng/ml; IL-2 10 ng/ml; IL-7 20 ng/ml; lipids 50 ng/ml). Cells were incubated at 37°C and the colonies that formed were topped up with 50μl StemPro media plus supplements at 14 +/- 2 days as necessary. At 21 +/- 2 days a visual size assessment of colonies was undertaken prior to harvesting of cells for DNA extraction. Larger colonies (≥ 3000 cells in size) were transferred to fresh U bottomed 96 well plate (Thermofisher). The U bottomed plates were then centrifuged (500 x g for 5 min), media was discarded, and the cells were resuspended in 50μl PBS prior to freezing at -80°C. Smaller colonies (<3000 but >200 cells in size) were harvested into 96 well skirted LoBind plates (Eppendorf) and centrifuged (800 x g for 5 min). Supernatent was removed to 5-10ul using an aspirator prior to DNA extraction on the fresh pellet. For larger colonies DNA extraction was performed using the DNeasy 96 blood and tissue plate kit (Qiagen). The Arcturus Picopure DNA Extraction kit (ThermoFisher) was used to extract DNA from the smaller colonies. Both kits were used as per the manufacturer’s instructions.

#### Whole genome sequencing of colonies

A recently developed low input enzymatic fragmentation-based library preparation method^1,2^ was used to generate whole genome sequencing libraries from 1-5ng extracted DNA from each colony. Whole genome sequencing was performed at a mean sequencing coverage of 14X (8-35X) on either the Hiseq X or the NovaSeq platforms (Illumina). BWA *mem* was used to align 150bp paired end reads generated to the human reference genome (NCBI build37).

#### Single-base-substitution and indel calling

*CaVEMan* (used for calling SNVs) and *Pindel* (used for calling small indels) were run against an unmatched synthetic normal genome using in-house pipelines^3,4^. *CaVEMan* was run with the ‘normal contamination of tumour’ set to 0.05, otherwise standard settings were used. Default filters were also used, one of which excludes putative SNVs that are present in a large panel of normal samples, so excluding most of the germline single nucleotide polymorphisms (SNPs) from subsequent analysis. It leaves around 30,000-40,000 germline SNPs in most individuals, which represent inherited SNPs that are rare within the population. In addition to the default *CaVEMan* filters, thresholds were set to require putative variants to have a mean mapping score (ASMD) of at least 140 and fewer than half supporting reads being clipped (CLPM=0). *Pindel* was run with standard settings. A custom filter was then used to remove artefacts associated with the ‘low input’ library preparation method, including those due to cruciform DNA structures^5^.

Specifically, the custom ‘low input’ filter incorporates two additional filtering strategies. Firstly, a fragment-based filter, designed to remove overlapping reads that result from the relatively shorter insert sizes produced by this protocol, which can result in the double counting of variants. Secondly, a cruciform filter, which removes erroneous variants introduced due to the incorrect processing of cruciform DNA. For each variant, the standard deviation (s.d.) and median absolute deviation (MAD) of the variant position within the read was calculated separately for positive and negative strands reads. Where a variant was supported by a low number of reads for one strand, the filtering used statistics calculated from the reads derived from the other strand. It was required that either: (a) ≤90% of supporting reads report the variant within the first 15% of the read as determined from the alignment start, or (b) MAD > 0 and s.d. > 4. Where both strands were supported by sufficient reads, it was required for both strands separately to either: (a) ≤90% of supporting reads report the variant within the first 15% of the read as determined from the alignment start, (b) MAD > 2 and s.d. > 2, or (c) at least one strand has MAD > 1 and s.d. > 10.

Following this, *cgpVAF* (another bespoke algorithm) was used to generate a matrix of variant and normal reads at all sites that had a detected variant in any sample from a given individual. These algorithms are available from the Sanger Institute’s Cancer IT GitHub repository (<https://github.com/cancerit>).

Additional filtering on the read count and depth matrices containing several hundred samples per individual was then performed as follows: a) An exact binomial filter was used to remove variants with aggregated count distributions consistent with germline single nucleotide polymorphisms (SNPs)^6^. b) A beta-binomial filter was used to remove low-frequency artefacts, i.e. variants present at low frequencies across samples in a way not consistent with the sample-to-sample variation expected for acquired somatic mutations^6^. c) Sites with a mean depth below 8 and over 40 were removed. d) Thresholds for read count and VAF were used to filter out *in vitro* variants from the remaining mutations using a bespoke script. The thresholds were set to require a minimum variant read count of 2 or more and a variant allele fraction of 0.2 for autosomes and 0.4 for XY chromosomes (**Extended Fig. 2a**). e) For each site normal and variant read counts were aggregated from samples with ≥ 3 variant reads. A one-sided exact binomial test was used to filter mutations inconsistent with a true somatic mutation (p-value < 0.001). f) A final filtering step was the removal of mutations that best mapped to the ‘ancestral’ branch of the SNV-derived phylogenetic tree (only the case for 8 mutations in one individual). Custom R scripts, used for these filtering steps were adapted from Spencer Chapman *et al*^7^ ([https://github.com/emily-mitchell/normal_haematopoiesis/2_variant_filtering_tree_building/scripts/](https://github.com/emily-mitchell/normal_haematopoiesis/2_variant_filtering_tree_building/scripts/d)).

#### Structural variant and copy-number calling

Structural variants (SVs) were called using GRIDDS^8^, with all variants confirmed by visual inspection and by checking if they fit the distribution expected based on the SNV-derived phylogenetic tree. Specifically, GRIDSS with a default setting (version 2.9.4) was used to call SVs. SVs larger than 1kb in size with QUAL >=250 were included. For SVs smaller than 30kb, SVs with QUAL >=300 were only included. Furthermore, SVs that had assemblies from both sides of the breakpoint were only considered if they were supported by at least four discordant and two split reads. We further filtered out SVs for which the standard deviation of the alignment positions at either ends of the discordant read pairs was smaller than five. To remove potential germline SVs and artefacts, we generated the panel of normal by adding in-house normal samples (n=350) to the GRIDSS panel of normal. SVs found in at least three different samples in the panel of normal were removed.

Autosomal copy number aberrations (CNAs) were called using another in-house algorithm, ASCAT (Allele-Specific Copy number Analysis of Tumours)^9^, which was run against a single sample selected from each individual. The matched sample was selected to have a coverage > 15X, no loss of Y and to be a singleton in the phylogenetic tree (no coalescences post birth). The ASCAT output was manually interpreted through visual inspection. ASCAT was unable to accurately call copy number changes on the haploid sex chromosomes in males. Therefore, we ran the in-house algorithm BRASS (BReakpoint AnalySiS)^10^ to generate an intermediate file containing information on binned read counts across 500bp segments of the genome. A comparison of the mean coverage of the X and Y chromosomes was used to call Y loss in individual samples, which was then validated by visual inspection of read depth.

#### Filtering at the colony level

As outlined in the main text we removed some colonies from the dataset due to low coverage (17 samples), being technical duplicates (34 samples) and for showing evidence of non-clonality or contamination (7 samples). We used a peak VAF threshold of < 0.4 (after the removal of *in vitro* variants) as well as visual inspection of the VAF distribution plots to identify colonies with evidence of non-clonality (**Extended Fig. 2a-c**). Visual inspection was particularly important in the cord blood samples where there was greater variability in the distribution of variant allele frequencies due to the lower mutation burden. In these samples the VAF threshold of < 0.4 was therefore less stringently applied.

#### Validation of mutation calls

Mutation spectrums were compared between the set of shared mutations (those present in 2 or more colonies), which are those we have the greatest confidence in, and private mutations (present in only one sample), in which we have lower confidence (**Extended Fig. 2e**). The mutation spectrums are almost identical, providing evidence that are private mutation set does not contain excess artefacts.

#### Mutation burden analysis

SNV and indel burden analysis was performed by first correcting the mutation and indel burden to a sequencing depth of 30, by fitting an asymptotic regression to the data (function *NLSstAsymptotic*, R package *stats*) (**Extended Fig. 2d**). Subsequently, linear mixed effects models were used to test for a linear relationship between age and number of SNVs or number of indels (function *lmer*, R package *lme4*). Number of mutations or indels per colony was regressed using log-likelihood maximisation and age as a fixed effect, with the interaction between age and donor as a random effect. Progenitor samples were excluded from this analysis.

age.mut <- lmer(sub_adj ~ age + (age | donor_id), data = summ_cut[summ_cut$cell_type == "HSC",], REML = F)

age.indel <- lmer(indel_adj ~ age + (age | donor_id), data = summ_cut[summ_cut$cell_type == "HSC",], REML = F)

#### Telomere analysis

Telomere length for each colony sequenced on the Hiseq platform (corresponding to the telomere length in the founding HSC/MPP or HPC) was estimated from the ratio of telomeric to sub-telomeric reads using the algorithm *Telomerecat*^11^. Colonies sequenced on the Novaseq platform could not be used as telomeric reads are removed by a QC step prior to bam file creation.

Linear mixed effects models were used to test for a linear relationship between age and telomere length across all the adults. Cord blood samples were excluded due to possible non-linearity of the relationship between age and telomere length in very early life^12^.

age.tel <- lmer(tel_length ~ age + (age | donor_id), data = subset(summ_cut, summ_cut$platform == "hiseq" & !summ_cut$donor_id %in% c("CB001") & summ_cut$cell_type == "HSC"), REML = F)

Normality testing of the telomere length distributions was performed in R using the Shapiro-Wilk normality test (function *shapiro.test*) and visualised using Q-Q plots and density plots (functions *ggqqplot* and *ggdensity*). The percentage of outlying HSC/MPPs per individual was calculated using the interquartile range criterion (all samples outside the following interval are considered as outliers I = [q0.25−1.5⋅IQR; q0.75+1.5⋅IQR]) (function *boxplot.stats*). For all individuals the only outliers in the data had longer than expected rather than shorter than expected telomeres.

#### Construction of phylogenetic trees

*MPBoot*, a maximum parsimony tree approximation method^13^, was used to build phylogenetic trees of the relationships between their sampled HSPCs. Variants were genotyped as ‘present’ in a sample if 2 or more variant reads supported the variant. Variants were genotyped as ‘absent’ in a sample if 0 variant reads were present at a given site and depth at that site was 6 or more. Sites that did not fall into either of the above categories were marked as ‘unknown’. This genotype matrix was used as the input for *MPBoot*, which outputs unscaled trees with uninformative branch lengths (**Extended Fig. 3a**). A maximum likelihood approach and the original count data was then used to assign each mutation in an individual’s dataset to a branch in their *MPBoot* generated phylogenetic tree (<https://github.com/NickWilliamsSanger/treemut>). Tree edge lengths were then made proportional to the number of mutations assigned to the branch (**Extended Fig. 3b**).

The sensitivity of mutation calling in each sample was used to correct phylogeny branch lengths for sequencing coverage. Sensitivity was calculated as the fraction of known germline variants identified by CaVEMan in a specific sample. Mutation burden was corrected by multiplying the number of variants by 1/sensitivity for private branches. The sensitivity was adjusted to allow for the higher sensitivity on shared branches due to multiple samples containing the variant. Specifically, sensitivity was assessed by measuring the ability of the mutation-calling algorithms to detect heterozygous germline single nucleotide polymorphisms (SNPs) in each sample. Heterozygous SNPs should have the same VAF distribution and sensitivity as true somatic mutations. For private branches, the SNV component of branch lengths was scaled according to:

$$=\frac{n_{SNV}}{p_{i}}$$

Where $n_{cSNV}$is the corrected number of SNVs in sample *i*, $n_{SNV}$is the uncorrected number of SNVs called in sample *i* and $p_{i}$is the proportion of germline SNPs called by the Caveman algorithm in sample *i*.

For shared branches, it was assumed that (1) the regions of low sensitivity were independent between samples, (2) if a somatic mutation was called in at least one sample within the clade, it would also be correctly called (or ‘rescued’) in other samples in the clade (even in lower sensitivity samples). Shared branches were therefore scaled according to:

$$\frac{n_{SNV}}{1-\pi_{i}(1-p_{i})}$$

Where the product is taken for $1-p_{i}$ for each sample *i* within the clade. However, both of these assumptions will not hold true in all cases. Firstly, regions with low coverage are not randomly distributed, with some genomic regions likely to have low coverage in multiple samples. Secondly, while many mutations will be ‘rescued’ in subsequent samples once they have been called in a first sample - because the *treemut* algorithm for mutation assignment uses original read count data, meaning that even a single variant read in a subsequent sample is likely to result in the mutation being correctly assigned - this will not be true in every case. Some samples with very low coverage have 0 variant reads at a given site will by chance. In this situation, a mutation may not be correctly placed. While these factors may lead to an under-correction of shared branches, this approach provides a reasonable approximation. Corrected SNV burdens for each sample can then be calculated as the sum of corrected ancestral branch lengths back to the root of the phylogeny. **Supplementary Figs. 1-2** show the phylogenies with branch lengths corrected for differences in sequencing depth.

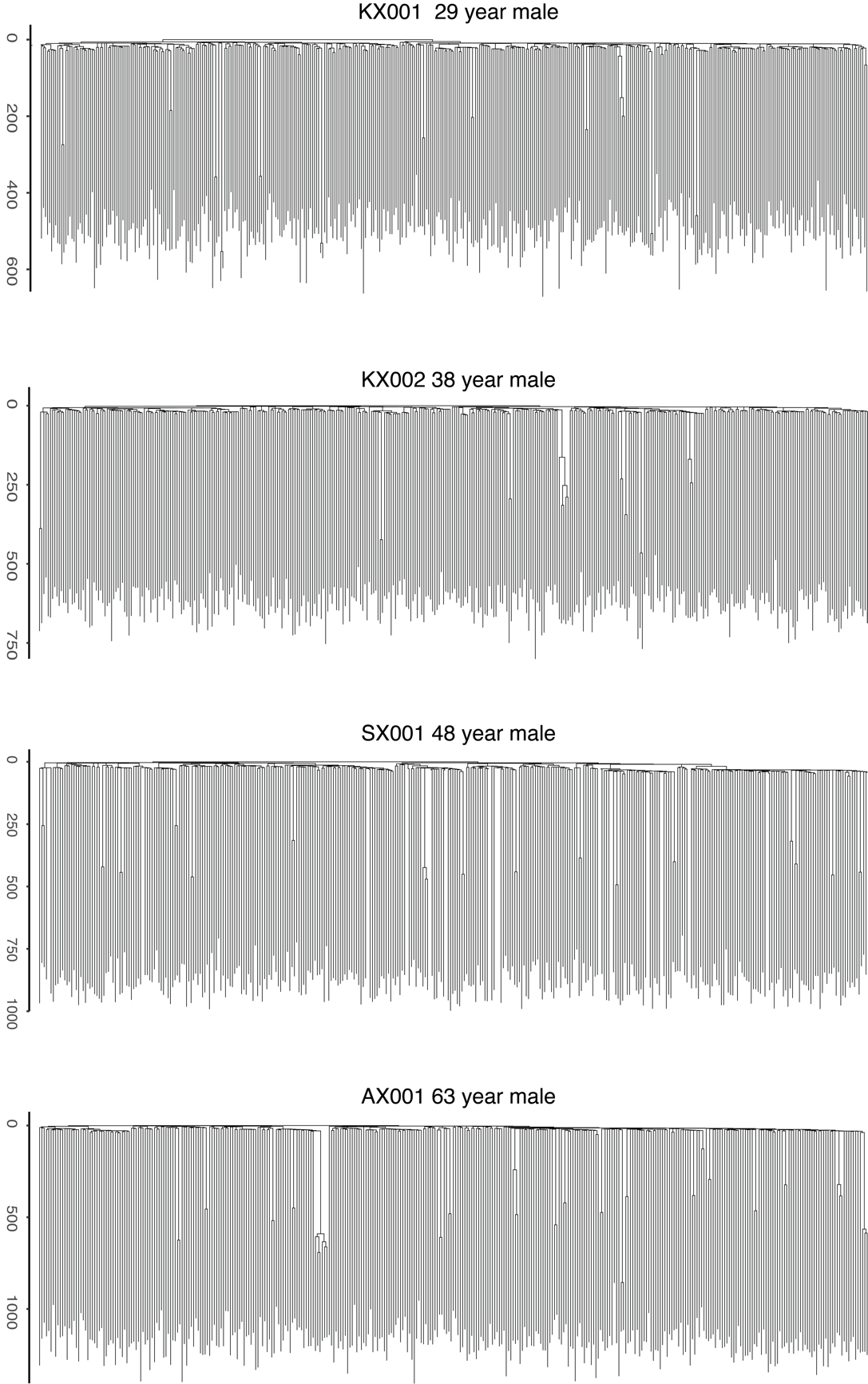

**Supplementary Fig.1|** **Raw phylogenies for the four youngest adult donors.** Phylogenies shown with raw mutation count branch lengths adjusted for sequencing depth of the sample using sensitivity.

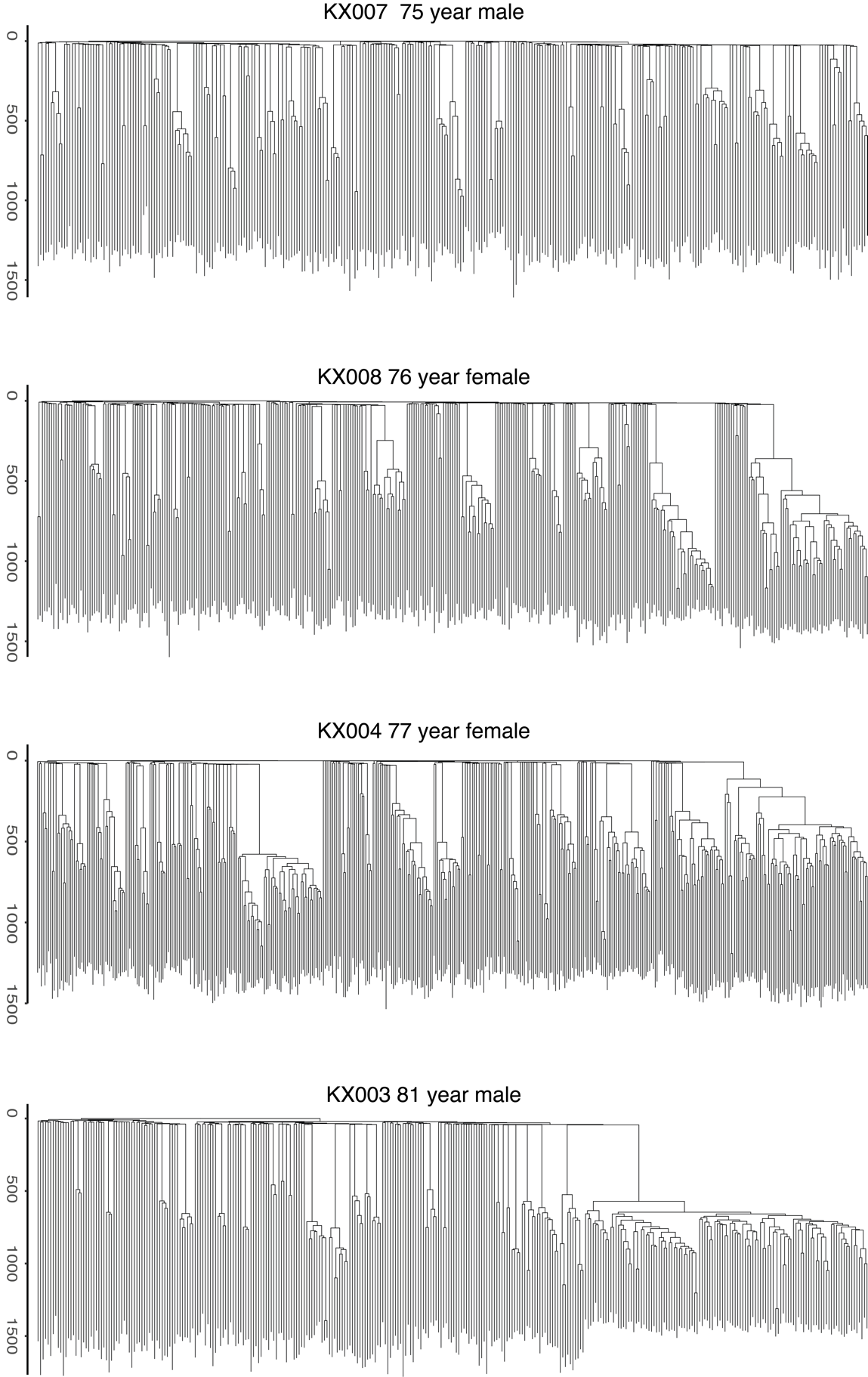

**Supplementary Fig.2| Raw phylogenies for the four elderly adult donors.** Phylogenies shown with raw mutation count branch lengths adjusted for sequencing depth of the sample using sensitivity.

The phylogenies were then made ultrametric using a bespoke algorithm to make all branch lengths equal (**Extended Fig. 3c**, **Supplementary Code**). Starting from the root of the tree and moving progressively towards each tip, the fraction of time for the given shared branch is calculated as the fraction of remaining time times the number of mutations on the given shared branch divided by the mean number of mutations of all descendants from that shared branch. The function is called recursively, updating the fraction of remaining time, as the algorithm moves from root to tip. This algorithm therefore has the property that the most confident timings (nodes near the root) are defined first, anchoring the timings of subsequent, less confident nodes. Given the tight linear accumulation of mutations in HSPCs with age, the mutation branch lengths correspond to molecular time, which can be converted to time in years (**Extended Fig. 3d**).

Validation of the phylogeny
To assess the robustness, internal consistency and stability of the shared variants and inferred phylogenies we used several approaches:

(1) *Bootstrapping of the original read counts.*

One well-established approach to assessing the robustness of individual clades in a phylogeny is to repeatedly bootstrap the mutation matrix and re-build the phylogeny, observing in what proportion of bootstraps each clade is retained. *MPBoot* incorporates a bootstrap approximation method. However, somatic data has a well-established ‘root’ (the human reference genome) which makes this approach less applicable in our setting where we have high confidence in early splits that our supported by multiple samples, even if the numbers of mutations on the branch are low. With our data type the major cause of uncertainty is knowing exactly which cells carry a mutation, and what impact this would have on the inferred tree structure. Therefore, to better assess this type of uncertainty, we used an alternative bootstrapping approach as per Spencer Chapman *et al*^7^. Specifically, we used a partially-filtered mutation set and bootstrapped the read counts for each colony at each locus. We then subjected the raw read count data to the same filtering and phylogeny-building approach as was used on the original data, with 1000 replicates per individual. The only exception was the beta-binomial filter. This was applied to the simulated data before the read count boot-strapping step. As with conventional approaches, the bootstrap phylogenies were then compared to the observed phylogeny to assess the proportion of bootstraps in which each clade is retained or lost. This was compared to the conventional mutation bootstrapping approximation performed by *MPBoot* (**Supplementary Fig. 3a**). Quartet divergence and Robinson-Foulds similarities were calculated using the tqDist algorithm40 implemented in the R package Quartet v1.2.041. The bootstrapping analysis was performed for one of our elderly adult HSCPC phylogenies to ensure the finding of clonal expansions in the phylogeny was robust. We found the bootstrap phylogenies had high correlation to the observed phylogeny (**Supplementary Fig. 3b**) with a median Robinson-Fould similarity of 0.951 and quartet divergence of 0.999.

(2) *Assessment of internal consistency of genotype matrices using the disagreement score.*

This score is based on the observation that in a perfect phylogeny any pair of mutations should either be in discrete clades or nested one within the other. To test the consistency of our data with this assumption we calculated a ‘disagreement score’. For every pair of loci the number of cells in disagreement with this assumption was calculated. The mean score across all pairs was then calculated in such a way that cells with unknown genotypes were assumed to be in agreement. These scores from all the observed phylogenies were then compared to scores generated from random shuffles of the corresponding genotype table, internal to each locus. In this way the disagreement score in the observed genotype table can be compared to one that has been randomly generated. In all the observed trees the ‘disagreement score’ was extremely low compared to that obtained after random shuffling (**Supplementary Fig. 3d**), showing our data has high internal consistency and the phylogenies are close to that expected in a perfect phylogeny (which would have a disagreement score of 0).

(3) *Comparison of the MPBoot phylogeny with other phylogeny inference methods.*

To assess the reliability and stability of the phylogeny generated by MPBoot we used the same genotype data as input into two alternative algorithms: IQ-TREE37 and SCITE38. IQ-TREE is a stochastic algorithm which infers phylogenies using maximum-likelihood. The Jukes-Cantor-type model for binary data was used as an appropriate model for single-cell whole-genome data.  SCITE is an algorithm designed for somatic single-cell data. It uses Markov chain Monte Carlo sampling with an error model that takes potential false positives and false negatives into account for tree scoring. We used false positive and false negative rates of 0.001. Both these alternative algorithms produced phylogenies for KX003 (81 year male) with high agreement to the original *MPBoot* phylogeny with Robinson-Foulds distances of <0.05 for both comparisons (**Supplementary Fig. 3e,f**).

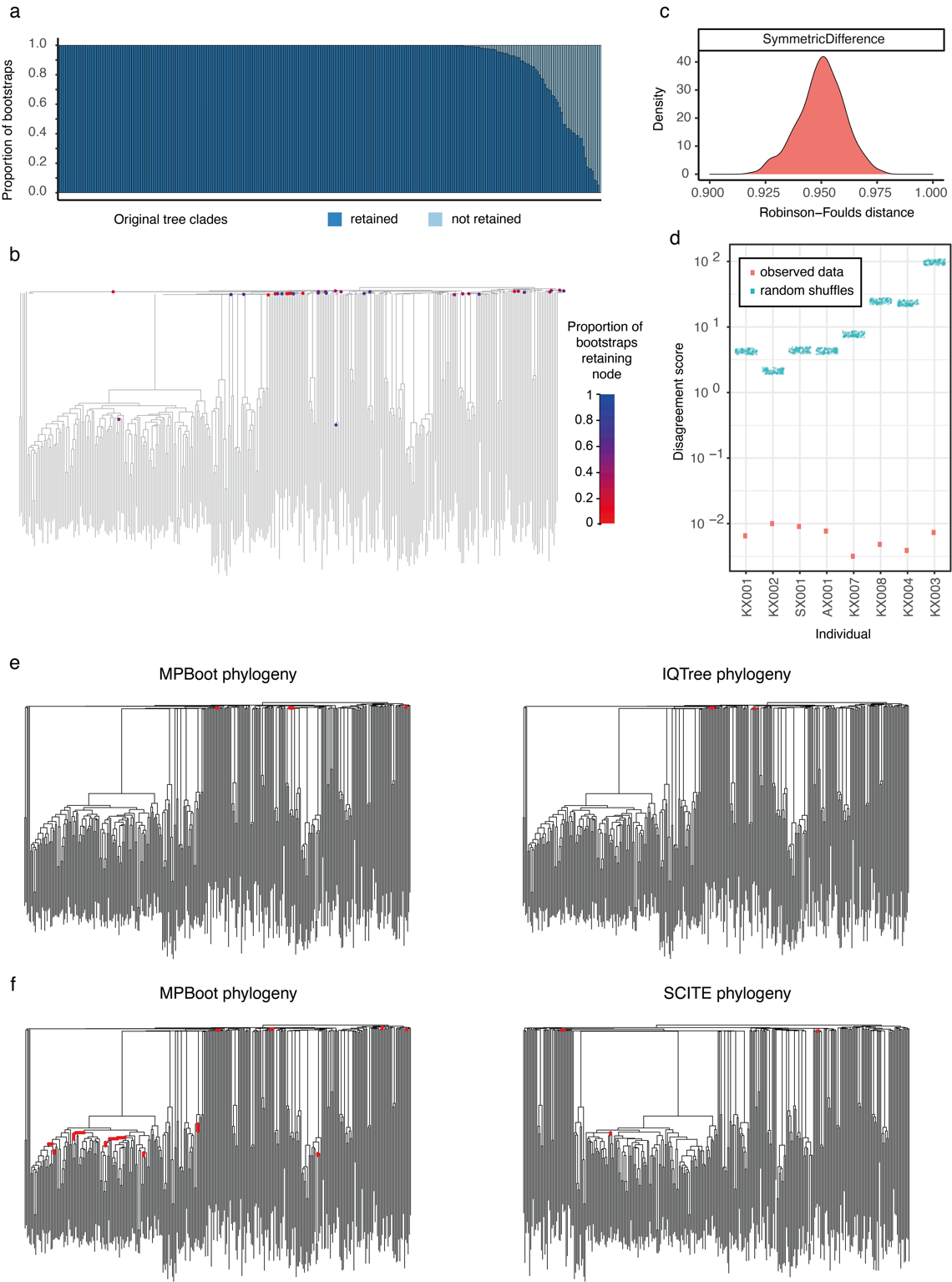

**Supplementary Fig.3|Phylogeny benchmarking. a,** Robustness of each clade in the KX003 phylogeny (81-year male) using bootstrapping of the raw sequencing read count data. The proportion of bootstraps in which a clade is retained is shown, ordered by decreasing robustness. **b,** KX003 phylogeny annotated to show all nodes that have <90% bootstrap support. The nodes are highlighted with the average bootstrap support value. **c,** Comparison of the sequencing read count bootstrap trees to the original trees by Robinson-Foulds similarity. **d,** Internal consistency of the genotype matrix for each adult individual as demonstrated by the disagreement score. The random shuffles have been displayed ‘jittered’. A perfect phylogeny has a score of zero. **e-f,** Comparison of KX003 phylogenies generated by MPBoot and by the alternative phylogeny inference methods IQTree and SCITE.

Inferring HSC population size trajectories

The R package *phylodyn*^14^, provides a well-established approach to inferring historic population size trajectories from the pattern of coalescent events (more specifically the density of these events in historic time blocks) in a phylogenetic tree created from a random sample of individuals in the population. It’s use has been pioneered in pathogen epidemiology^14^ and has also been previously applied to HSPC data from a single individual^15^. **Extended Fig. 5b and 6a** show how *phylodyn* can accurately recover simulated population trajectories using sample sizes similar to those we have used and illustrates how the number of coalescent events in a given time window in the tree informs on population size through time (assuming a constant rate of HSC symmetric cell division and a neutrally evolving population). We used *phylodyn* to infer historic changes in LT-HSC *Nτ* from the ultrametric phylogenies of the four youngest adults in the cohort (**Fig. 4a**).

#### Using Rsimpop to simulate HSC populations

Simulations of complete HSC populations from conception to the age of sampling were performed for each individual using the R package *rsimpop*^16^ (<https://github.com/NickWilliamsSanger/rsimpop>*). Rsimpop* utilises a birth-death model with specified somatic mutation accumulation rate and symmetric cell division rate, to simulate a complete HSC population. Each cell within the population has a rate of symmetric division and a rate of symmetric differentiation (or death). Asymmetric divisions do not impact on the HSC phylogeny and are not accounted for in the model.

Let $\alpha$ be the background rate of symmetric self-renewal cell divisions, measured in divisions per day. We model selective advantage of driver containing clone $i$ as $s_{i}$. The increased rate of symmetric division $\alpha_{i}=\alpha(1+s_{i})$. We assume during the early population growth phase that the total population grows unrestrained by death. Once the specified population size, $N$, is reached (within the first few years of life) then the death rate, $\beta$, for each cell matches the average division rate in the full population:

$$\sum^{cells} \beta=\sum^{wild type cells} \alpha+\sum_{i} \sum^{cells in clone i} (1+s_{i})\alpha$$

Thus giving

$$\beta=\frac{\left( N-\sum_{i} N_{i} \right)\alpha+\sum_{i} N_{i}(1+s_{i})\alpha}{N}$$

In the case of a single driver mutation containing clone with selection coefficient $s$ then the deterministic phase behaviour is governed by a logistic growth function:

$$N_{m}=N\frac{1}{1+exp\left( -\alpha s\left( t-t_{m} \right) \right)}$$

For some constant $t_{m}$ (see Williams *et al)*^17^*.*

In the early stages of the exponential growth process, it exhibits an annual rate of growth $S$:

$S=exp\left( \alpha s \right)-1$.

For multiple competing driver mutation containing clones, each with modest population sizes, it is expected that the above single clone approximation will apply for the individual competing clones. Once one or more of the competing clones represents a significant fraction of the overall population then the dynamics will be more complex. For cells containing more than one driver mutation the fitness effect on *S* is additive.

The above model is implemented using the Gillespie algorithm. The waiting time until the next event is exponentially distributed, with a rate given by the total division rate + total death rate. This event is then ‘division’ with probability=total division rate/(total division rate + total death rate). If the event is ‘division’ then the choice of which cell divides is given by a probability proportional to the cell’s division rate, whereas if the event is ‘death’ then all cells are equally likely to be chosen.

Implementation was in C++ with an R based wrapper as an R package *rsimpop*. The simulator maintains a genealogy of the extant cells, together with a record of the number of symmetric divisions on each branch, the absolute timing of any acquired drivers and the absolute timings of branch start and end. The package also provides mechanisms for sub-setting simulated genealogies whilst preserving the above per branch information.

#### HSC population size modelling

We first investigated simple neutral models of HSC populations (from which selection is absent). The cell phylogenies, constructed from singe cell genomes, include estimated branch lengths, from which we can calculate node heights, and hence the time intervals between successive coalescent events. In the case of a neutral model, the genomic data provides information about the trajectory of the product *Nτ* (population size x time between symmetric self-renewal cell divisions).

However, the genomic data cannot provide information separately about N (population size), or *τ* (time between symmetric cell divisions). Furthermore, in the case of a neutral model, all the information provided by the genomic data, about the trajectory of the product *Nτ*, is contained in sequence of inter-coalescent intervals calculated from the phylogeny. This sequence of inter-coalescent intervals is precisely the information which the *phylodyn* package uses to infer the trajectory of the product *Nτ*.

Here, our aim was to perform additional Bayesian inferences about the parameters of neutral models from the phylogenies. Specifically, we want to compute marginal posterior densities (providing point estimates accompanied by credible intervals) for the ‘LT-HSC *Nτ*’ parameter for the first 2-3 decades of life, and two additional parameters representing the midlife fold-change in *Nτ* (elderly donors only), and late-life fold-change in *Nτ* (all donors).
We chose flat prior densities on wide intervals (**Extended Fig. 7a**) to represent prior uncertainty about the values of these parameters, so that the resulting the marginal posterior densities could be compared with the inferences from the *phylodyn* package.

An additional motivation for performing these Bayesian inferences on neutral models, was to enable us to perform posterior predictive checks (PPC), in order to decide if the observed phylogenies are compatible with neutral models. Note that a separate donor-specific posterior distribution was generated (sampled) for each donor (donor-specific ABC), and a separate donor-specific posterior predictive p-value was computed for each donor (donor-specific PPC). Each donor-specific ABC for the neutral model was performed using the ABC rejection method (R package *abc*)^18,19^.

We used the population trajectory from *phylodyn* to identify the time period prior to the increase related to a ST-HSC/MPP contribution, and the timing of the midlife and late-life fold-change in *Nτ* (**Fig. 4a and Supplementary Fig. 11**). We used our data to inform our choices for the time between symmetric cell divisions, which was set at 1 year (after the initial population growth phase in the first few years of life). We set the rate of mutation accumulation at 15 mutations per year with an additional 1 mutation for every cell division (both of these were drawn from a Poisson distribution centred on the input value).

In the younger individuals (aged < 65) estimates of *Nτ* in the first few decades of life could be made due to the absence of the effect of positive selection (**Extended Fig. 10**). However, in the older individuals (aged > 75), estimates of *Nτ* could not be reliably calculated in the phylogenies due to the confounding effect of positive selection. Here we focussed on using the PPC method to decide whether the neutral model changes in population size (in the form of a bottleneck in the population in mid-life) is compatible with each of the observed trees.

The Bayesian inferences about the parameters of these neutral models were performed using Approximate Bayesian Computation (ABC) methods (in which large numbers of simulations of the data are performed using *rsimpop*, in place of computation of the likelihood function).

In order to apply these methods, the sequence of inter-coalescent intervals was replaced by a set of summary statistics (the ‘number of lineages’ in the tree through time at three points). For each donor, the marginal posterior densities for the parameters of interest are plotted alongside the corresponding prior densities, to illustrate how the data has reduced our uncertainty about the values of these parameters. For each donor, we can also use the sample from the (approximate) posterior distribution (generated by donor-specific ABC) for the parameters of the neutral model, to perform donor-specific PPC.

We first generate a large sample of simulated data sets from the posterior predictive distribution, and from this we can estimate a donor-specific posterior predictive p-value.
The purpose of this donor-specific PPC is to decide if the observed phylogeny obtained from each donor is compatible with the proposed neutral model (while taking account of our uncertainty about the parameter values in the model). Here we are concerned with all features of the observed phylogenies (not only those features which are informative about the parameters of the neutral model). For observed phylogenies and simulated phylogenies, we can compute a chi-squared discrepancy variable which incorporates many summary statistics (including clade size statistics). The posterior predictive p-value is computed from the upper tail-area probability under the distribution of the difference between the simulated chi-squared discrepancy and the observed chi-squared discrepancy^20^. If the p-value is close to zero, then the observed data is extreme (an outlier) compared to the data predicted under the proposed model (taking account of our uncertainty about the parameter values in the model). Thus, when the p-value is close to zero, this is evidence that the observed phylogeny is not compatible with the neutral model.

#### HSC population size estimate

Using a Monte Carlo simulation approach, we sampled from the distributions of each variable 500,000 times, calculating the value of N for each set of randomly sampled variables. This was done using the following distributions: telomeric shortening rate per division: uniform(minimum = 30, maximum = 100); symmetric division rate: uniform(minimum = 0.8, maximum = 1.0); *Nτ* : uniform(minimum = 50,000, maximum = 250,000); average telomere loss per year: uniform(minimum = 30, maximum = 40).

(<https://github.com/emily-mitchell/normal_haematopoiesis/6_population_modelling/scripts/estimating_N.Rmd>)

#### Analysis of driver variants

Variants identified were annotated with VAGrENT (Variation Annotation GENeraTor) (https://github.com/cancerit/VAGrENT) to identify protein coding mutations and putative driver mutations in each dataset. **Table S4** lists the 17 genes we have used as our top clonal haematopoiesis genes (those identified by Fabre *et al* as being under positive selection in a targeted sequencing dataset of 385 older individuals)., whose ‘oncogenic’ and ‘possible oncogenic’ mutations (as assessed independently by EM and PC) are shown in **Figures 2 and 4**. In the same table we also list a more extensive list of 92 clonal haematopoiesis genes that were interrogated. In order to explore a wider set of cancer gene mutations we used the 723 genes listed in Cosmic’s cancer gene census (https://cancer.sanger.ac.uk/census).

#### dN/dS analysis

We used the R package *dndscv*^21^ (<https://github.com/im3sanger/dndscv>) to look for evidence of positive selection in our dataset. The *dndscv* package compares the observed ratio of missense, truncating and nonsense to synonymous mutations, with that expected under a neutral model. It incorporates information on the background mutation rate of each gene and uses trinucleotide-context substitution matrices. The approach provides a global estimate of selection in the dataset, from which the number of excess protein coding, or ‘driver mutations’ can be estimated. In addition, it identifies specific genes that are under significant positive selection. To check for potential bias in our results, we also ran the program excluding sites that are masked by the CaVEMan normal panel in both the numerator and the denominator (a total of 175 million sites) and using a pentanucleotide context correction (rather than the standard trinucleotide context correction). The exclusion of masked sites decreased the global dN/dS estimate, but only by approximately 0.004, while use of the pentanucleotide correction increased the estimate by approximately 0.006.

#### Driver mutation acquisition rate estimation

The dN/dS results were used to derive an estimate of the number of driver mutations acquired per HSC per year. Linear mixed effects models were used to test for a linear relationship between age and the number of non-synonymous mutations. Colonies with a sequencing depth <14 were excluded.

age.non_syn.depth <- lmer(number_non_syn ~ age + (age | donor_id), data = subset(summ_cut, mean_depth > 14), REML = F)

This linear regression analysis found that non-synonymous mutations are acquired at a rate of 0.12/HSC/year (CI_95%_=0.11-0.13) and the dN/dS estimates inform that 1 in 12 to 1 in 34 non-synonymous mutations in the dataset are drivers. We used these estimates in a Monte Carlo simulation approach, sampling from the distributions of each variable 500,000 times, calculating the value of N for each set of randomly sampled variables. This was done using the following distributions: non-synonymous mutation acquisition per year: uniform(minimum = 0.11, maximum = 0.13); fraction of drivers: uniform(minimum = 0.029 (1/34), maximum = 0.083 (1/12)).

(<https://github.com/emily-mitchell/normal_haematopoiesis/5_dNdS/scripts/estimating_driver_acquisition_rate.Rmd>)

#### Y loss analysis

We observe a series of phylogenetic trees from male individuals in which some clades have lost the Y chromosome. By eye, these clades seem to be larger than clades that have not lost Y. To test this formally, we use a randomisation / Monte Carlo test to define the null expected distribution of clade size. For each Monte Carlo iteration, we draw branches of the phylogenetic tree at random - one random branch for each observed instance of Y-loss. These branches are sampled (with replacement) from the set of all extant branches at the matched time-point in that individual, and the eventual clade size of that draw measured. For each simulation, the geometric mean (to allow for the log-normality of observed clade sizes) of clade sizes is calculated. We can then compare the distribution of geometric means from the Monte Carlo draws with the observed geometric mean.

(<https://github.com/emily-mitchell/normal_haematopoiesis/11_LOY_simulations/scripts/Loss_of_Y_simulations.Rmd>)

#### Modelling positive selection in the HSC population

Considering evidence from the dN/dS analysis, we aimed to investigate more elaborate models of HSC population dynamics, by incorporating positive selection acting on driver mutations. Here, as before, we use ABC methods to make inferences about the parameters of the model (incorporating positive selection), and posterior predictive checks (PPC), in order to decide if the observed phylogenies are compatible with this relatively simple non-neutral model (incorporating positive selection). In this non-neutral model, a static HSC population of 100,000 cells undergoing 1 symmetric self-renewal division per year, we explored a range of parameter values for the number of drivers introduced into the population per year, as well as the shape and rate of the gamma distribution used to define the distribution of fitness effects these drivers were drawn from (**Extended Fig. 8a**).

We used a threshold of 5% for the minimum fitness effect of these drivers (equivalent to a selection coefficient of 0.05) as Watson *et al* predicted drivers with a fitness effect of 4% or less could not expand to a VAF > 1% over the human lifespan^22^. We chose flat prior densities on wide intervals (**Extended Fig. 8a**) to represent prior uncertainty about the values of these parameters.

First, a separate donor-specific posterior distribution was generated (sampled) for each donor (donor-specific ABC). The simulations were performed using *rsimpop*, and the donor-specific ABC (ridge regression on the re-scaled, and logit-transformed, parameter values) was performed using the R package *abc*.

Second, we used a sequence of four ABC regression steps to generate a sample from the (approximate) multiple-donor posterior distribution on the combined data from the four oldest donors. The simulations were again performed using *rsimpop*, and the ABC regression steps were again performed using the R package *abc*^18^. In the case of non-neutral models, it is no longer the case that all the information provided by the genomic data, about the parameters of the model, is contained in sequence of inter-coalescent intervals (calculated from the phylogeny). Therefore, additional summary statistics (including clade size statistics) were used in the ABC steps.

A separate donor-specific posterior predictive p-value (donor-specific PPC) was computed for each donor (not only for the four oldest donors), based on the (approximate) multiple-donor posterior distribution on the combined data from the 4 oldest donors. In this case, the sample from each donor-specific posterior predictive distribution was generated by repeated sampling of parameter values from the multiple-donor posterior distribution, and then re-simulating the model (using *rsimpop*) conditional on the donor-specific sample size (number of single cell genomes) and donor age. As before, the posterior predictive p-value is computed from the upper tail-area probability under the distribution of the difference between the simulated chi-squared discrepancy and the observed chi-squared discrepancy^20^.

The purpose of this donor-specific PPC is to decide if the observed phylogeny obtained from each donor is compatible with the simple non-neutral model (while taking account of our uncertainty about the parameter values in the model). If the p-value is close to zero, then the observed data is extreme (an outlier) compared to the data predicted under the simple non-neutral model. This is interpreted as evidence that the observed phylogeny is not compatible with the simple non-neutral model, and that more elaborate models need to be considered.

#### Phylofit estimation of selection coefficients

We used the algorithm *phylofit* to estimate the selection coefficients of known and unknown drivers in our phylogenies. *Phylofit* uses an efficient MCMC approach to model selection within a clade using the probability density of coalescence times and the population size trajectory. As such it can be thought of as a parametric adaptation of the *phylodyn* model.

The starting point for *phylofit* is Equation 1 in Lan *et al* ‘An Efficient Bayesian Inference Framework for Coalescent-Based Nonparametric Phylodynamics’^23^:

$$P\left( t_{1},..,t_{n}|N(t) \right)=\prod_{k=2}^{n} \binom{k}{2}\frac{1}{N(t_{k-1})}e^{-\int_{t_{k}}^{t_{k-1}} \binom{k}{2}\frac{1}{N(t_{k-1})}dt}$$

Where $\left\{ t_{k} | k\in1..n \right\}$ are the timings of the time ordered coalescences belonging to the driver mutation containing clade, $t_{1}$is the first coalescence of the expansion and $t_{n}$ is the sampling time. These times are expressed as the interval between the event and the sampling time (assumed to be isochronous).

Substituting our formula for the cell count of the driver mutation containing clade $N(t)$ (in our case aberrant cell count refers to expanded clades both with and without known drivers) and performing the integral, eliminating terms that do not depend on overall population size, $N$ , the trajectory midpoint, $t^{(m)}$, and the selective coefficient, $\hat{s}=\alpha s,$ we arrive at the following log-likelihood:

$$L\left( t_{1},..,t_{n}|\hat{s},t^{\left( m \right)},N \right)=$$

$$\left( n-1 \right)\log\left( N \right)+\sum_{k=2}^{n} \left( log(1+\exp\left( \hat{s}\left( t_{k-1}-T+t^{(m)} \right) \right) \right)-$$

$$\frac{1}{\hat{s}N}\sum_{k=2}^{n} \left( \binom{k}{2}\exp\left( \hat{s}\left( t_{k-1}-T+t^{(m)} \right) \right)\left( \exp\left( \hat{s}\left( t_{k-1}-t_{k} \right) \right)-1 \right) \right)+$$

$$\frac{1}{N}\sum_{k=2}^{n} \left( \binom{k}{2}\hat{s}\left( t_{k-1}-t_{k} \right) \right)$$

Where recall the annualised selective coefficient is $S=\exp\left( \alpha s \right)-1=\exp\left( \hat{s} \right)-1$

We incorporate this central likelihood equation into a Bayesian model with uniform priors on log(N), $\hat{s}$ and $t^{(m)}$.

$$\hat{s}\sim U\left( 0.001,2 \right)$$

$$t^{(m)}\sim U\left( a,b \right)$$

$$log10(N)\sim U\left( 4,6 \right)$$

$$\boldsymbol{t}\sim Phylo(\hat{s},t^{\left( m \right)},N)$$

Here $Phylo$ is the probability distribution described by the log-likelihood function specified above.

Additionally, assuming unbiased sampling, we can optionally incorporate the number of sampled driver mutation containing colonies $n_{mut}$ out of $n_{tot}$ total colonies as an additional layer in the model:

$$n_{mut}\sim\mathrm{Binomial}\left( n_{tot} ,\frac{1}{1+exp\left( -\hat{s}\left( T-t^{\left( m \right)} \right) \right)} \right)$$

The parameters, $a$ and $b,$ setting the realistic range for the midpoint depend on whether the last component of the model is active and are detailed in the code.

The above models were coded in R and Rstan and inferred using the Rstan implementation of Stan’s No-U-Turn sampler variant of Hamiltonian Monte Carlo method*. Models were fitted across three chains each with 20,000 iterations including 10,000 burn-in iterations.

The input data for this approach is an ultrametric tree. We obtain the ultrametric tree for this analysis using the methods already outlined. The code used to run *phylofit* can be found at (<https://github.com/emily-mitchell/normal_haematopoiesis/7_phylofit/scripts/phylofit.R>)

The *phylofit* algorithm was validated by assessing the correctness of the selection coefficient inference when the algorithm was run on single driver mutation clones with a known selective coefficient. The procedure was as follows:

Simulate population with initial division rate of 0.1 per day ($\alpha=0.1)$until population has grown to the target equilibrium population size.

- Set symmetric division rate to 1 per year ($\alpha=0.5/365)$and simulate neutral evolution until time 𝑇=5 years.
- Save the state of the simulation (*)
- Introduce the driver with the specified selection coefficient.
- If the driver lineage dies out before the sampling age is reached, or has less than 2% clonal fraction at the sampling age, then return to the saved state (*) and continue.

An unbiased sub-sample of cells is taken from the extant population of cells. The *phylofit* algorithm was then applied to the mutant clade in the sub-sampled simulated ultrametric phylogenetic tree.

The algorithm was found to recover the selection coefficients over a range of values of selection coefficient (**Supplementary Fig. 4**).

*Stan Development Team (2020). “RStan: the R interface to Stan.” R package version 2.21.2, <http://mc-stan.org/>.

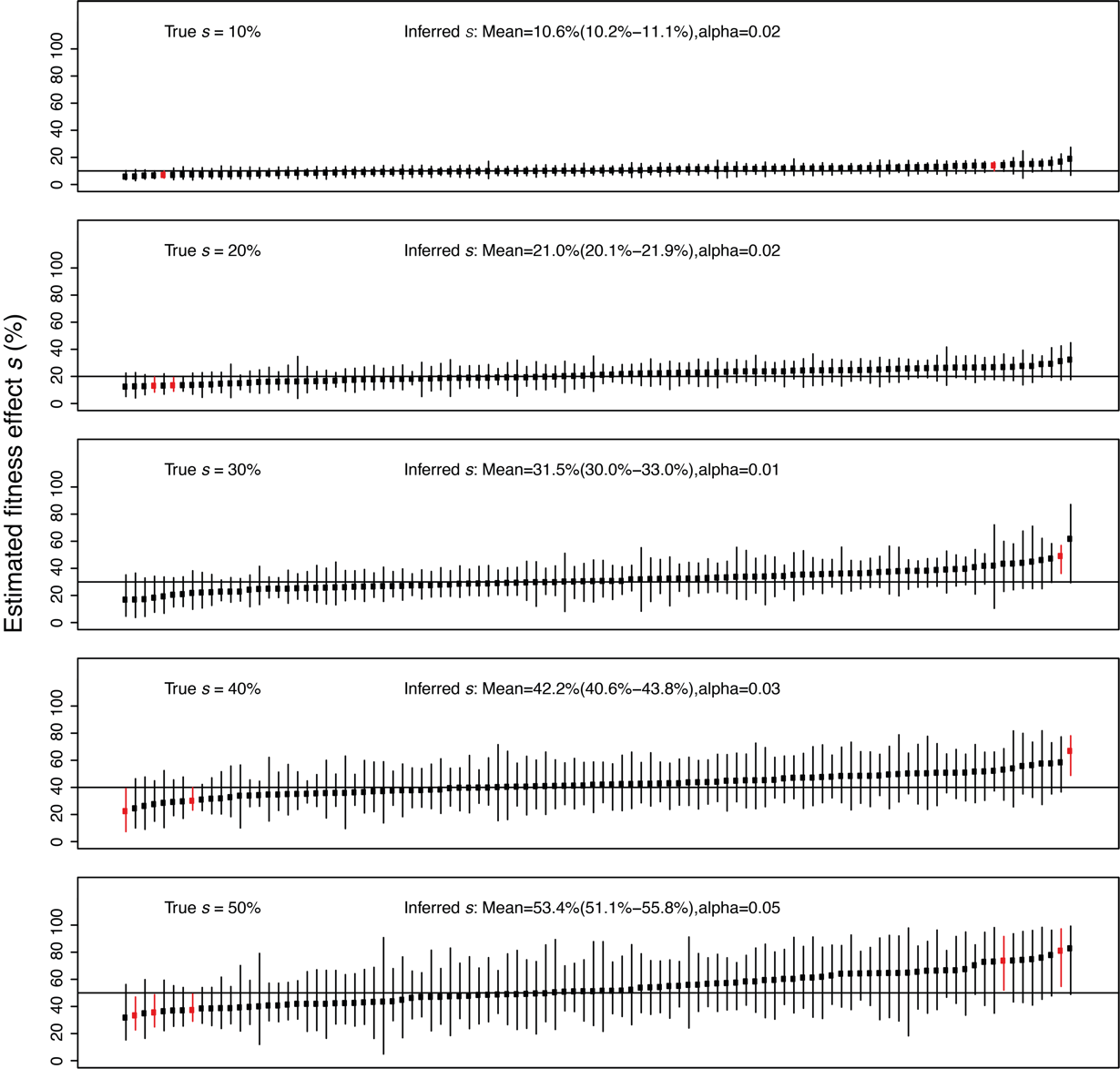

**Supplementary Fig.4|Phylofit Benchmarking.** The inference of annualised fitness effect, *s*. The *phylofit* results (prior *s* range is 0-100% and log10(N) is 4 to 6) are shown for one hundred simulations for each of five values of *s* (=10%, 20%, 30%, 40% and 50%) and N=100,000 cells. The vertical lines show the 95% credibility intervals of the inferred selection coefficients with red lines highlighting instances where the true selection coefficient lies outside the 95% credibility interval (“alpha” is the proportion of such cases). The sample mean estimate of *s* and the corresponding 95% confidence interval are also shown. The benchmarking shows that on average the selection coefficient is accurately recovered with little bias.

#### Analysis of Acute Myeloid Leukaemia (AML) genomes

Mutations in *ZNF318* and *HIST2H3D* were identified in recently described tumour WGS data from 263 patients with AML and myelodysplastic syndromes (MDS) seen at Washington University School of Medicine in St. Louis^24^. Sequencing, initial processing using the *hg38* human reference genome, and full variant calling details for this dataset were described previously^24^. Briefly, identification of SNVs and indels in *ZNF318* and *HIST2H3D* was performed with *Varscan2* run in SNV and indel mode using custom parameters to enhance sensitivity (--min-reads2=3, --min-coverage=6, --min-var-freq=0.02, and --p-value=0.01), along with indel callers *Pindel* and *Manta* using default parameters. Variant calls identified via these approaches were merged and harmonized using a custom python script and annotated with VEP using *Ensembl* version 90. Only one potentially pathogenic variant was found in *ZNF318*. All identified variants are listed in **Table S7**, the majority are likely to be rare germline variants as the sequencing strategy did not incorporate a matched normal sample.

In addition there were no variants in *ZNF318* or *HIST2H3D* reported in 200 cases of AML in the TCGA dataset^25^, compared to 51 cases with DNMT3A mutations. Similarly of 71 AML cases in the JAMA study^26^ 23 cases had DNMT3A mutations but none had *ZNF318* or *HIST2H3D* mutations. Both studies used a tumour/normal WGS approach and reported all somatic variants identified as part of their supplementary datasets.

#### Data availability

Additional data is available on github (https://github.com/emily-mitchell/normal _haematopoiesis). Raw sequencing data is available on EGA (accession number EGAD00001007851).

#### Code availability

Code is available on github:

<https://github.com/emily-mitchell/normal_haematopoiesis/>

#### Acknowledgements

Samples were provided by the Cambridge Blood and Stem Cell Biobank, which is supported by the Cambridge NIHR Biomedical Research Centre, Wellcome Trust - MRC Stem Cell Institute and the Cambridge Experimental Cancer Medicine Centre, UK. This research was supported by the Cambridge NIHR BRC Cell Phenotyping Hub. We thank Timothy Ley for his help with analysis of AML genomes.

#### Author Contributions

#### Funding

#### Competing Interests Declaration

### Supplementary Background

#### Definition of phylodynamics

Phylodynamics is defined as the study of how population level evolutionary processes act to shape phylogenies. To date the phylodynamic approach has been applied most commonly to rapidly evolving viral populations, where it has been used to characterise transmission dynamics^27^.

#### Definition of *Nτ*

One fundamental tenet of phylodynamics is that the frequency of coalescent events in the trees is defined by *Nτ,* where *N* is the population size and *τ* is the generation time. This means that the same phylogeny could be obtained from a population of 100,000 with a generation time of 1 year (*Nτ* = 100,000) and a population of 25,000 with a generation of 4 years (again *Nτ* = 100,000).

#### Phylodynamic principles

It has been shown that in a neutrally evolving population the pattern of coalescent events in a phylogeny created from a random sample of individuals can be used to infer historic population size changes^23^. Specifically, in populations of a constant size (*N*) and generation time *(τ)* there will be more coalescent events (which define individuals that are related) observed in a small compared to a large population. The reason for this difference in phylogenies from small and large populations can be understood by imagining the predicted phylogeny obtained from sampling 10 random individuals from a population of 50 individuals, compared to sampling the same number from a population of 500. We would expect to have a higher chance of sampling siblings and cousins from the smaller population than the larger, which manifests as more coalescent events in the phylogeny from the smaller population (**Supplementary Fig. 5**). This concept can be taken a step further, such that in a population with a fluctuating population size, more coalescent events will be observed in time ‘windows’ where the population size is small compared to when it is larger.

The action of genetic drift in a population means that a proportion of lineages are lost stochastically per unit time, and therefore the older the lineages in a population the more coalescent events will be observed per unit time. This is accounted for in phylodynamic models such as *phylodyn*.

The action of positive selection in a population will alter the pattern of coalescent events in a phylogeny if it results in detectable clonal expansion. This means that inferences of population size and historic population dynamics are only valid in populations that do not have evidence high levels of positive selection. The approach also relies on the random sampling of cells within the population.

#### Application of phylodynamics to stem cell populations

When applied to stem cell populations, *N* is the number of stem cells in the population and *τ* is the generation time. Stem cells can divide in three distinct ways. The first is a symmetric self-renewal division that creates 2 daughter stem cells, so increasing the stem cell population and being the equivalent of stem cell birth. The second is a symmetric differentiation division that results in 2 differentiated daughter cells, which is the equivalent of stem cell death. The final type is an asymmetric division, which produces one stem cell and one differentiated cell and therefore does not alter the size of the stem cell population. It can be seen that only the symmetric self-renewal division results in a daughter progeny that increases the size of the stem cell population. From the phylodynamic perspective stem cell generation time is therefore defined as the time between symmetric self-renewal divisions.

Due to the requirement to sample random cells within a population, it is impossible to robustly apply phylodynamic methodology to somatic stem cells in solid organs. However, the haematopoietic system is the one example of a somatic stem cell population that can be randomly sampled, either through peripheral or cord blood sampling, or by taking a large bone marrow sample from multiple bones. The sampling of large volumes of bone marrow (50-80ml) from deceased organ donors provides the additional advantage that these individuals are highly likely to have had high levels of circulating cytokines at the time of sampling which is known to mobilise HSCs within the bone marrow^28,29^. This makes the haematopoietic stem cell population sampled in these ways the ideal candidate for the application of phylodynamic methods. Nevertheless, interpretation of the phylogenies created by sampling HSCs can be non-intuitive. The next section therefore expands on the simulated phylogenies provided in **Extended Figs. 6 and 7** to aid interpretation of our results.

### Supplementary Simulations

In all the simulated phylogenies illustrated below, the R package *rsimpop* was used to simulate a full neutrally evolving HSC population of size *N*. At a given age 380 cells were sampled at random from the full population to allow creation of comparable phylogenies to those we have obtained from real HSC/MPPs. In all simulations the generation time (*τ*) was set at 1 year meaning *Nτ* = *N*. A *phylodyn* plot is also shown for each phylogeny to show how accurately the population trajectory could be recreated from the pattern of coalescent events. In *phylodyn* plots the downward dips in the trajectory (black line) represent coalescent events in the phylogeny.

#### Effect of population size

Increasing *N* reduces the number of coalescent events in the phylogeny of cells with a fixed generation time sampled from an individual of a given age (**Supplementary Fig. 5**). At age 30 there is loss of resolution in the *phylodyn* output between *Nτ* = 500,000 and *Nτ* = 750,000.

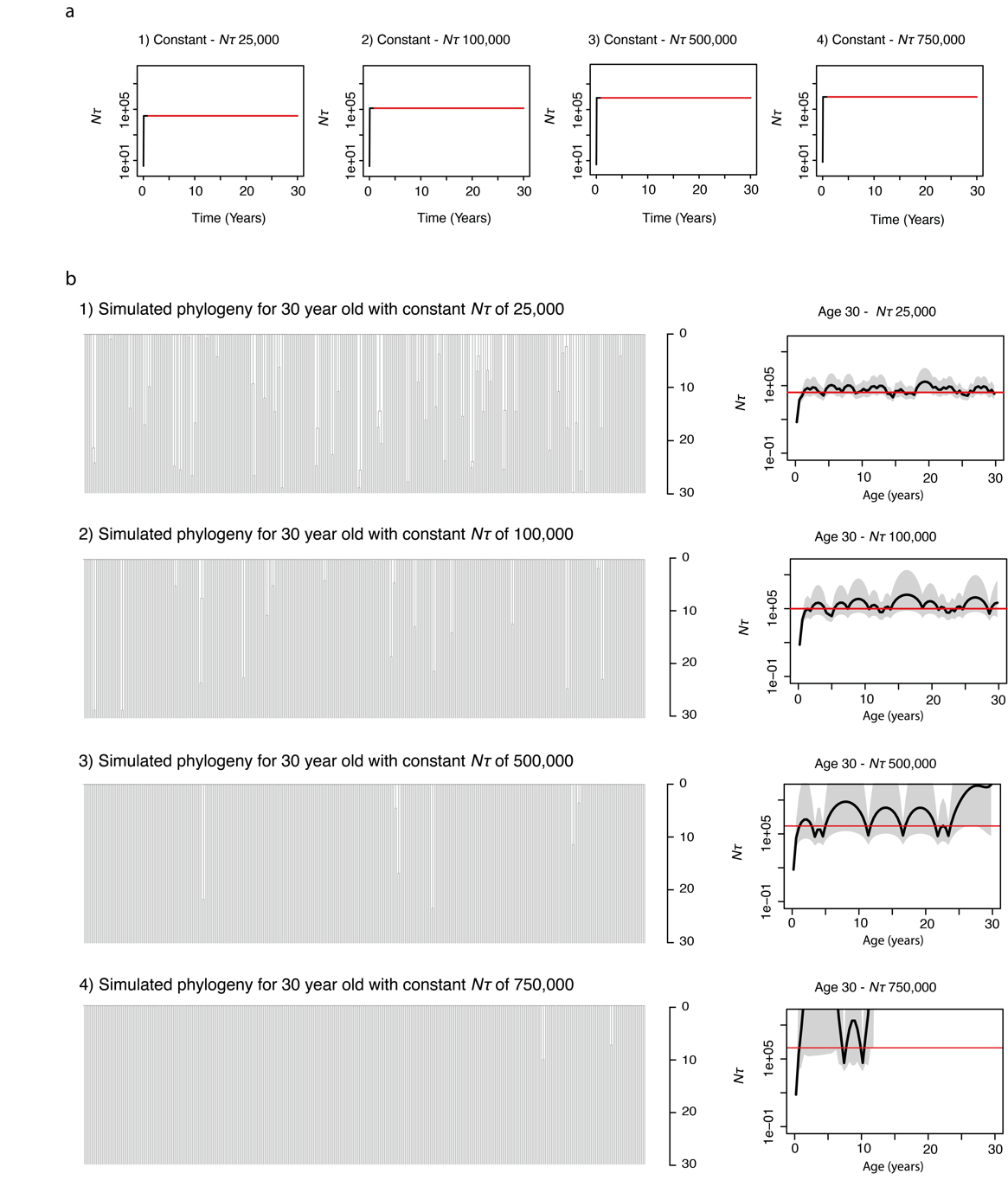

**Supplementary Fig.5|Effect of population size. a,** Trajectories of *Nτ* used as input to *rsimpop* for the simulations to create phylogenies in b. Note the Y axis depicting *Nτ* is on a log scale. **b,** Phylogenies created by randomly sampling 380 cells from the final full simulated population of between 25,000 cells (Phylogeny 1) and 750,000 cells (Phylogeny 4). Phylogenies 1 to 4 are all derived from simulations of the HSC population up to the age of 30 years. Each simulation has an *Nτ* of 100,000. In all cases *Nτ* is the same as the population size (*N*), as the generation time (*τ*) is 1 year. The *phylodyn* trajectories to the right of each simulated phylogeny use the pattern of coalescent events to recover the input trajectories for *Nτ*.

#### Effect of age

Increasing age allows a more accurate estimate of *Nτ* due to the higher number of coalescent events per unit time (**Supplementary Fig. 6**). This increase in the number of coalescent events per unit time for a given population size occurs as a result of genetic drift.

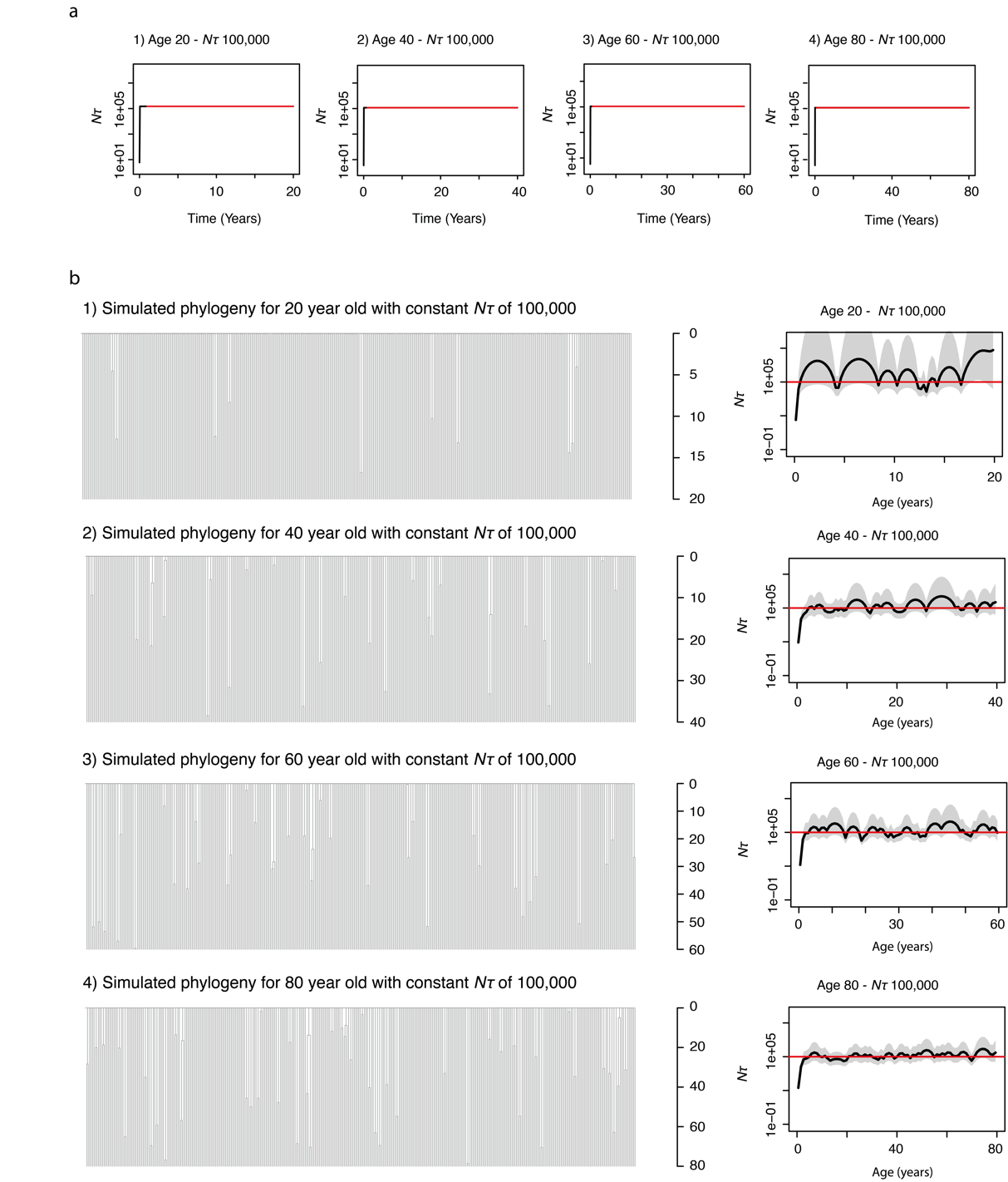

**Supplementary Fig.6|Effect of age. a,** Trajectories of *Nτ* used as input to *rsimpop* for the simulations to create phylogenies in b. Note the Y axis depicting *Nτ* is on a log scale. **b,** Phylogenies created by randomly sampling 380 cells from the final full simulated population of 100,000 cells at between age 20 (Phylogeny 1), age 40 (Phylogeny 2), age 60 (Phylogeny3) and age 80 (Phylogeny 4). Each simulation has a constant *Nτ* of 100,000 In all cases *Nτ* is the same as the population size (*N*), as the generation time (*τ*) is 1 year. The *phylodyn* trajectories to the right of each simulated phylogeny use the pattern of coalescent events to recover the input trajectories for *Nτ*.

#### Population decline

A decline in population size is reflected by an increase in the number of coalescent events captured per unit time as compared to when the population was larger (**Supplementary Fig. 7**). Again, the older the individual the more accurately *phylodyn* is able to recover the true simulated population size trajectory. Decreases in *Nτ* to less than 25,000 can be reasonably accurately captured by *phylodyn*. In younger individuals this is best observed as an increase in the frequency of bumps in the trajectory.

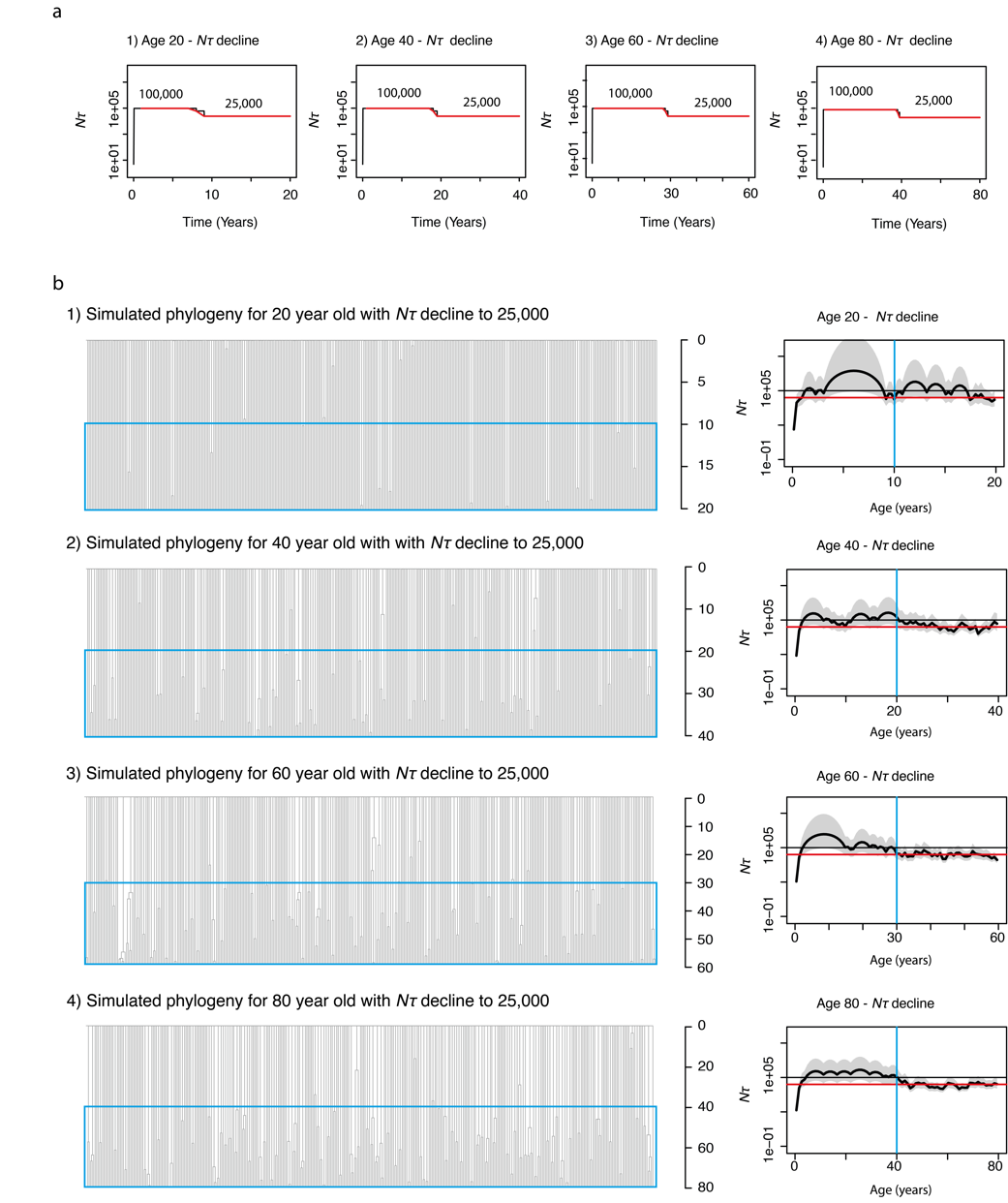

**Supplementary Fig.7|Effect of population decline. a,** Trajectories of *Nτ* used as input to *rsimpop* for the simulations to create phylogenies in b. Note the Y axis depicting *Nτ* is on a log scale. **b,** Phylogenies created by randomly sampling 380 cells from the final full simulated population of 25,000. Each simulation has an initial *Nτ* of 100,000 with a decline to 25,000. In all cases *Nτ* is the same as the population size (*N*), as the generation time (*τ*) is 1 year. The blue boxes indicate the period of time in which the population size is decreased. The *phylodyn* trajectories to the right of each simulated phylogeny use the pattern of coalescent events to recover the input trajectories for *Nτ*. The blue line marks the time of change in *Nτ*.

#### Population growth

An increase in population size is reflected by a decrease in the number of coalescent events captured per unit time as compared to when the population was smaller (**Supplementary Fig. 8**). Again, the older the individual, the more accurately *phylodyn* is able to recover the true simulated population size trajectory. Increases in population size to over 500,000 result in a loss of resolution (and overestimation of *Nτ* in individuals < 40). In younger individuals the change in population size is best observed as a reduction in the frequency of coalescent events (bumps in the trajectory), but the magnitude of the change cannot be accurately determined.

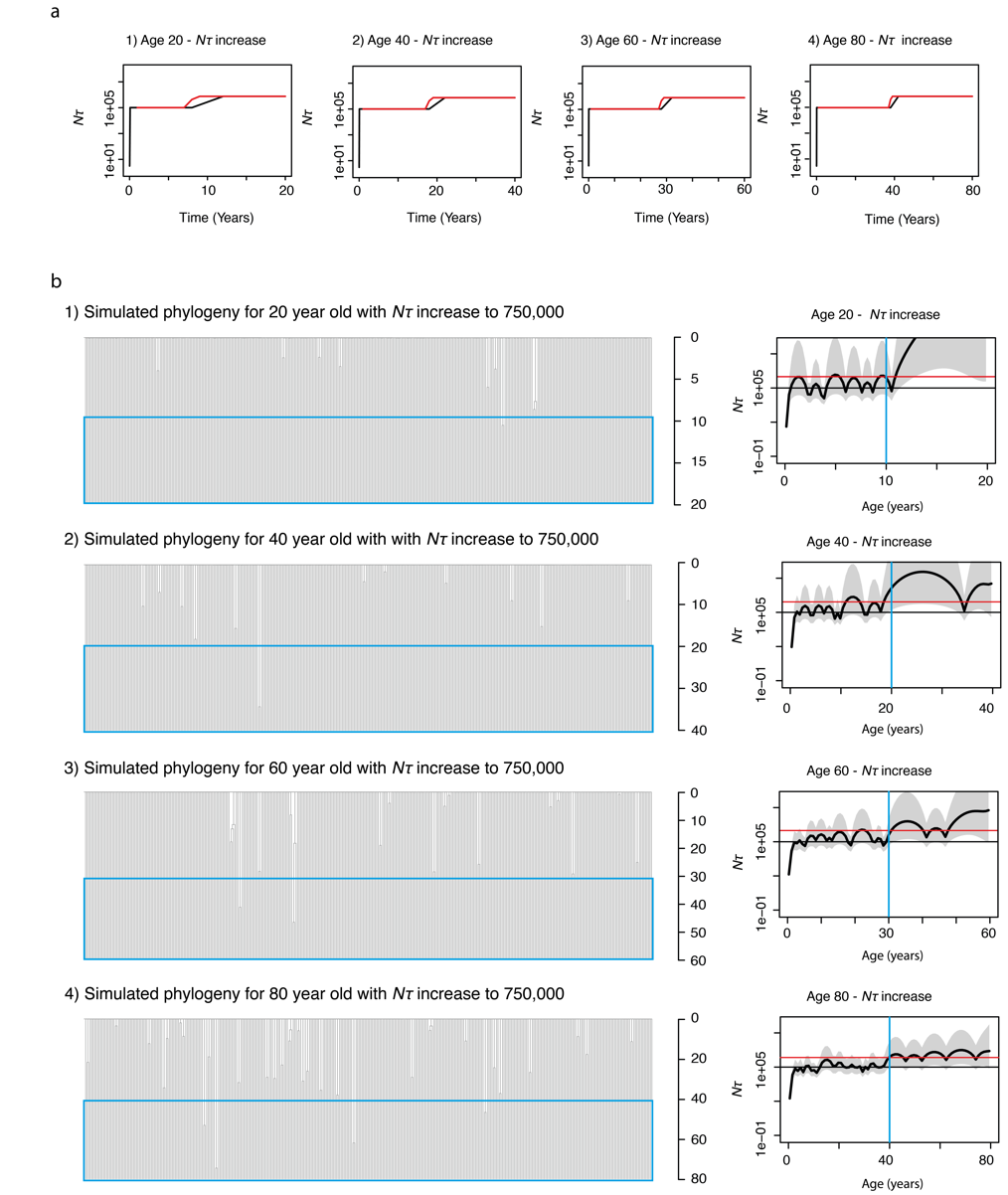

**Supplementary Fig.8|Effect of population increase. a,** Trajectories of *Nτ* used as input to *rsimpop* for the simulations to create phylogenies in b. Note the Y axis depicting *Nτ* is on a log scale. **b,** Phylogenies created by randomly sampling 380 cells from the final full simulated population of 750,000. Each simulation has an initial *Nτ* of 100,000 with an increase to 750,000 in midlife. In all cases *Nτ* is the same as the population size (*N*), as the generation time (*τ*) is 1 year. The blue boxes indicate the period of time in which the population size is increased. The *phylodyn* trajectories to the right of each simulated phylogeny use the pattern of coalescent events to recover the input trajectories for *Nτ*. The blue line marks the time of change in *Nτ*.

#### Population bottlenecks

‘Bottlenecks’ in the population represent periods of time with a reduced population size compared to baseline. These can be recovered accurately by *phylodyn* at all ages, given a reduction to in *Nτ* from 100,000 to 10,000 during the bottleneck period (**Supplementary Fig. 9**).

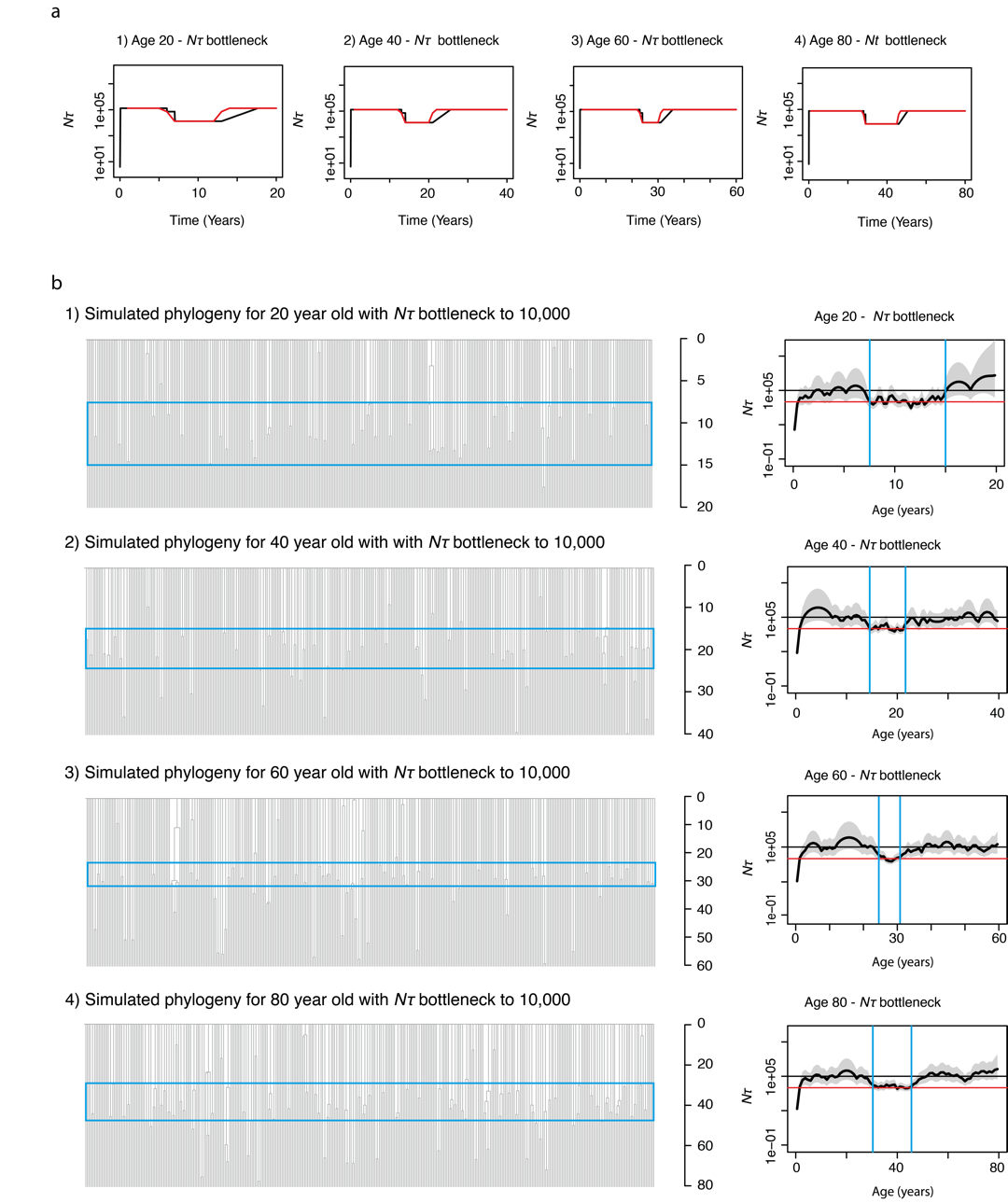

**Supplementary Fig.9|Effect of population ‘bottleneck’. a,** Trajectories of *Nτ* used as input to *rsimpop* for the simulations to create phylogenies in b. Note the Y axis depicting *Nτ* is on a log scale. **b,** Phylogenies created by randomly sampling 380 cells from the final full simulated population of 100,000. Each simulation has an initial *Nτ* of 100,000 with a decline to 10,000 during a period of midlife. In all cases *Nτ* is the same as the population size (*N*), as the generation time (*τ*) is 1 year. The blue boxes indicate the period of time in which the population size is decreased. The *phylodyn* trajectories to the right of each simulated phylogeny use the pattern of coalescent events to recover the input trajectories for *Nτ*. The blue lines mark the times of change in *Nτ*.

#### Positive selection

Positive selection can also be simulated in the phylogenies as illustrated in **Supplementary Fig. 10 and Extended Fig. 10**. These figures show phylogenies drawn from HSC populations where *N* is 100,00 and *τ* is 1 year, with the population as a whole acquiring 200 driver mutations per year, although not all of these will be fixed in the population. The fitness effect of the driver mutations is drawn from a fitness effect gamma distribution (with shape = 0.73 and rate = 33) that incorporates a fitness effect threshold of 5% (**Fig. 5f**). These parameters allow accurate recapitulation of the observed phylogenies across the human lifespan. The simulations illustrate how, although driver mutations are present in the phylogenies of individuals aged below 40, they do not typically impact the pattern of observed coalescences until later in life. This observation provides support for the accuracy of our estimates of *Nτ* in the two youngest individuals in our cohort. In addition, the simulations demonstrate how large clones typically only become detectable after the age of 60, despite the founding driver mutations having been acquired decades earlier (typically in the first 3-4 decades of life). They also illustrate the range of older phylogenies (similar to the range of topologies in our real phylogenies) that can be generated from the stochastic process of driver acquisition. The simulations show how by age 115 years the haematopoietic system could commonly be sustained by just two clones with no known driver mutations, as has been previously reported in a single real individual^30^.

The simulations demonstrate a high prevalence of cells containing multiple drivers by the age of 80, such that in later life clonal competition between driver containing clones with different fitness effects can cause complex clonal dynamics. This is illustrated by that fact that some of the highlighted clades remain stable in size over the last few decades of life, while others may even decline in size. In all illustrated cases one or more ‘fittest’ clones continues to expand into extreme old age.

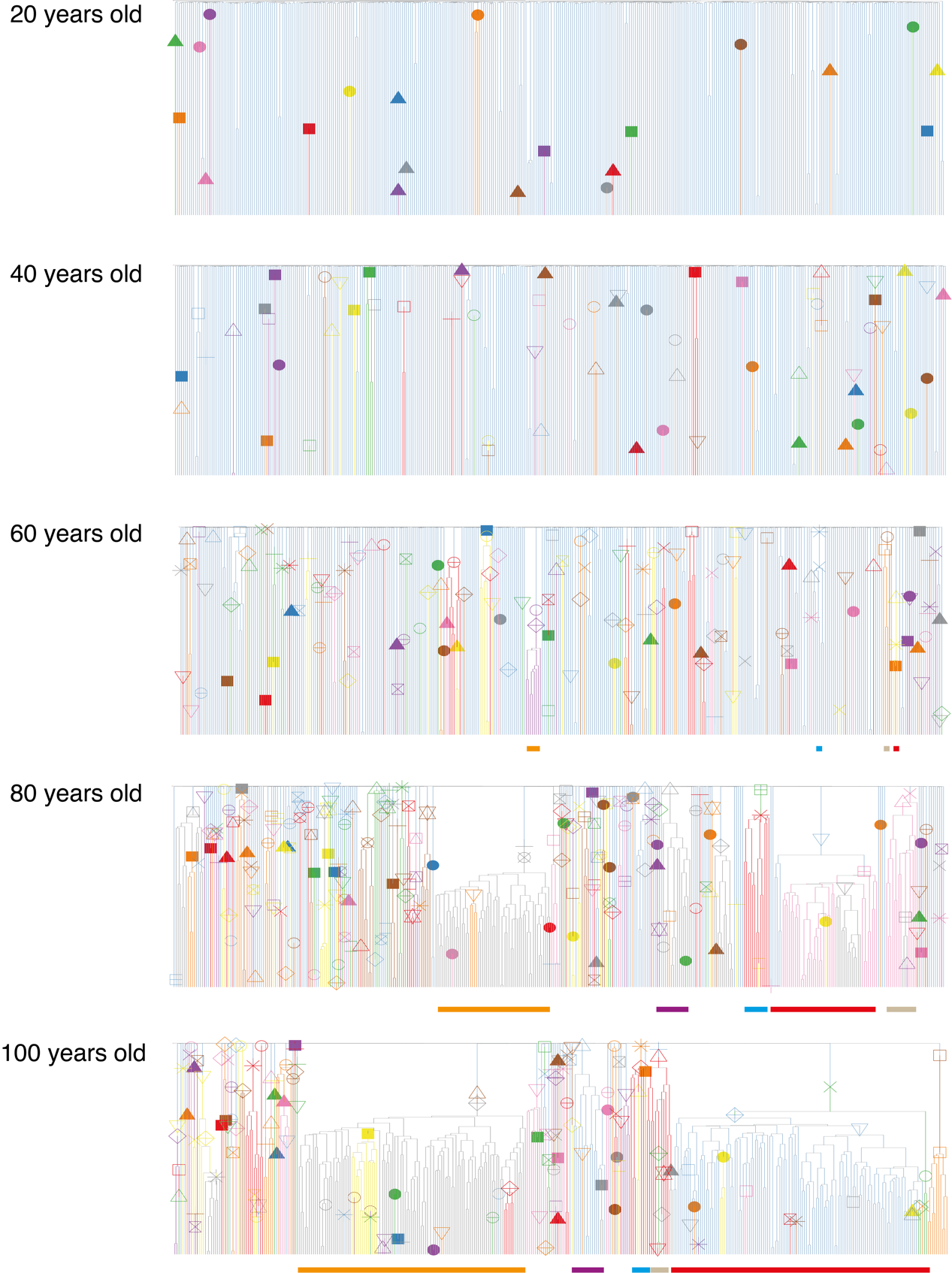

**Supplementary Fig.10|** Phylogenies of 380 cells sampled from a population of 100,000 cells that has been maintained at a constant *Nτ* over life, with incorporation of positively selected ‘driver mutations’. The driver mutations have a fitness effect > 5% (drawn from a gamma distribution with shape = 0.73 and rate = 33) and enter the population at a rate of 200 per year. These are the optimal estimates of these parameters based on our ABC modelling. The inclusion of these driver mutations is able to recapitulate a similar clade size distribution to that observed in the real HSPC phylogenies of the observed individuals across the whole age range. However, including driver mutations does not fully recapitulate the observed lack of coalescent events in the last 10-15 years of life, showing that an increase in *Nτ* over this time is also required to fully recreate the patterns of coalescences in the real phylogenies. Driver mutations are marked with a symbol and their descendent clades are coloured. In all cases *Nτ* is the same as the population size (*N*) as the generation time (*τ*) in all simulations is fixed at 1 year. The symbols / colours are not consistent for driver mutations between plots. The largest clades are therefore coloured in a consistent way beneath the plots to show how their size changes over time. Between the ages of 60 and 80 almost all clades expand.

Supplementary Results

#### Phylodyn trajectories for the older individuals

*Phylodyn* trajectories for the older individuals (age > 75) (**Supplementary Fig. 11**) cannot be reliably interpreted due to the presence of multiple positively selected clades in each case. However, the trajectories were used to inform the timing of populations size changes in the ABC modelling approach for HSC population size (as below).

| Time period | 75 year old | 76 year old | 77 year old | 81 year old |
| --- | --- | --- | --- | --- |
| Change 1 | 10-19 | 15-24 | 10-19 | 15-19 |
| Mid-life bottleneck | 20-45 | 25-50 | 20-45 | 20-50 |
| Change 2 | 46-60 | 51-60 | 46-60 | 51-60 |

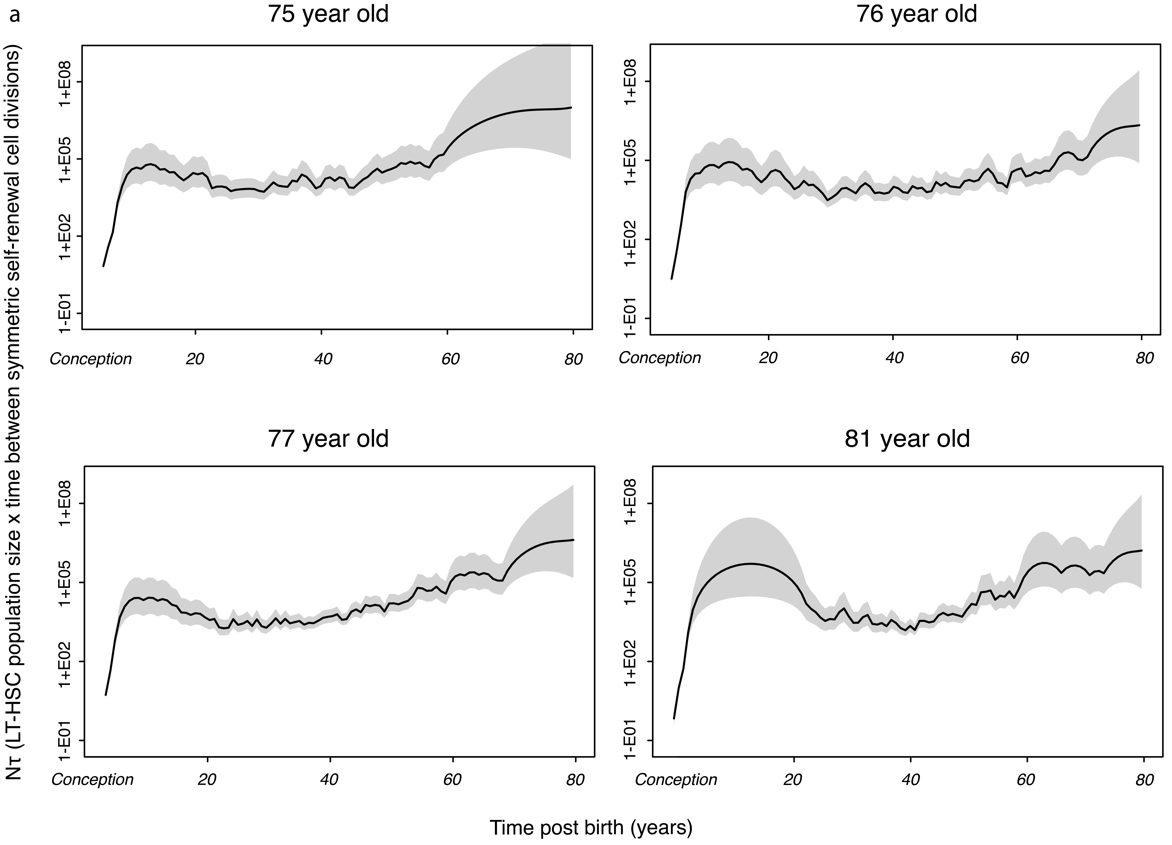

**Supplementary Fig.11|** *Phylodyn* plots illustrating the trajectory of *Nτ* for human LT-HSCs in the four adult donors aged >75 if the pattern of coalescent events in their respective phylogenies was not confounded by the presence of positive selection. The black line represents the trajectory of LT-HSC *Nτ,* with the shaded grey area on either side representing the 95% credibility interval.

#### Posterior distributions for ‘driver modelling’ parameters

Three parameters were included in the ABC driver modelling: 1) rate of the gamma distribution of fitness effects, 2) shape of the gamma distribution of fitness effects 3) Drivers (with s>5%) entering the HSC population of size 100,000 per year. The posterior distribution for the number of drivers (with *s*>5%) entering the HSC population of 100,000 per year is shown in **Extended Fig. 9b**. The posterior distributions for the other two parameters are shown below in **Supplementary Fig. 12**, along with 2D plots showing the relationship between all three parameters.

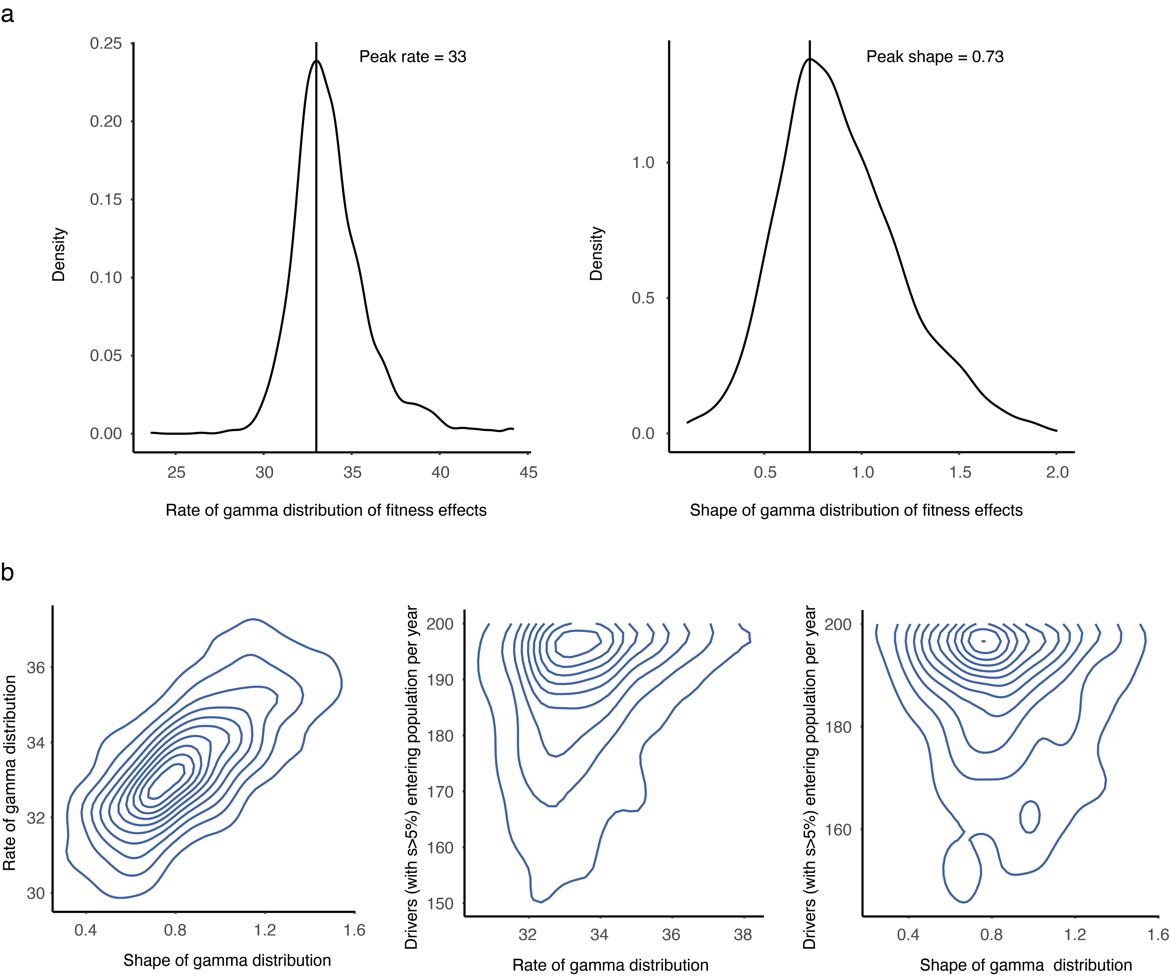

**Supplementary Fig.12| a,** Posterior distribution of the rate and shape of the gamma distribution of selection coefficients. Black line shows peak estimate. **b,** 2D plots showing the relationship between the posterior distributions of the three parameters estimated in the driver modelling approach: 1) rate of the gamma distribution of fitness effects, 2) shape of the gamma distribution of fitness effects 3) Drivers (with s>5%) entering the HSC population of size 100,000 per year.

#### Putative additional novel drivers

Additional possible novel driver genes were identified on the branches of phylogenies leading to expanded clades (**Supplementary Fig. 13** and **Extended Fig. 9c**). Cancer gene variants, as included in the Cosmic Cancer Census v.92 gene set (**Table S4**), which comprises a set of 723 genes causally implicated in cancer development. The top 1500 dN/dS gene hits (**Table S6**) were also interrogated and included only where a manual check of gene function was not incompatible with a possible mechanism to explain clonal expansion.

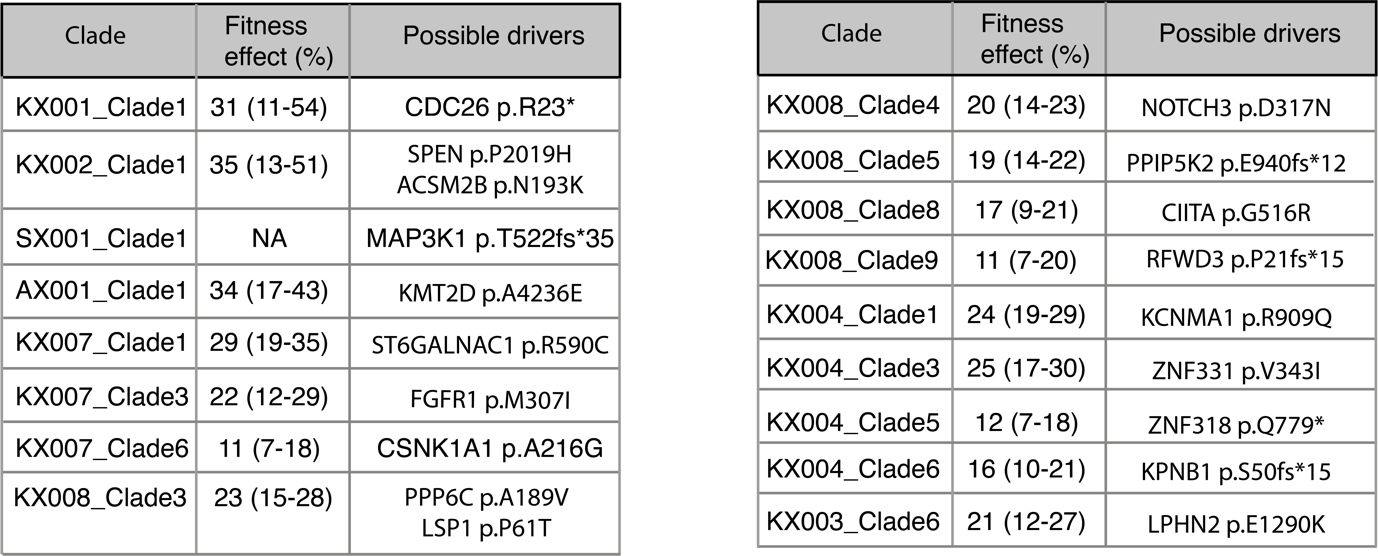

**Supplementary Fig.13|** Table showing putative identified drivers for all four expanded clades in the young adult individuals (clade size 3 – 5). There are strong candidate drivers for these four individuals, whose expansions all have relatively high fitness effects, as would be expected for them to have been able to expand to a detectable level by a young age. The fitness effect was not calculated for SX001_Clade1 due to the clade size being 3 (too small for a meaningful *phylofit* analysis). For the elderly adult individuals a clade size cut off of ≥ 5 was used as clade sizes of up to 4 can occasionally be seen under a neutral model by this age.

3.	Jones, D. *et al.* cgpCaVEManWrapper: Simple Execution of CaVEMan in Order to Detect Somatic Single Nucleotide Variants in NGS Data. doi:10.1002/cpbi.20.

4.	Raine, K. M. *et al.* cgpPindel: Identifying Somatically Acquired Insertion and Deletion Events from Paired End Sequencing. doi:10.1002/0471250953.bi1507s52.

13.	Thi Hoang, D. *et al.* MPBoot: fast phylogenetic maximum parsimony tree inference and bootstrap approximation. doi:10.1186/s12862-018-1131-3.
