## Supplementary material for "Clonal dynamics of haematopoiesis across the human lifespan": Mathematical basis for Approximate Bayesian Computation

### Posterior predictive model checking (PPC) methods which can be applied to Approximate Bayesian Computations (ABC)

### 1 The motivations for posterior predictive model checking

In Bayesian inference, much attention has been given to problems of model choice, and the computation of Bayes factors (Kass and Raftery, 1995). For situations where there is some degree of consensus about the space of alternative models which need to be considered, Bayes factors provide a general solution to the problems of model choice, which is compelling and widely accepted. An alternative approach is to introduce a general model, in which all the alternative models under consideration are included as special cases. We can then employ the usual Bayesian inference strategy of computing the marginal posterior densities for the relevant parameters of the general model.

However, there are many situations where the space of alternative models is vast, and difficult to circumscribe. For example, in population biology there are generally many different time sequences of population sizes and migration rates (representing population expansions, contractions, local extinctions and colonisation events) which are plausible a priori. Increasingly advanced Bayesian methods (Berthier et al., 2002; Beaumont, 2003; Csilléry et al., 2012) have revealed the wide range of models which may be relatively strongly supported by the available data (genotypes of sampled individuals, together with historical records of species abundance or site occupancy). If local adaptation (natural selection) is also considered, the space of (a priori) plausible models is further expanded. Similarly, in the field of developmental biology, and stem cell biology in particular, many different trajectories of stem cell population sizes and rates of cell division, are plausible a priori. Again Bayesian methods reveal that a wide range of models may be relatively strongly supported by the evidence from genomic data (genome sequences of sampled cells, together with targeted deep sequencing of bulk samples), and other data (microscopy, cell sorting, etc). See for example Lee-Six et al. (2018); Chapman et al. (2021). More elaborate models could also be considered. There may be sub-populations within a stem cell population which are spatially or functionally distinct (for example, quiescent verses cycling sub-populations).

Furthermore, some somatic mutations may have effects on rates of cell division, resulting in heritable variation between cells, which is subject to selection. Here too the space of plausible models is vast.

These situations, where the potential space of plausible models is vast, present us with a particular difficulty. Bayes factors and marginal posterior densities can only tell us about the relative strength of the evidence (from the data) supporting each alternative model in the space of models, or each point (parameter vector) in the parameter space. These methods can never tell us that the evidence (from the data) does not support any of the alternative models included in the current space of models, or any of the points in the parameter space, and that other models not yet under consideration, may be more strongly supported by the data. In particular, it may be that the observed data set is an extreme outlier, under all the alternative models included in the current space of models, or any of the points in the parameter space. No amount of inspection of marginal posterior densities, or computation of Bayes factors for specific model comparisons, can alert us to such an inadequacy of the current space of models, or the current parameter space. The recommendations of Bayesian purists to consider an ever larger space of models may be right in principle, but it is not always helpful in the context of an investigation which must deliver conclusions, subject to time constraints.

Attempts have been made to fill this apparent gap in Bayesian methodology. Box (1980) recommended procedures for *model criticism* as part of a Bayesian analysis of the data, including residual plots (in the case of regression models) and prior predictive p-values. The term *posterior predictive model checking* (which can be abbreviated to PCC) was introduced by Rubin (1984) to refer to methods which have been developed for comparing the observed data with posterior predictive distributions, in order to make decisions about the compatibility of the proposed models with the observed data.

In the frequentist approach to statistical inference, tail-area probabilities (computed from the sampling distribution of a statistic) have been used as a criteria for deciding if the observed data should be considered *extreme* under a point hypothesis or composite hypothesis (leading to rejection of the hypothesis). From a Bayesian perspective, we require any decision procedure to take account of our uncertainty about the model, and the parameter values.

Bayesian principles suggest that a *posterior* predictive distribution (rather than a sampling distribution) is the relevant distribution with respect to which we should define what we mean by *extreme* data. This is the probability distribution of a new unobserved data set, conditional on the observed data set. However, the precise definition of an *outlier*, or an *extreme* value, is more elusive. If we find *any* features of the observed data which is extreme under the *posterior* predictive distribution, then we may conclude that none of the models in the space of models (on which we have computed, or sampled, posterior distributions) is compatible with the observed data. This implies that we should consider functions of the data array which are sensitive to every feature (or dimension) of the data array. As we shall see below, we can also consider functions of the data array which include the parameter vector of the model as an argument.

The model $M$ (with parameter vector $\boldsymbol{\theta} \in \mathbb{R}^P$) determines the density $p(\mathbf{y}; \boldsymbol{\theta})$, of the sampling distribution, for each value of the parameter vector $\boldsymbol{\theta}$ in a parameter space $\mathcal{R} \subset \mathbb{R}^P$. In order to determine a posterior predictive distribution for the model $M$, we need to know the observed data array $\mathbf{Y}_0$, and we need to specify the prior density $\pi(\boldsymbol{\theta})$ on the parameter space $\mathcal{R}$. In order to completely determine a posterior predictive distribution it is also necessary to know which functions of the data array are constrained to have the same values for the new unobserved data array as they have for the observed data array. These functions of the data array are what Gelman et al. (1996) (pages 754–755) refer to as *auxiliary statistics*. Let $\boldsymbol{u} = (u_1, \ldots, u_C) = \boldsymbol{u}(\mathbf{y})$ denote the vector of auxiliary statistics $(u_1(\mathbf{y}), \ldots, u_C(\mathbf{y}))$. Let $\mathbf{U}_0 = \boldsymbol{u}(\mathbf{Y}_0)$ denote the vector of auxiliary statistics computed from the observed data array.

The issue of conditioning the new unobserved data to satisfy constraints (specified by the auxiliary statistics) has been discussed by Rubin (1984) and Gelman et al. (1996). In many situations the only auxiliary statistic will be the sample size. In the case of a planned experiment, the vector of auxiliary statistics will be a specification of the experimental design. (This experimental design may be amended to take account of the register of missing data from the observed data set.) In survey sampling, the vector of auxiliary statistics will be a specification of the sampling plan. In developmental biology the data will often includes a representation of a *sample phylogeny* constructed on a sample of single cell genomes (together with mutation assignments to branches), from a series of donor individuals. In this situation the auxiliary statistics will typically be the number of single cell genomes in the sample from each donor, along with the age of each donor. The total number of mutations recorded from each donor, or information about genome coverage, may also be included in the vector of auxiliary statistics.

The conditional density $p\left(\mathbf{y};\boldsymbol{\theta},\boldsymbol{u}\left(\mathbf{y}\right)=\mathbf{U}_0\right)$ of the sampling distribution, is determined by value of the parameter vector $\boldsymbol{\theta}$, and by the value $\mathbf{U}_0$ of the the vector of auxiliary statistics. The posterior density is $\pi\left(\boldsymbol{\theta}\left|\mathbf{Y}_0\right.\right)$. The posterior predictive density is

$$\pi\left(\mathbf{y}\left|\mathbf{Y}_0,\boldsymbol{u}\left(\mathbf{y}\right)=\mathbf{U}_0\right.\right)=\int_{\mathcal{R}}\pi\left(\boldsymbol{\theta},\mathbf{y}\left|\mathbf{Y}_0,\boldsymbol{u}\left(\mathbf{y}\right)=\mathbf{U}_0\right.\right)\boldsymbol{d\theta}, \tag{1}$$

where

$$\pi\left(\boldsymbol{\theta},\mathbf{y}\left|\mathbf{Y}_0,\boldsymbol{u}\left(\mathbf{y}\right)=\mathbf{U}_0\right.\right)=p\left(\mathbf{y};\boldsymbol{\theta},\boldsymbol{u}\left(\mathbf{y}\right)=\mathbf{U}_0\right)\pi\left(\boldsymbol{\theta}\left|\mathbf{Y}_0\right.\right). \tag{2}$$

We may refer to the probability distribution represented by the density $\pi\left(\boldsymbol{\theta},\mathbf{y}\left|\mathbf{Y}_0,\boldsymbol{u}\left(\mathbf{y}\right)=\mathbf{U}_0\right.\right)$ as the *joint posterior predictive distribution.* As we shall see, PPC methods have been developed which can use the *joint* posterior predictive distribution, represented by the density $\pi\left(\boldsymbol{\theta},\mathbf{y}\left|\mathbf{Y}_0,\boldsymbol{u}\left(\mathbf{y}\right)=\mathbf{U}_0\right.\right)$, directly, rather than restricting attention to the usual (*marginal*) posterior predictive distribution, represented by the density $\pi\left(\mathbf{y}\left|\mathbf{Y}_0, \boldsymbol{u}\left(\mathbf{y}\right)=\mathbf{U}_0\right.\right)$.

When it comes to specifying what region of the sample space should be considered to represent extreme events (or outliers) under this posterior predictive distribution, in general we are faced with making arbitrary choices from a very wide range of possibilities. The problem is somewhat simpler in situations where the space of models under consideration is restricted to linear models in which the array of records $\mathbf{Y} = (Y_1, \ldots, Y_d)$, can be expressed in the form

$$Y_r = M_r\left(\boldsymbol{\theta}\right) + S_r\left(\boldsymbol{\theta}\right) Z_r, \tag{3}$$

for $r = 1, \ldots, d$, where $M_r\left(\boldsymbol{\theta}\right)$ and $S_r\left(\boldsymbol{\theta}\right)$ denote respectively the mean and standard deviation of the sampling distribution of the record $Y_r$ (conditional on the parameter vector $\boldsymbol{\theta}$). Under this linear model, the vector of standardised residuals $\mathbf{Z} = (Z_1, \ldots, Z_d)$ is drawn from the $d$-dimensional standard normal distribution

$$\mathbf{Z} \sim N\left(\mathbf{0}_d, \mathbf{I}_d\right). \tag{4}$$

This is a spherically symmetric distribution. It follows directly from the law 4, that the sampling density of the array of records $\mathbf{y}$, is

$$p\left(\mathbf{y}; \boldsymbol{\theta}\right) = \prod_{r=1}^{d} \phi\left(z_r\right), \tag{5}$$

where $z_r$ is the standardised residual

$$z_r = \frac{y_r - M_r\left(\boldsymbol{\theta}\right)}{S_r\left(\boldsymbol{\theta}\right)}, \tag{6}$$

for $r = 1, \ldots, d$; and where

$$\phi\left(z\right) = \frac{1}{\sqrt{2\pi}} \exp\left(-\frac{1}{2}z^2\right), \tag{7}$$

is the probability density function of the standard normal distribution. The sampling density of the array of records $\mathbf{y}$, can therefore also be expressed in the form

$$p(\mathbf{y};\boldsymbol{\theta}) = (2\pi)^{-\frac{d}{2}} \exp\left(-\frac{1}{2} D(\mathbf{y};\boldsymbol{\theta})\right), \tag{8}$$

where

$$D(\mathbf{y};\boldsymbol{\theta}) = \sum_{r=1}^{d} {z_r}^2, \tag{9}$$

is the squared Euclidean distance of the point $\mathbf{z}$ from the origin. From the expression 9 we can see that the sampling density $p(\mathbf{y};\boldsymbol{\theta})$ depends on the data array $\mathbf{y}$ only through the scalar function $D(\mathbf{y};\boldsymbol{\theta})$. This immediately removes much of the arbitrariness from the criteria for what is classified as a outlier. If the value of the squared Euclidean distance $D(\mathbf{Y}_0;\boldsymbol{\theta})$ is extreme (or an outlier), then we can categorise the observed data array $\mathbf{Y}_0$ as extreme.

This approach can also be applied to situations where a vector of standardised residuals can be computed, and where, under the model, the distribution of these standardised residuals approximates the standard normal distribution $N(\mathbf{0}_d, \mathbf{I}_d)$. For example, the same approach can be extended from linear models to generalised linear models (GLMs). This is the basis of the classical *goodness-of-fit* test for Poisson models.

Once we have identified a scalar statistic (function of the data array) which can be used to determine which data arrays are extreme (or outliers), we can define a tail-area probability. If the data array $\mathbf{Y}$ was generated under model $M$ (with parameter vector $\boldsymbol{\theta}$), then the squared Euclidean distance $D(\mathbf{Y};\boldsymbol{\theta})$ represents a draw from a $\chi^2$ distribution (with $\nu = d - 1$ degrees of freedom). The $\chi^2$ distribution (with $\nu$ degrees of freedom) has probability density function

$$h(u;\nu) = \frac{2^{-\frac{\nu}{2}}}{\Gamma\left(\frac{\nu}{2}\right)} u^{\frac{\nu}{2}-1} \exp\left(-\frac{1}{2}u\right), \tag{10}$$

for $u \geq 0$. Let $H(u;\nu)$ denote the cumulative distribution function

$$H(u;\nu) = \int_0^u h(v;\nu)\, dv, \tag{11}$$

of the $\chi^2$ distribution (with $\nu$ degrees of freedom). The tail-area probability for the squared Euclidean distance $D(\mathbf{y}; \boldsymbol{\theta})$ is

$$P(\mathbf{Y}; \boldsymbol{\theta}) = 1 - H(D(\mathbf{Y}; \boldsymbol{\theta}); \nu = d - 1).$$

If the data array $\mathbf{Y}$ was generated under model $M$, with the specified value $\boldsymbol{\theta} = \boldsymbol{\Theta}$, for the parameter vector, then the tail-area probability $P = P(\mathbf{Y}; \boldsymbol{\Theta})$ is a draw from the uniform distribution on the unit interval. An alternative statement of this property is

$$\mathbb{P}[P \leq \alpha] = \alpha, \tag{12}$$

for all $\alpha$ in the unit interval. Here $\mathbb{P}[\cdot]$ denotes probability under the model $M$, with parameter vector $\boldsymbol{\theta} = \boldsymbol{\Theta}$.

Robins et al. (2000) refer to a p-value (tail-area probability) having this property as a *frequentist p-value.* (They also define a more general class of p-values which they refer to as *asymptotic frequentist p-values.*) These authors were motivated by a desire to find p-values which have some kind of Bayesian interpretation, while at the same time retaining a frequentist uniform distribution property. The closely related work of Bayarri and Berger (2000) has a similar motivation.

We can specify a threshold value $\alpha$ (= 0.05, 0.01, 0.001), and adopt the convention that the model $M$ is rejected whenever $P(\mathbf{Y}; \boldsymbol{\theta}) < \alpha$. If we were to apply this procedure to a sequence of data sets $\mathbf{Y}_i$ ($i = 1, 2, \ldots$) drawn independently from the sampling distribution $p(\mathbf{y}; \boldsymbol{\Theta}_0)$, we would compute a sequence of p-values $P_i = P(\mathbf{Y}_i; \boldsymbol{\Theta})$, and the *type I error* rate would then be exactly $\alpha$. Recall that type I error refers to rejecting a model (or null hypothesis) when it is in fact true.

The tail-area probability $P(\mathbf{Y}; \boldsymbol{\theta})$ can be interpreted as an index of how extreme (or surprising) the observed data $\mathbf{Y}$ is under model $M$ (with parameter vector $\boldsymbol{\theta}$). When the tail-area probability is close to zero, this is interpreted as meaning that the model $M$ (with parameter vector $\boldsymbol{\theta}$) is not compatible with the observed data $\mathbf{Y}$, or that the model $M$ fails to explain the observed data $\mathbf{Y}$.

A problem with this version of the frequentist approach is that in general we don't know the value of the parameter vector $\boldsymbol{\theta}$, on which the p-value $P(\mathbf{Y};\boldsymbol{\theta})$ depends. Robins et al. (2000) review the different methods by which the parameter $\boldsymbol{\theta}$ is eliminated from the p-value calculation within the frequentist framework, and the consequences for the frequentist distribution of the p-value. (See also Bayarri and Berger (2000).) A widely used method for eliminating the parameter $\boldsymbol{\theta}$ from the p-value calculation is to insert a point estimate $\hat{\boldsymbol{\theta}}$ of the parameter vector $\boldsymbol{\theta}$. The point estimate $\hat{\boldsymbol{\theta}}$ is given by a function of the data alone. This ensures that $\hat{D}(\mathbf{Y}) = D\left(\mathbf{Y};\hat{\boldsymbol{\theta}}\right)$ is a function of the data alone. We can now compute the frequentist p-value

$$P\left(\mathbf{Y}_0;\hat{\boldsymbol{\theta}},\mathbf{U}_0\right) = \int_{\mathbf{y}:\hat{D}(\mathbf{y})>\hat{D}(\mathbf{Y}_0)} p\left(\mathbf{y};\boldsymbol{\theta}=\hat{\boldsymbol{\theta}},\boldsymbol{u}(\mathbf{y})=\mathbf{U}_0\right) \boldsymbol{d}\mathbf{y}. \tag{13}$$

However, from a Bayesian point of view it would be preferable to have an index of extremeness (or surprise) which takes account of our uncertainty about the value of the parameter vector $\boldsymbol{\theta}$. Box (1980) suggested using the *prior* predictive p-values of the form

$$\begin{aligned} P(\mathbf{Y}_0;\mathbf{U}_0) &= \int_{\mathcal{R}} P(\mathbf{Y}_0;\boldsymbol{\theta},\mathbf{U}_0)\,\pi(\boldsymbol{\theta})\,\boldsymbol{d\theta} \\ &= \int_{\mathcal{R}}\int_{\mathbf{y}:\hat{D}(\mathbf{y})>\hat{D}(\mathbf{Y}_0)} p(\mathbf{y};\boldsymbol{\theta},\boldsymbol{u}(\mathbf{y})=\mathbf{U}_0)\,\pi(\boldsymbol{\theta})\,\boldsymbol{d}\mathbf{y}\boldsymbol{d\theta}. \end{aligned} \tag{14}$$

These prior predictive p-values are also frequentist p-values in the sense of Robins et al. (2000). More specifically, suppose that we adopt the convention that the model $M$ is rejected whenever $P(\mathbf{y};\mathbf{U}_0) < \alpha$. If we were to apply this procedure to a sequence of data sets $\mathbf{Y}_i$ $(i = 1, 2, \ldots)$ generated by sampling $\boldsymbol{\Theta}_i$ from the prior density $\pi(\boldsymbol{\theta})$, and then generating $\mathbf{Y}_i$ from the (conditional) sampling distribution $p(\mathbf{y};\boldsymbol{\Theta}_i,\boldsymbol{u}(\mathbf{y})=\mathbf{U}_0)$, we would compute a sequence of p-values $P_i = P(\mathbf{Y}_i;\mathbf{U}_0)$, and the *type I error* rate would then be exactly $\alpha$.

Guttman (1967) appears to have been the first to advocate the use of *posterior* predictive p-values of the form

$$\begin{aligned}P(\mathbf{Y}_0;\mathbf{U}_0) &= \int_{\mathcal{R}} P(\mathbf{Y}_0;\boldsymbol{\theta},\mathbf{U}_0)\,\pi(\boldsymbol{\theta}\,|\mathbf{Y}_0)\,\boldsymbol{d\theta} \\ &= \int_{\mathcal{R}}\int_{\mathbf{y}:\hat{D}(\mathbf{y})>\hat{D}(\mathbf{Y}_0)} p(\mathbf{y};\boldsymbol{\theta},\boldsymbol{u}(\mathbf{y})=\mathbf{U}_0)\,\pi(\boldsymbol{\theta}\,|\mathbf{Y}_0)\,\boldsymbol{dy}\boldsymbol{d\theta},\end{aligned} \tag{15}$$

as the most appropriate index of extremeness, from a Bayesian point of view. However, Guttman (1967) favoured the terminology of *goodness-of-fit*, rather than extremeness, or surprise. Rubin (1984) also advocated the use of *posterior* predictive p-values (and was apparently unaware of the earlier work of Guttman (1967)). Rubin (1984) uses the terms *model monitoring* and *posterior predictive checks* (which can be abbreviated to PCC), to express the purpose of these methods.

Meng (1994) first enlarged the scope of these methods by pointing out that for any scalar function $D(\mathbf{y};\boldsymbol{\theta})$ of the data array $\mathbf{y}$, and the parameter vector $\boldsymbol{\theta}$, we can define a tail-area probability of the form

$$P(\mathbf{Y}_0;\boldsymbol{\theta},\mathbf{U}_0) = \int_{\mathbf{y}:D(\mathbf{y};\boldsymbol{\theta})>D(\mathbf{Y}_0;\boldsymbol{\theta})} p(\mathbf{y};\boldsymbol{\theta},\boldsymbol{u}(\mathbf{y})=\mathbf{U}_0)\,\boldsymbol{dy}, \tag{16}$$

and therefore we can define a posterior predictive p-value of the form

$$\begin{aligned}P(\mathbf{Y}_0;\mathbf{U}_0) &= \int_{\mathcal{R}} P(\mathbf{Y}_0;\boldsymbol{\theta},\mathbf{U}_0)\,\pi(\boldsymbol{\theta}\,|\mathbf{Y}_0)\,\boldsymbol{d\theta} \\ &= \int_{\mathcal{R}}\int_{\mathbf{y}:D(\mathbf{y};\boldsymbol{\theta})>D(\mathbf{Y}_0;\boldsymbol{\theta})} p(\mathbf{y};\boldsymbol{\theta},\boldsymbol{u}(\mathbf{y})=\mathbf{U}_0)\,\pi(\boldsymbol{\theta}\,|\mathbf{Y}_0)\,\boldsymbol{dy}\boldsymbol{d\theta}.\end{aligned} \tag{17}$$

Meng (1994) introduced the term *discrepancy variable* for functions of the data array $\mathbf{y}$, and the parameter vector $\boldsymbol{\theta}$. The squared Euclidean distance, defined in equation 5, is an example of such a discrepancy variable. The advantage of using a discrepancy variable (such as the squared Euclidean distance defined in equation 5) to define the tail-area probability, is that the resulting criteria for the observed data being *extreme* (or an *outlier*), under model $M$, is not reliant in any way on obtaining an accurate point estimate of the parameter vector $\boldsymbol{\theta}$ of the model. Meng (1994) does not address the difficult problem of the choice of the discrepancy variable with respect to which the tail-area probability is defined.

Meng (1994) emphasises two (closely related) interpretations of the posterior predictive p-value $P(\mathbf{Y}_0; \mathbf{U}_0)$ (as defined in equation 17). Firstly, the p-value $P(\mathbf{Y}_0; \mathbf{U}_0)$ can be interpreted as the probability that the value of the discrepancy variable computed from the observed data array $\mathbf{Y}_0$ is *extreme* under model $M$. Secondly, the p-value $P(\mathbf{Y}_0; \mathbf{U}_0)$ can be interpreted as the (posterior predictive) expected value of the tail-area probability $P(\mathbf{Y}_0; \boldsymbol{\theta}, \mathbf{U}_0)$ (as defined in equation 16), which itself is a frequentist index of surprise or extremeness for the observed data $\mathbf{Y}_0$, under model $M$.

Meng (1994) derived formulae for the posterior predictive p-value for some problems of interest, including the classic Behrens-Fisher problem. Meng (1994) also investigated the sampling properties of the posterior predictive p-value $P(\mathbf{Y}; \mathbf{U}_0)$ whenever the data $\mathbf{Y}$ is drawn from the prior predictive distribution.

Gelman et al. (1996) continued the development of the PPC methodology along the lines suggested by Meng (1994), in which posterior predictive p-values are defined in terms of discrepancy variables (rather than sample statistics). These authors favour the term *assessment* of models, in place of hypothesis *testing*, and often use the term *posterior predictive assessment of model fitness* as an alternative to PPC. Gelman et al. (1996) contains little in the way of guidance on the choice of the discrepancy variable. However, these authors do support the use of the squared Euclidean distance, defined in equation 5, as an appropriate discrepancy variable in a wide range of situations. (Similar advice is also offered in Gelman et al. (2004, 2013).)

The authors Rubin (1984); Meng (1994); Gelman et al. (1996) all emphasise the interpretation of posterior predictive p-values as posterior probabilities. This interpretation does not rely on any frequentist uniform distribution property. However, it would be reassuring to have some guarantee of good frequentist performance for Bayesian procedures based on posterior predictive p-values, and Meng (1994) was able to prove an encouraging result of this type. Suppose that we have a sequence of data sets $\mathbf{Y}_i$ $(i = 1, 2, \ldots)$ generated by sampling $\boldsymbol{\Theta}_i$ from the prior density $\pi(\boldsymbol{\theta})$, and then generating $\mathbf{Y}_i$ from the (conditional) sampling distribution $p(\mathbf{y}; \boldsymbol{\Theta}_i, \boldsymbol{u}(\mathbf{y}) = \mathbf{U}_0)$. For each of these data sets we compute a posterior predictive p-value $P_i = P(\mathbf{Y}_i; \mathbf{U}_0)$. If we adopt the convention that the model $M$ is rejected whenever $P(\mathbf{y}; \mathbf{U}_0) < \alpha$, then the *type I error* rate is

$$\mathbb{P}[P \leq \alpha] \leq 2\alpha, \tag{18}$$

for all $\alpha$ in the range $0 \leq \alpha < \frac{1}{2}$ (Meng (1994), on page 1157). Here $\mathbb{P}[\cdot]$ denotes probability under the *joint prior predictive distribution*

$$\pi(\boldsymbol{\theta}, \mathbf{y} \,|\boldsymbol{u}(\mathbf{y}) = \mathbf{U}_0) = p(\mathbf{y}; \boldsymbol{\theta}, \boldsymbol{u}(\mathbf{y}) = \mathbf{U}_0)\, \pi(\boldsymbol{\theta}). \tag{19}$$

So, a procedure of rejecting the null model whenever the posterior predictive p-value $P(\mathbf{y}; \mathbf{U}_0)$ falls below some small value $\alpha$, has a small *type I error* rate (lower than $2\alpha$).

Gelman et al. (1996) emphasises that it is straightforward to estimate a posterior predictive p-value of the general type defined in equation 17, if we have a Monte Carlo sampler for the posterior density $\pi(\boldsymbol{\theta} \,|\mathbf{Y}_0)$, and a simulator for the (conditional) sampling density $p(\mathbf{y}; \boldsymbol{\theta}, \boldsymbol{u}(\mathbf{y}) = \mathbf{U}_0)$.

Using the Monte Carlo sampler, we first generate a sample of $N$ observation $\boldsymbol{\Theta}_j$ $(j = 1, \ldots, N)$ from the posterior density $\pi(\boldsymbol{\theta} \,|\mathbf{Y}_0)$. For each observation $\boldsymbol{\Theta}_j$, we can then use the simulator to generate a data array $\mathbf{Y}_j$, drawn from the sampling density $p(\mathbf{y}; \boldsymbol{\Theta}_j, \boldsymbol{u}(\mathbf{y}) = \mathbf{U}_0)$. In this way we generate a sample of observations $(\boldsymbol{\Theta}_j, \mathbf{Y}_j)$ $(j = 1, \ldots, N)$ from the *joint* posterior predictive distribution (equation 2). We can now compute the *observed* discrepancy $D(\mathbf{Y}_0; \boldsymbol{\Theta}_j)$ (*realised* discrepancy, in the terminology of Gelman et al. (1996)), and the *simulated* discrepancy $D(\mathbf{Y}_j; \boldsymbol{\Theta}_j)$ (*predictive* discrepancy, in the terminology of Gelman et al. (1996)).

The proportion of observations in the Monte Carlo sample which satisfy the condition $D(\mathbf{Y}_j; \boldsymbol{\Theta}_j) > D(\mathbf{Y}_0; \boldsymbol{\Theta}_j)$, is a Monte Carlo estimate

$$\hat{P}\left(\mathbf{Y}_0;\mathbf{U}_0\right) = \frac{1}{N}\sum_{j=1}^{N}\mathbb{I}\left\{D\left(\mathbf{Y}_j;\boldsymbol{\Theta}_j\right) > D\left(\mathbf{Y}_0;\boldsymbol{\Theta}_j\right)\right\}, \tag{20}$$

of the posterior predictive p-value defined in equation 17. Here $\mathbb{I}\{s\}$ denotes an indicator function which takes the value 1 when the condition $s$ is satisfied, and the value 0 otherwise.

Note however that in the case of the squared Euclidean distance (equation 5), our ability to compute $D(\mathbf{y};\boldsymbol{\theta})$ is reliant on having a formula for the mean $M_r(\boldsymbol{\theta})$ and the standard deviation $S_r(\boldsymbol{\theta})$ (for $r = 1, \ldots, d$). When the model under consideration is a linear model, the required formulae are available. As already mentioned, the same approach can be extended from linear models to generalised linear models (GLMs). Next, we outline how this approach can be extended to other situations.

A widely applicable approach to the task of constructing a discrepancy variable which is sensitive to every feature (or dimension) of the data array, is to first define a vector of summary statistics $\boldsymbol{t}(\mathbf{y}) = (t_1(\mathbf{y}), \ldots, t_D(\mathbf{y}))$, which can be computed from the data array $\mathbf{y}$, and which is sensitive to every feature of the data array.

Given a data array $\mathbf{Y}$, and a parameter vector $\boldsymbol{\theta}$, we can compute the corresponding vector of standardised residuals $\mathbf{Z} = (Z_1, \ldots, Z_D)$, where

$$Z_r = \frac{t_r(\mathbf{Y}) - M_r(\boldsymbol{\theta})}{S_r(\boldsymbol{\theta})}, \tag{21}$$

for $r = 1, \ldots, D$, where $M_r(\boldsymbol{\theta})$ and $S_r(\boldsymbol{\theta})$ denote respectively the mean and standard deviation of the marginal distribution of the statistic $t_r(\mathbf{Y})$, when the data array $\mathbf{Y}$ is drawn from the sampling distribution (which has density $p(\mathbf{y};\boldsymbol{\theta},\boldsymbol{u}(\mathbf{y}) = \mathbf{U}_0)$). We can now define the (squared Euclidean distance) discrepancy variable

$$D(\mathbf{Y};\boldsymbol{\theta}) = \sum_{r=1}^{D} {Z_r}^2.$$

We can not avoid some arbitrates in the choice of summary statistics to include in the vector $\boldsymbol{t}(\mathbf{y}) = (t_1(\mathbf{y}), \ldots, t_D(\mathbf{y}))$.

In the computation of the Monte Carlo estimate $\hat{P}(\mathbf{Y}_0;\mathbf{U}_0)$ (equation 20) of the posterior predictive p-value (equation 17), the *simulated* discrepancy $D\left(\mathbf{Y}_j;\mathbf{\Theta}_j\right)$ is computed from the *simulated* standardised residuals

$$Z_{j,r} = \frac{t_r\left(\mathbf{Y}_j\right) - M_r\left(\mathbf{\Theta}_j\right)}{S_r\left(\mathbf{\Theta}_j\right)}, \tag{22}$$

while the *observed* discrepancy $D\left(\mathbf{Y}_0;\mathbf{\Theta}_j\right)$ is computed from the *observed* standardised residuals

$$Z_{j,0,r} = \frac{t_r\left(\mathbf{Y}_0\right) - M_r\left(\mathbf{\Theta}_j\right)}{S_r\left(\mathbf{\Theta}_j\right)}, \tag{23}$$

for $r = 1, \ldots, D$.

### 2 Posterior predictive p-value estimates which are applicable to Approximate Bayesian Computations (ABC)

When we do not have a computationally useful formula for the likelihood function, we can resort to approximate Bayesian computation (ABC) methods Beaumont et al. (2002); Blum and François (2010); Csilléry et al. (2012) in order to sample form the posterior distribution. Whenever we lack such a formula for the likelihood function, we also lack a formula for the sampling distribution of the data (which depends on the likelihood function together with a normalising factor which depends on the data). In these situations we generally also lack computationally useful formulae for the marginal distributions of individual statistics $t_r\left(\mathbf{Y}\right)$ $(r = 1, \ldots, D)$, and for the moments of these marginal distributions (as functions of the parameter vector $\boldsymbol{\theta}$).

However, provided that we have a Monte Carlo sampler (for example, an ABC sampler) for the posterior density $\pi\left(\boldsymbol{\theta}\,|\mathbf{Y}_0\right)$, and a simulator for the model $M$, we can construct estimates of the posterior predictive p-value $P\left(\mathbf{Y}_0\right)$ (equation 17), from a stratified Monte Carlo sample, as follows. Using the Monte Carlo sampler, we first generate a sample of $N$ observation $\mathbf{\Theta}_j$ $(j = 1, \ldots, N)$ from the posterior density

$\pi(\boldsymbol{\theta}\,|\mathbf{Y}_0)$. For each observation $\boldsymbol{\Theta}_j$, we can then use the data simulator to generate a sub-sample of $R$ observations $\mathbf{Y}_{j,k}$ ($k = 1, \ldots, R$), drawn from the sampling density $p(\mathbf{y}; \boldsymbol{\Theta}_j, \boldsymbol{u}(\mathbf{y}) = \mathbf{U}_0)$. In this way we generate a stratified sample of observations $(\boldsymbol{\Theta}_j, \mathbf{Y}_{j,k})$ ($j = 1, \ldots, N$; $k = 1, \ldots, R$) from the joint posterior predictive distribution (equation 2).

For each observation $\boldsymbol{\Theta}_j$, we can compute the estimates

$$\hat{M}_r(\boldsymbol{\Theta}_j) = \frac{1}{R}\sum_{k=1}^{R} t_r(\mathbf{Y}_{j,k}), \tag{24}$$

and

$$\left(\hat{S}_r(\boldsymbol{\Theta}_j)\right)^2 = \frac{1}{R}\sum_{k=1}^{R}\left(t_r(\mathbf{Y}_{j,k}) - \hat{M}_r(\boldsymbol{\Theta}_j)\right)^2, \tag{25}$$

for $r = 1, \ldots, D$. So, for each observation $\boldsymbol{\Theta}_j$, we have a vector of estimated means $\hat{\boldsymbol{M}}(\boldsymbol{\Theta}_j) = \left(\hat{M}_1(\boldsymbol{\Theta}_j), \ldots, \hat{M}_D(\boldsymbol{\Theta}_j)\right)$, and a vector of estimated standard deviations $\hat{\boldsymbol{S}}(\boldsymbol{\Theta}_j) = \left(\hat{S}_1(\boldsymbol{\Theta}_j), \ldots, \hat{S}_D(\boldsymbol{\Theta}_j)\right)$.

Now, for each of observations $(\boldsymbol{\Theta}_j, \mathbf{Y}_{j,k})$ we can compute the (estimated) *simulated* standardised residuals

$$\hat{Z}_{j,k,r} = \frac{t_r(\mathbf{Y}_{j,k}) - \hat{M}_r(\boldsymbol{\Theta}_j)}{\hat{S}_r(\boldsymbol{\Theta}_j)}, \tag{26}$$

and the (estimated) *observed* standardised residuals

$$\hat{Z}_{j,0,r} = \frac{t_r(\mathbf{Y}_0) - \hat{M}_r(\boldsymbol{\Theta}_j)}{\hat{S}_r(\boldsymbol{\Theta}_j)}, \tag{27}$$

for $r = 1, \ldots, D$.

For each of observations $(\boldsymbol{\Theta}_j, \mathbf{Y}_{j,k})$ we can compute the (estimated) *simulated* discrepancy

$$\hat{D}(\mathbf{Y}_{j,k}; \boldsymbol{\Theta}_j) = \sum_{r=1}^{D} \hat{Z}^2_{j,k,r}, \tag{28}$$

and the (estimated) *observed* discrepancy

$$\hat{D}\left(\mathbf{Y}_0;\mathbf{\Theta}_j\right)=\sum_{r=1}^{D}\hat{Z}_{j,0,r}^2. \tag{29}$$

For each of observations $\mathbf{\Theta}_j$ from the posterior density $\pi\left(\boldsymbol{\theta}\,|\mathbf{Y}_0\right)$, the proportion of observations in the Monte Carlo sample which satisfy the condition $D\left(\mathbf{Y}_j;\mathbf{\Theta}_j\right)>D\left(\mathbf{Y}_0;\mathbf{\Theta}_j\right)$, is a Monte Carlo estimate

$$\hat{P}\left(\mathbf{Y}_0;\mathbf{\Theta}_j,\mathbf{U}_0\right)=\frac{1}{R}\sum_{k=1}^{R}\mathbb{I}\left\{\hat{D}\left(\mathbf{Y}_{j,k};\mathbf{\Theta}_j\right)>\hat{D}\left(\mathbf{Y}_0;\mathbf{\Theta}_j\right)\right\}, \tag{30}$$

of the posterior predictive p-value $P\left(\mathbf{Y}_0;\mathbf{\Theta}_j,\mathbf{U}_0\right)$ (equation 16), and

$$\begin{aligned}\hat{P}\left(\mathbf{Y}_0;\mathbf{U}_0\right)&=\frac{1}{N}\sum_{j=1}^{N}\hat{P}\left(\mathbf{Y}_0;\mathbf{\Theta}_j,\mathbf{U}_0\right)\\&=\frac{1}{NR}\sum_{j=1}^{N}\sum_{k=1}^{R}\mathbb{I}\left\{\hat{D}\left(\mathbf{Y}_{j,k};\mathbf{\Theta}_j\right)>\hat{D}\left(\mathbf{Y}_0;\mathbf{\Theta}_j\right)\right\},\end{aligned} \tag{31}$$

is a Monte Carlo estimate of the posterior predictive p-value $P\left(\mathbf{Y}_0;\mathbf{U}_0\right)$ (equation 17). (As before, $\mathbb{I}\{s\}$ is an indicator function which takes the value 1 when the condition $s$ is satisfied, and the value 0 otherwise.) See Algorithm 1 for an outline of a procedure for computing this estimate of the posterior predictive p-value $P\left(\mathbf{Y}_0;\mathbf{U}_0\right)$.

We end this section by considering the computational cost of computing the estimate $\hat{P}\left(\mathbf{Y}_0;\mathbf{U}_0\right)$ (equation 31) of the posterior predictive p-value $P\left(\mathbf{Y}_0;\mathbf{U}_0\right)$. We ignore for now the costs (which may be considerable) associated with generating the sample of $N$ observations from the posterior density $\pi\left(\boldsymbol{\theta}\,|\mathbf{Y}_0\right)$ (or an approximation to this the posterior density). Of the remaining computational costs, the most substantial is typically the time cost of performing the large number of simulations.

In order to obtain an accurate estimate $\hat{P}\left(\mathbf{Y}_0;\mathbf{U}_0\right)$ (equation 31) of the posterior predictive p-value $P\left(\mathbf{Y}_0;\mathbf{U}_0\right)$, we need to sample on the order of $N=1000$ observations from the posterior density $\pi\left(\boldsymbol{\theta}\,|\mathbf{Y}_0\right)$. For each of these observations $\mathbf{\Theta}_j$, we also need to obtain accurate estimates of the parameters ($\hat{\boldsymbol{M}}\left(\mathbf{\Theta}_j\right)$, $\hat{\boldsymbol{S}}\left(\mathbf{\Theta}_j\right)$) of the conditional sampling density $p\left(\boldsymbol{t};\mathbf{\Theta}_j,\boldsymbol{u}\left(\mathbf{y}\right)=\mathbf{U}_0\right)$ of the vector $\boldsymbol{t}=\boldsymbol{t}\left(\mathbf{y}\right)$. Therefore it is

**Algorithm 1** PPC for ABC

**for** $j$ in 1:$N$ **do**
  Sample $\boldsymbol{\Theta}_j \sim \pi\left(\boldsymbol{\theta} \,|\mathbf{Y}_0\right)$
  **for** $k$ in 1:$R$ **do**
    Simulate data array $\mathbf{Y}_{j,k} \sim p\left(\mathbf{y}; \boldsymbol{\Theta}_j, \boldsymbol{u}\left(\mathbf{y}\right) = \mathbf{U}_0\right)$
    (which satisfies condition $\boldsymbol{u}\left(\mathbf{Y}_{j,k}\right) = \mathbf{U}_0$);
    Compute vector of summary statistics $\mathbf{T}_{j,k} = \boldsymbol{t}\left(\mathbf{Y}_{j,k}\right)$;
  **end for**
  From the sample of vectors $\mathbf{T}_{j,k}$ ($k = 1, \ldots, R$):
  Compute vector of estimated means $\hat{\boldsymbol{M}}\left(\boldsymbol{\Theta}_j\right)$ of summary statistics;
  Compute vector of estimated standard deviations $\hat{\boldsymbol{S}}\left(\boldsymbol{\Theta}_j\right)$
  of summary statistics;
  Compute the *observed* discrepancy $\hat{D}\left(\mathbf{Y}_0; \boldsymbol{\Theta}_j\right)$;
  **for** $k$ in 1:$R$ **do**
    Compute the *simulated* discrepancy $\hat{D}\left(\mathbf{Y}_{j,k}; \boldsymbol{\Theta}_j\right)$;
  **end for**
**end for**
Compute posterior predictive p-value estimate $P\left(\mathbf{Y}_0; \mathbf{U}_0\right)$
using equation 31;

necessary to performed on the order of $R = 1{,}000$ simulations for each observation $\boldsymbol{\Theta}_j$.

It is possible to perform multiple simulations in parallel on a computing cluster. (Each simulation can be run independently.) If $N_{\text{parallel}}$ denotes an average of the number of simulations that run in parallel on a cluster, at any one time, then an estimate of the time cost of generating a sample of $NR$ observations from the *joint posterior predictive distribution*, is

$$C_{\text{PPC}} = NR\,\frac{c_{\text{sim}}}{N_{\text{parallel}}} \tag{32}$$

where $c_{\text{sim}}$ is the time cost of completing one simulation.

### 3 Approximate Bayesian computation (ABC)

In principle, we could sample from the posterior density $\pi\left(\boldsymbol{\theta} \,|\mathbf{Y}_0\right)$ by using an MCMC (Markov chain Monte Carlo) sampler (Berthier et al., 2002; Beaumont, 2003). However, in many applications we do not have a computationally useful formula for the likelihood function, and as a consequence standard MCMC methods are not applicable.

An alternative approach is to first generate a very large sample from the joint density $\pi(\boldsymbol{\theta}, \mathbf{y})$ of the parameter vector $\boldsymbol{\theta}$ and the data array $\mathbf{y}$, and then to construct from this input a sample from an approximation to the posterior (conditional) density $\pi(\boldsymbol{\theta} \,|\mathbf{Y}_0)$. The methods available for constructing this conditional sample include rejection sampling, and various regression methods, and more usually a combination of rejection sampling and regression. These methods are often referred to collectively as ABC (approximate Bayesian computation). The more descriptive term *likelihood-free* Bayesian computation is also used. Pritchard et al. (1999) first used a rejection sampling method for Bayesian computation (estimation of marginal posterior densities), which combined the use of the prior as the proposal distribution for the parameter values, with an acceptance region defined in terms of a distance which is computed from summary statistics. The earlier use of a rejection sampling method for Bayesian computation by Tavaré et al. (1997) was not strictly likelihood-free (and for this reason its applicability was limited to relatively simple models). The first ABC regression method was developed by Beaumont et al. (2002). Other likelihood-free Bayesian computation methods have also been developed, including ABC-MCMC (Marjoram et al., 2003; Wegmann et al., 2009), and ABC-PRC Sisson et al. (2007). Methods of this type are appropriate for Bayesian inference problems where it is relatively easy to generate simulated data under the statistical model, while it computationally costly to compute the likelihood function.

In order to describe the ABC method, we need to introduce some additional notation. Let $\boldsymbol{\theta}$ denote a vector of parameters for our model. These are the parameters which we are treating as unobserved (their values are uncertain). Let $\boldsymbol{t} = (t_1, \ldots, t_D) = \boldsymbol{t}(\mathbf{y})$ denote a vector of summary statistics $(t_1(\mathbf{y}), \ldots, t_D(\mathbf{y}))$, which can be computed from the data. From now on, we need to distinguish between the observed data set, and various simulated data sets. Let $\mathbf{T}_0 = \boldsymbol{t}(\mathbf{Y}_0)$ denote the vector of summary statistics computed from the observed data set, and let $\mathbf{T}_i = \boldsymbol{t}(\mathbf{Y}_i)$ denote the vector of summary statistics computed from the $i$th simulated data set.

In ABC we perform a large number of iterations $N_{\text{sim}}$ of a rejection sampling procedure. These iterations can be performed in parallel on a computing cluster. Each iteration $i$ begins with a proposal step in which a vector of parameter values $\mathbf{\Theta}_i$ is drawn from the joint prior distribution. This vector of parameter values is then passed to a data simulation algorithm. The data simulation algorithm generates a simulated data set, from which we compute the vector of summary statistics $\mathbf{T}_i$. We now have an observation $(\mathbf{\Theta}_i, \mathbf{T}_i)$ which is drawn from the joint prior distribution of the parameter vector $\boldsymbol{\theta}$ and the vector $\boldsymbol{t}$ of summary statistics. This is followed by an acceptance/rejection step, in which we compute the Euclidean distance $d_i = d(\mathbf{T}_0, \mathbf{T}_i)$ to determine if the vector $\mathbf{T}_i$ lies within a ball $B(\mathbf{T}_0; \delta)$ with centre $\mathbf{T}_0$ and radius $\delta$. This ball is the acceptance region. Only observations $(\mathbf{\Theta}_i, \mathbf{T}_i)$ for which the vector $\mathbf{T}_i$ lies within a ball $B(\mathbf{T}_0; \delta)$ are accepted.

In practice, rather than specifying the radius $\delta$ of the ball in advance, it is usually more convenient to specify the required number of observations $N$, in the accepted sample. If we rank the $N_{\text{sim}}$ observations from the proposal distribution with respect to the the Euclidean distance $d_i = d(\mathbf{T}_0, \mathbf{T}_i)$, we can then retain the $N$ having the shortest Euclidean distances. The proportion

$$P_\delta = \frac{N}{N_{\text{sim}}},$$

of iterations for which are accepted determines the radius $\delta$ the acceptance region $B(\mathbf{T}_0; \delta)$ (also referred to as the *tolerance*).

We number the accepted observations $i = 1, 2, ..., N$, and let $(\mathbf{\Theta}_i^*, \mathbf{T}_i^*)$ denote the $i$th accepted observation. The sequence of parameter vectors $\mathbf{\Theta}_i^*$ $(i = 1, 2, ..., N)$ from the accepted observations, constitute a sample from a first approximation to the posterior distribution of the parameter $\boldsymbol{\theta}$. As also pointed out by Beaumont et al. (2002), it is possible to improve on this approximation, by using a regression method to adjust the parameter vectors $\mathbf{\Theta}_i^*$ according to the displacement of the vector $\mathbf{T}_i^*$ from the observed vector of summary statistics $\mathbf{T}_0$.

The ABC rejection method outlined above involves two approximations, in addition to the inevitable Monte Carlo error of any simulation-based approach. First, since the observed data $\mathbf{Y}_0$, only enters the algorithm via the vector of summary statistics $\mathbf{T}_0 = \boldsymbol{t}(\mathbf{Y}_0)$, we can only hope to compute the conditional density $\pi(\boldsymbol{\theta}\,|\boldsymbol{t} = \mathbf{T}_0)$, rather than the true posterior density of the parameter $\boldsymbol{\theta}$ (unless the vector of summary statistics $\boldsymbol{t}$ is a *sufficient* statistic for the parameter $\boldsymbol{\theta}$). The second approximation is a consequence of the acceptance region extending beyond the point $\boldsymbol{t} = \mathbf{T}_0$. Using the ABC rejection method, we compute the conditional density $\pi(\boldsymbol{\theta}\,|\boldsymbol{t} \in B(\mathbf{T}_0; \delta))$, rather than the conditional density $\pi(\boldsymbol{\theta}\,|\boldsymbol{t} = \mathbf{T}_0)$. It is the error resulting from this second approximation which we hope to reduce by using an ABC regression method.

The insight behind these ABC regression methods (Beaumont et al., 2002; Blum and François, 2010) is that the sequence of parameter vectors $\boldsymbol{\Theta}_i^*$ ($i = 1, 2, ..., N$) from the accepted observations, can be represented in the form

$$\boldsymbol{\Theta}_i^* = \boldsymbol{B}\mathbf{T}_i^* + \mathbf{Z}_i, \tag{33}$$

where $\boldsymbol{B}$ is a matrix of coefficients, and each $\mathbf{Z}_i$ is a vector of residuals. Furthermore, as a first approximation, we can assume that the residuals are drawn from a distribution which does not depend on the vector of summary statistics $\mathbf{T}_i^*$. This is a linear model. An observation from the conditional density $\pi(\boldsymbol{\theta}\,|\boldsymbol{t} = \mathbf{T}_0)$ could therefore be represented in the same way, as

$$\boldsymbol{\Theta}_{i,0}^* = \boldsymbol{B}\mathbf{T}_0 + \mathbf{Z}_i. \tag{34}$$

Now, from the sequence of accepted observations $(\boldsymbol{\Theta}_i^*, \mathbf{T}_i^*)$ ($i = 1, 2, ..., N$), we can obtain a point estimate $\hat{\boldsymbol{B}}$ of the matrix of coefficients (using least squares regression, or an alternative method). We can now compute the vectors of empirical residuals

$$\hat{\mathbf{Z}}_i = \boldsymbol{\Theta}_i^* - \hat{\boldsymbol{B}}\mathbf{T}_i^*, \tag{35}$$

for $i = 1, 2, ..., N$. The $N$ vectors of empirical residuals form a matrix $\hat{\mathbf{Z}}$ of empirical residuals. From this sample of residual vectors, we could estimate the distribution of residuals. Alternatively, we could use the sample of empirical residual vectors $\hat{\mathbf{Z}}_i$

directly. The resulting predicted value of the parameter vectors

$$\begin{aligned}\boldsymbol{\Theta}^*_{i,0} &= \hat{\boldsymbol{B}}\mathbf{T}_0 + \hat{\mathbf{Z}}_i \\ &= \boldsymbol{\Theta}^*_i + \hat{\boldsymbol{B}}\left(\mathbf{T}_0 - \mathbf{T}^*_i\right), \end{aligned} \tag{36}$$

should more closely approximate a sample from the conditional density $\pi\left(\boldsymbol{\theta}\left|\boldsymbol{t} = \mathbf{T}_0\right.\right)$.

In Beaumont et al. (2002) the regression method employed was in fact local linear regression. In local linear regression each accepted observation $(\boldsymbol{\Theta}^*_i, \mathbf{T}^*_i)$ is assigned a weight $W_i = w\left(d_i\right)$, where $w\left(\cdot\right)$ a monotonically decreasing function of the Euclidean distances $d_i = d\left(\mathbf{T}_0, \mathbf{T}_i\right)$. The regression coefficients and the residuals are then estimated using weighted least squares regression.

Ordinary least squares regression and local linear (weighted least squares) regression methods can fail for various reasons when the number of predictor variables (summary statistics in the case of ABC) is increased. The inclusion in the regression model of predictor variables which are only weakly correlated (or uncorrelated) with the response variables (the parameters in the case of ABC) can inflate the sampling variance of the predicted values, resulting in less accurate predictions. There is an additional problem which arises from the reliance of ABC regression methods on an acceptance region which is defined by a Euclidean distance. The inclusion in the Euclidean distance of summary statistic which are only weakly correlated (or uncorrelated) with the model parameters, can contribute to erroneous inclusion or exclusion of observations within the acceptance region. More generally, if a weighted regression method is used, the inclusion of summary statistic which are only weakly correlated (or uncorrelated) with the model parameters (in the Euclidean distance from which the weights are calculated), can lead to anomalous weightings of the observations in the regression calculations.

More advanced regression methods (Wegmann et al., 2009; Blum, 2010; Blum and François, 2010; Csilléry et al., 2012) can also be used to approximate the conditional density $\pi\left(\boldsymbol{\theta}\left|\boldsymbol{t} = \mathbf{T}_0\right.\right)$. Wegmann et al. (2009) recommended partial least squares regression (PLS regression) (Wold, 1966), which these authors incorporated into a likelihood-free MCMC algorithm (ABC-MCMC). The R package *abc* (Csilléry et al., 2012) includes the option of ridge regression (Hoerl, 1962), and a neural network method, along with local linear regression.

In many applications of ABC (rejection and regression) methods, the main computational cost is the time cost of performing the large number of simulations required. In order to generate a sample of $N$ = 1000 observations from the (approximate) posterior distribution, it is necessary to performed on the order of $N_{\text{sim}}$ = 100,000 (or perhaps 1,000,000) simulations. As already mentioned, it is possible to perform multiple simulations in parallel on a computing cluster. (Each simulation can be run independently.) If $N_{\text{parallel}}$ denotes an average of the number of simulations that run in parallel on a cluster, at any one time, then an estimate of the time cost of generating a sample of $N$ observations from the (approximate) posterior distribution, is

$$C_{\text{ABC}} = N_{\text{sim}} \, \frac{c_{\text{sim}}}{N_{\text{parallel}}} \tag{37}$$

where $c_{\text{sim}}$ is the time cost of completing one simulation.

### 4 Using sequential ABC steps to sample from posteriors generated by data from multiple donors

The ABC regression method (outlined in Section 3) allows us to sample form an *approximate* posterior distribution generated from a vector of summary statistics $\mathbf{T}_0 = \boldsymbol{t}(\mathbf{Y}_0)$, computed from the observed data array $\mathbf{Y}_0$. As long as the regression methods employed in the ABC require appreciable data compression (from a data array $\mathbf{Y}_0$ with a large number of components, to a relatively low dimensional vector of summary statistics $\mathbf{T}_0 = \boldsymbol{t}(\mathbf{Y}_0)$), this would appear to limit the application of ABC regression methods to any problem where much of the information available in the data can not be captured by a relatively low dimensional vector of summary statistics. At the very least we might expect these ABC methods to waste much of the information provided by these data sets, and thus to relinquish the usual claim that Bayesian methods can make use of all the available information.

However, there are situations where it is possible to sample form approximate posterior distributions generated from multiple vectors of summary statistics, by applying the ABC regression method sequentially, and feeding-in the vectors of summary statistics one-at-a-time. Suppose that we have observed data $\mathbf{Y}_0 = \left(\mathbf{Y}_0^{(1)}, \ldots, \mathbf{Y}_0^{(m)}\right)$ from multiple individuals ($i = 1, \ldots, m$). If we can make the modelling assumption that (conditional on the unobserved values of the model parameters) the data arrays from individuals $i = 1, \cdots, m$, are statistically independent, then in principle we can perform the desired Bayesian computation by performing a sequence of Bayesian computations in which the individual-specific data arrays, $\mathbf{Y}_0^{(1)}, \ldots, \mathbf{Y}_0^{(m)}$, are feed-in one-at-a-time. This is the type of situation where we can generate a sample from the approximate posterior distribution by performing a sequence of ABC regression steps in which the individual-specific vectors of summary statistics, $\mathbf{T}_0^{(1)} = \boldsymbol{t}\left(\mathbf{Y}_0^{(1)}\right), \ldots, \mathbf{T}_0^{(m)} = \boldsymbol{t}\left(\mathbf{Y}_0^{(m)}\right)$, are fed-in one-at-a-time.

We now make these modelling assumptions more precise. The joint sampling distribution of the sequence of data arrays, $\mathbf{y}^{(1)}, \ldots, \mathbf{y}^{(m)}$ (from individuals $i = 1, \cdots, m$), is assumed to have a density of the form

$$p\left(\mathbf{y}^{(1)}, \ldots, \mathbf{y}^{(m)}; \boldsymbol{\theta}\right) = \prod_{i=1}^{m} p\left(\mathbf{y}^{(i)}; \boldsymbol{\theta}\right). \tag{38}$$

Here the data arrays from individuals $i = 1, \cdots, m$, are statistically independent, conditional on the parameter vector $\boldsymbol{\theta}$.

If the sequence of data arrays $\mathbf{y}^{(1)}, \ldots, \mathbf{y}^{(m)}$, has a joint sampling distribution with density of the form 38, then the *marginal* sampling density of the sequence of vectors $\boldsymbol{t}^{(1)}, \ldots, \boldsymbol{t}^{(m)}$, has a joint sampling distribution with density of the form

$$p\left(\boldsymbol{t}^{(1)},\ldots,\boldsymbol{t}^{(m)};\boldsymbol{\theta}\right)=\prod_{i=1}^{m}p\left(\boldsymbol{t}^{(i)};\boldsymbol{\theta}\right). \tag{39}$$

If the observed individual-specific vectors of summary statistics (computed from the observed data arrays) are $\mathbf{T}_0^{(1)}$, ... , $\mathbf{T}_0^{(m)}$, then the (multiple-individual) posterior density is

$$\begin{aligned}\pi^{*(m)}\left(\boldsymbol{\theta}\right)=&\pi\left(\boldsymbol{\theta}\middle|\mathbf{T}_0^{(1)},\ldots,\mathbf{T}_0^{(m)}\right)\\=&\frac{\left(\prod_{i=1}^{m}p\left(\mathbf{T}_0^{(i)};\boldsymbol{\theta}\right)\right)\pi\left(\boldsymbol{\theta}\right)}{\pi\left(\mathbf{T}_0^{(1)},\ldots,\mathbf{T}_0^{(m)}\right)},\end{aligned} \tag{40}$$

where

$$\pi\left(\boldsymbol{t}^{(1)},\ldots,\boldsymbol{t}^{(m)}\right)=\int_{\mathcal{R}}\left(\prod_{i=1}^{m}p\left(\boldsymbol{t}^{(i)};\boldsymbol{\theta}\right)\right)\pi\left(\boldsymbol{\theta}\right)\boldsymbol{d\theta}, \tag{41}$$

is the marginal density of the sequence of vectors $\boldsymbol{t}^{(1)}$, ... , $\boldsymbol{t}^{(m)}$.

Suppose that we only have the data array $\mathbf{Y}_0^{(1)}$, and the vector of summary statistics, $\mathbf{T}_0^{(1)}=\boldsymbol{t}\left(\mathbf{Y}_0^{(1)}\right)$, from the first individual in the sequence. The posterior density, conditional on the vector $\boldsymbol{t}^{(1)}$ (only) is given by

$$\begin{aligned}\pi^{*(1)}\left(\boldsymbol{\theta}\right)=&\pi\left(\boldsymbol{\theta}\middle|\mathbf{T}_0^{(1)}\right)\\=&\frac{p\left(\mathbf{T}_0^{(1)};\boldsymbol{\theta}\right)\pi\left(\boldsymbol{\theta}\right)}{\pi\left(\mathbf{T}_0^{(1)}\right)},\end{aligned} \tag{42}$$

where $\pi\left(\boldsymbol{t}^{(1)}\right)$ is the marginal density of the vector $\boldsymbol{t}^{(1)}$. It is possible to sample from (an approximation to) the posterior distribution $\pi^{*(1)}\left(\boldsymbol{\theta}\right)$ using an ABC rejection method, or an ABC regression method. In the case of the rejection method, at each iteration $j$, a parameter vector $\mathbf{\Theta}_j$ is sampled from the prior density $\pi(\boldsymbol{\theta})$. From the parameter vector $\mathbf{\Theta}_j$ we can generate a simulated data array $\mathbf{Y}_j^{(1)}$, which is drawn from the sampling density $p\left(\mathbf{y}^{(1)}; \mathbf{\Theta}_j\right)$. From the simulated data array $\mathbf{Y}_j^{(1)}$, we compute the vector of summary statistics $\mathbf{T}_j^{(1)} = \boldsymbol{t}\left(\mathbf{Y}_j^{(1)}\right)$. (The vector $\mathbf{T}_j^{(1)}$ is drawn from the sampling density $p\left(\boldsymbol{t}^{(1)}; \mathbf{\Theta}_j\right)$.) We now have an observation $\left(\mathbf{\Theta}_j, \mathbf{T}_j^{(1)}\right)$ which is drawn from the joint density

$$\pi^{(1)}\left(\boldsymbol{\theta}, \boldsymbol{t}^{(1)}\right) = p\left(\boldsymbol{t}^{(1)}; \boldsymbol{\theta}\right)\pi(\boldsymbol{\theta}).$$

This is the proposal density in our rejection sampling procedure. We compute the Euclidean distance $d_j = d\left(\mathbf{T}_0^{(1)}, \mathbf{T}_j^{(1)}\right)$ to determine if the vector $\mathbf{T}_j^{(1)}$ lies within a ball $B\left(\mathbf{T}_0^{(1)}; \delta\right)$ with centre $\mathbf{T}_0^{(1)}$ and radius $\delta$. Only observations $\left(\mathbf{\Theta}_j, \mathbf{T}_j^{(1)}\right)$ for which the vector $\mathbf{T}_j^{(1)}$ lies within this ball are accepted. The parameter vector from each *accepted* observation is then an independent draw from the conditional density $\pi\left(\boldsymbol{\theta} \left| \boldsymbol{t}^{(1)} \in B\left(\mathbf{T}_0^{(1)}; \delta\right)\right.\right)$, which is an approximation to the conditional density $\pi^{*(1)}(\boldsymbol{\theta}) = \pi\left(\boldsymbol{\theta} \left| \boldsymbol{t}^{(1)} = \mathbf{T}_0^{(1)}\right.\right)$.

Next, we will discover how to sample from the posterior density, conditional on any sub-sequence of observed vectors $\mathbf{T}_0^{(1)}, \ldots, \mathbf{T}_0^{(s)}$. Our starting point is equation 40, for the posterior density which is conditional on the sequence of vectors $\mathbf{T}_0^{(1)}, \ldots, \mathbf{T}_0^{(m)}$. It follows immediately from the definition of the conditional probability density $\pi\left(\boldsymbol{t}^{(m)} \left| \boldsymbol{t}^{(1)}, \ldots, \boldsymbol{t}^{(m-1)}\right.\right)$ that we can express the marginal density $\pi\left(\boldsymbol{t}^{(1)}, \ldots, \boldsymbol{t}^{(m)}\right)$, as

$$\pi\left(\boldsymbol{t}^{(1)}, \ldots, \boldsymbol{t}^{(m-1)}, \boldsymbol{t}^{(m)}\right) = \pi\left(\boldsymbol{t}^{(m)} \left| \boldsymbol{t}^{(1)}, \ldots, \boldsymbol{t}^{(m-1)}\right.\right)\pi\left(\boldsymbol{t}^{(1)}, \ldots, \boldsymbol{t}^{(m-1)}\right). \tag{43}$$

Making use of this fact, and the factorisation of the sampling distribution (equation 39, which is a consequence of the assumption of statistical independence), we can re-write equation 40 in the form

$$\begin{aligned}
\pi^{*(m)}(\boldsymbol{\theta}) =& \pi\left(\boldsymbol{\theta}\left|\mathbf{T}_0^{(1)},\ldots,\mathbf{T}_0^{(m)}\right.\right) \\
=& \frac{\left(\prod_{i=1}^{m} p\left(\mathbf{T}_0^{(i)};\boldsymbol{\theta}\right)\right)}{\pi\left(\mathbf{T}_0^{(1)},\ldots,\mathbf{T}_0^{(m)}\right)} \cdot \pi(\boldsymbol{\theta}) \\
=& \frac{p\left(\mathbf{T}_0^{(m)};\boldsymbol{\theta}\right)}{\pi\left(\mathbf{T}_0^{(m)}\left|\mathbf{T}_0^{(1)},\ldots,\mathbf{T}_0^{(m-1)}\right.\right)} \cdot \pi^{*(m-1)}(\boldsymbol{\theta}),
\end{aligned} \tag{44}$$

where

$$\begin{aligned}
\pi^{*(m-1)}(\boldsymbol{\theta}) =& \pi\left(\boldsymbol{\theta}\left|\mathbf{T}_0^{(1)},\ldots,\mathbf{T}_0^{(m-1)}\right.\right) \\
=& \frac{\left(\prod_{i=1}^{m-1} p\left(\mathbf{T}_0^{(i)};\boldsymbol{\theta}\right)\right)}{\pi\left(\mathbf{T}_0^{(1)},\ldots,\mathbf{T}_0^{(m-1)}\right)} \cdot \pi(\boldsymbol{\theta}).
\end{aligned} \tag{45}$$

If we define $\pi^{*(0)}(\boldsymbol{\theta}) = \pi(\boldsymbol{\theta})$, then the recursion 44 also holds true when $m = 1$. Notice that the equation 45 for the density $\pi^{*(m-1)}(\boldsymbol{\theta})$ is of exactly the same form as the equation 40 for the density $\pi^{*(m)}(\boldsymbol{\theta})$.

From the factorisation (equation 44) of the density $\pi^{*(m)}(\boldsymbol{\theta})$, it is apparent that if we can sample from the density $\pi^{*(m-1)}(\boldsymbol{\theta})$ (by whatever method is available), then we can also sample from the density $\pi^{*(m)}(\boldsymbol{\theta})$ using a rejection method (or a regression method), as follows. At each iteration $j$, a parameter vector $\boldsymbol{\Theta}_j$ is sampled from the *prior* density $\pi^{*(m-1)}(\boldsymbol{\theta})$. From the parameter vector $\boldsymbol{\Theta}_j$ we can generate a simulated data array $\mathbf{Y}_j^{(m)}$, which is drawn from the sampling density $p\left(\mathbf{y}^{(m)};\boldsymbol{\Theta}_j\right)$. From the simulated data array $\mathbf{Y}_j^{(m)}$, we compute the vector of summary statistics $\mathbf{T}_j^{(m)} = \boldsymbol{t}\left(\mathbf{Y}_j^{(m)}\right)$. (The vector $\mathbf{T}_j^{(m)}$ is drawn from the sampling density $p\left(\boldsymbol{t}^{(m)};\boldsymbol{\Theta}_j\right)$.) We now have an observation $\left(\boldsymbol{\Theta}_j, \mathbf{T}_j^{(m)}\right)$ which is drawn from the joint density

$$\pi^{(m)}\left(\boldsymbol{\theta}, \boldsymbol{t}^{(m)}\right) = p\left(\boldsymbol{t}^{(m)};\boldsymbol{\theta}\right)\pi^{*(m-1)}(\boldsymbol{\theta}).$$

This is the proposal density. We compute the Euclidean distance $d_j = d\left(\mathbf{T}_0^{(m)}, \mathbf{T}_j^{(m)}\right)$ to determine if the vector $\mathbf{T}_j^{(m)}$ lies within a ball $B\left(\mathbf{T}_0^{(m)}; \delta\right)$ with centre $\mathbf{T}_0^{(m)}$ and radius $\delta$. Only observations $\left(\boldsymbol{\Theta}_j, \mathbf{T}_j^{(m)}\right)$ for which the vector $\mathbf{T}_j^{(m)}$ lies within this ball are accepted. The parameter vector from each *accepted* observation is then an independent draw from a conditional density, which is an approximation to the conditional density $\pi^{*(m)}(\boldsymbol{\theta})$.

We have established that if we can sample from the density $\pi^{*(m-1)}(\boldsymbol{\theta})$ (by whatever method is available), then we can also sample from the density $\pi^{*(m)}(\boldsymbol{\theta})$, as claimed. We have already seen that we can sample from the density $\pi^{*(1)}(\boldsymbol{\theta})$. Therefore, we can sample from every density in the sequence $\pi^{*(1)}(\boldsymbol{\theta})$, $\pi^{*(2)}(\boldsymbol{\theta})$, ..., $\pi^{*(m-1)}(\boldsymbol{\theta})$, $\pi^{*(m)}(\boldsymbol{\theta})$. Furthermore, at each step $i$, in order to obtain a sample from the density $\pi^{*(i)}(\boldsymbol{\theta})$, we only have to condition on the data $\mathbf{T}_j^{(i)} = \boldsymbol{t}\left(\mathbf{Y}_j^{(i)}\right)$ from a single individual.

See Algorithm 2 for an outline of this sequential ABC sampling algorithm for generating a sample from the (approximate) posterior density $\pi^{*(m)}(\boldsymbol{\theta})$, which is conditional on the sequence of observed vectors $\mathbf{T}_0^{(1)}, \ldots, \mathbf{T}_0^{(m)}$. The validity of this sequential ABC sampling method depends crucially on the factorisation in equation 44 of the posterior density, and this factorisation of the posterior density is a consequence of the factorisation (equation 39) of the joint sampling distribution of the sequence of vectors $\boldsymbol{t}^{(1)}, \ldots, \boldsymbol{t}^{(m)}$. Recall that we introduced the assumption that (conditional on the values of the model parameters $\boldsymbol{\theta}$) the sequence of data arrays, $\mathbf{y}^{(1)}, \ldots, \mathbf{y}^{(m)}$ (from individuals $i = 1, \cdots, m$), are statistically independent (represented by the factorisation in 38). This implies the statistical independence (conditional on the values of the model parameters $\boldsymbol{\theta}$) of the sequence of vectors $\boldsymbol{t}^{(1)}, \ldots, \boldsymbol{t}^{(m)}$ (represented by the factorisation in 39).

Each step $i$ of the sequential ABC sampling algorithm (Algorithm 2) begins with sampling a large number $N_{\text{sim}}$ of observations $\boldsymbol{\Theta}_j^{(i)}$ ($j = 1, \ldots, N_{\text{sim}}$), from the *prior* $\pi^{*(i-1)}(\boldsymbol{\theta})$. At the first ABC step, the $N_{\text{sim}}$ observations $\boldsymbol{\Theta}_j^{(1)}$ are sampled directly from the prior density $\pi^{*(0)}(\boldsymbol{\theta}) = \pi(\boldsymbol{\theta})$. At each subsequent ABC step ($i = 2, \ldots, m$), we can generate a sample of $N_{\text{sim}}$ observations $\boldsymbol{\Theta}_j^{(i)}$, from the *prior* $\pi^{*(i-1)}(\boldsymbol{\theta})$, by sampling *with replacement* from the sample of $N$ *accepted* observations $\mathbf{\Theta}_j^{*(i-1)}$ ($j = 1$, …, $N$) obtained at the end of preceding ABC step.

There is a potential problem with this approach. If the ABC rejection method is used, the sample of $N$ *accepted* observations $\mathbf{\Theta}_j^{*(i)}$ ($j = 1, \ldots, N$) obtained at the end of step $i = 1, 2, \ldots, m$, can never contain any new values which were not already present in the original sample of $N_{\text{sim}}$ observations drawn from the prior density $\pi^{*(0)}(\boldsymbol{\theta}) = \pi(\boldsymbol{\theta})$. After many successive ABC rejection steps $m$), this can lead to depletion of the variation in the Monte Carlo sample $\mathbf{\Theta}_j^{*(m)}$ ($j = 1, \ldots, N$) from $\pi^{*(m)}(\boldsymbol{\theta})$.

In order to avoid this problem of variation depletion, we recommend using an ABC regression method at each ABC sampling step (rather than the ABC rejection method). When an ABC regression method is used, the $N$ regression adjusted values, $\mathbf{\Theta}_{j,0}^{*(i)}$ ($j = 1, \ldots, N$), in the sample obtained at the end of ABC sampling step $i$, are typically all new and distinct values. This is because the regression adjustment of each accepted observations $\mathbf{\Theta}_j^{*(i)}$ ($j = 1, \ldots, N$), depends on the vector $\mathbf{T}_j^{*(i)}$ computed from the simulated data array $\mathbf{Y}_j^{*(i)}$. Furthermore, these the regression adjustments depend on the point estimate $\hat{\boldsymbol{B}}$ of the matrix of coefficients, and the matrix $\hat{\mathbf{Z}}$ of empirical residuals, and these point estimates are determined by the sample of accepted observations $\left(\mathbf{\Theta}_j^{*(i)}, \mathbf{T}_j^{*(i)}\right)$, $j = 1, \ldots, N$.

There are also potential problems with the successive application of the ABC regression method to the output of the preceding ABC regression step. If the parameter vector $\boldsymbol{\theta}$ includes component parameters about which some of the vectors of summary statistics $\mathbf{T}_0^{(i)} = \boldsymbol{t}\left(\mathbf{Y}_0^{(i)}\right)$, provides very little information, then at each ABC regression step $i$, the corresponding components of the vectors of regression adjusted values $\mathbf{\Theta}_{j,0}^{*(i)}$ ($j = 1, \ldots, N$) may have inflated sampling variances. Therefore, in applications to models where this is an issue, the sequential ABC regression method outlined here may need to be modified. In our analysis of HSC whole genome sequence data from 8 donor individuals, it was found that the data from the younger individuals provides very little information about the parameters of the non-neural model. For this reason, we used the sequential ABC regression algorithm (Algorithm 2) to sample from the multiple-individual posterior density (equation 40), conditional on only the data from the 4 oldest donor individuals.

**Algorithm 2** Sequential ABC regression

**for** $i$ in 1:$m$ **do**
  **for** $j$ in 1:$N_{\text{sim}}$ **do**
    Sample from *prior* $\mathbf{\Theta}_j^{(i)} \sim \hat{\pi}^{*(i-1)}(\boldsymbol{\theta})$;
    Simulate data array $\mathbf{Y}_j^{(i)} \sim p\left(\mathbf{y}^{(i)}; \mathbf{\Theta}_j^{(i)}\right)$;
    Compute vector of summary statistics $\mathbf{T}_j^{(i)} = \boldsymbol{t}\left(\mathbf{Y}_j^{(i)}\right)$;
  **end for**
  Rank observations $\left(\mathbf{\Theta}_j^{(i)}, \mathbf{T}_j^{(i)}\right)$ $(j = 1, \ldots, N_{\text{sim}})$
  w.r.t. Euclidean distance $d_j = d\left(\mathbf{T}_0^{(i)}, \mathbf{T}_j^{(i)}\right)$;
  Accept the $N$ observations having the shortest distances $d_j = d\left(\mathbf{T}_0^{(i)}, \mathbf{T}_j^{(i)}\right)$;
  to obtain the accepted sample $\left(\mathbf{\Theta}_j^{*(i)}, \mathbf{T}_j^{*(i)}\right)$ $(j = 1, \ldots, N)$;
  Apply regression method to the accepted sample $\left(\mathbf{\Theta}_j^{*(i)}, \mathbf{T}_j^{*(i)}\right)$ $(j = 1, \ldots, N)$
  to obtain the adjusted values $\mathbf{\Theta}_{j,0}^{*(i)}$ $(j = 1, \ldots, N)$;
  The adjusted values $\mathbf{\Theta}_{j,0}^{*(i)}$ $(j = 1, \ldots, N)$ are a sample from $\hat{\pi}^{*(i)}(\boldsymbol{\theta})$;
**end for**

As mentioned in Section 3, in many applications of ABC (rejection and regression) methods, the main computational cost is the time cost of performing the large number of simulations required. In order to generate a sample of $N = 1000$ observations from the (approximate) posterior distribution, it is necessary to performed on the order of $N_{\text{sim}} = 100{,}000$ (or perhaps on the order of $N_{\text{sim}} = 1{,}000{,}000$) simulations. Fortunately, each simulation can be run independently, and it is therefore possible to perform multiple simulations in parallel on a computing cluster. If $N_{\text{parallel}}$ denotes an average of the number of simulations that can run in parallel on a cluster, at any one time, then an estimate of the time cost of generating a sample of $N$ observations from the (approximate) posterior distribution is $C_{\text{ABC}}$, given by equation 37.

However, the $m$ ABC regression steps in the sequential ABC sampling algorithm (Algorithm 2) must be performed in sequence, and can not be run in parallel. Therefore the total time cost of generating a sample of $N$ observations from the (approximate) posterior density $\pi^{*(m)}(\boldsymbol{\theta})$ is

$$C_{\text{ABC Seq}}(m) = m\, C_{\text{ABC}} = mN_{\text{sim}}\, \frac{c_{\text{sim}}}{N_{\text{parallel}}}. \tag{46}$$

This time cost, which increases linearly with the number $m$ of individuals in the sequence (and hence the number of ABC regression steps performed), is in contrast to the much lower time cost which can be achieved for the sequence of ABC sampling steps required generate samples from a sequence of individual-specific posterior densities, from the same sequence of individual-specific data arrays $\mathbf{Y}_0^{(i)}$.

For each individual $i$, the individual-specific posterior density is

$$\pi\left(\boldsymbol{\theta}\left|\mathbf{T}_0^{(i)}\right.\right)=\frac{p\left(\mathbf{T}_0^{(i)};\boldsymbol{\theta}\right)\pi\left(\boldsymbol{\theta}\right)}{\pi\left(\mathbf{T}_0^{(i)}\right)}. \tag{47}$$

This is the posterior density, conditional on the the individual-specific observed vector $\mathbf{T}_0^{(i)}$ (computed from the observed data array $\mathbf{Y}_0^{(i)}$) from individual $i$ (only). Here $\pi\left(\boldsymbol{t}^{(i)}\right)$ is the marginal density of the individual-specific vectors of summary statistics $\boldsymbol{t}^{(i)}$.

Notice that for each individual $i$, the prior used in the individual-specific Bayesian calculation (equation 47) is the same. We can therefore re-use the same sample of $N_{\text{sim}}$ parameter vectors $\boldsymbol{\Theta}_j$ (drawn from the prior $\pi\left(\boldsymbol{\theta}\right)$), and the same simulations, and hence the same sample of observations $\left(\boldsymbol{\Theta}_j,\mathbf{T}_j\right)$ ($j=1,\ldots,N_{\text{sim}}$) drawn from the proposal distribution. Therefore, the total time cost of generating $m$ separate samples, of $N$ observations, from each of the individual-specific posterior densities $\pi\left(\boldsymbol{\theta}\left|\mathbf{T}_0^{(i)}\right.\right)$ (for individuals $i=1,2,\ldots,m$), is $C_{\text{ABC}}$ (given by equation 37), and does not increase with the number $m$ of individuals. That is

$$C_{\text{ABC Sep}}\left(m\right)=C_{\text{ABC}}=N_{\text{sim}}\,\frac{c_{\text{sim}}}{N_{\text{parallel}}}. \tag{48}$$

In practice, the situation is often a little more complicated than this. In order to fully specify the sampling distribution from which a simulator for the model $M$ will sample, we need to specify the parameter vector $\boldsymbol{\theta}$, together with a vector of *auxiliary statistics*, $\mathbf{U}_0=\boldsymbol{u}\left(\mathbf{Y}_0\right)$, computed from the observed data array. These auxiliary statistics are the functions of the data array which are constrained to have the same values (for all data arrays drawn from the sampling distribution) as they have for the observed data array.

When we have a sequence of individual-specific data arrays, $\mathbf{Y}_0^{(1)}, \ldots, \mathbf{Y}_0^{(m)}$, in general, the vectors of auxiliary statistics, $\mathbf{U}_0^{(1)}, \ldots, \mathbf{U}_0^{(m)}$, will be different for each individual $i$ in the sequence. As a consequence of these differences in the auxiliary statistics, the simulations generated for use in an ABC sampler for individual-specific ABC posterior density (equation 47) for individual $i$, can not be re-used directly in an ABC sampler for the individual-specific ABC posterior density for some other individual.

Fortunately, in most cases, we can easily get around this problem by generating a single sample of $N_{\text{sim}}$ *master simulations*. From each of these master simulations, we can obtain a simulated data set satisfying any of the constraints represented by the vectors of auxiliary statistics, $\mathbf{U}_0^{(1)}, \ldots, \mathbf{U}_0^{(m)}$. To take a simple example, suppose that individuals 1, 2, …, $m$, have been given a questionnaire to complete, and that the data array $\mathbf{Y}_0^{(i)}$ contains the responses of individual $i$. Each observed data array $\mathbf{Y}_0^{(i)}$ may have some missing data, because the individual failed to record a response to some of the questions, and which data is missing may be different for different individuals. If we have a model for how individuals respond to this questionnaire, then we could in principle use a simulator to generate *master simulations* consisting of completed questionnaires. From these master simulations we can obtain individual-specific simulations for individual $i$, which satisfy the constraints represented by the vector of auxiliary statistics, $\mathbf{U}_0^{(i)}$, simply by censoring the data arrays from the master simulations.

In our analysis of HSC whole genome sequence data from 8 donor individuals, the individual-specific data array $\mathbf{Y}_0^{(i)}$ is a representation of a *sample phylogeny* on a sample of single cell genomes (together with mutation assignments to branches) from individual $i$. The vector of auxiliary statistics for individual $i$ is $\mathbf{U}_0^{(i)} = \left(n^{(i)}, a^{(i)}\right)$, where $n^{(i)}$ is the number of single cell genomes in the sample from individual $i$, and $a^{(i)}$ is the age of individual $i$. In general, the individuals 1, 2, …, $m$, differ in age, and in the number of single cell genomes sampled. Each simulation (performed using the *rsimpop* package, Williams et al. (2020)) returns a *population phylogeny*. Provided that the simulator is run for the age of the oldest donor individual in the sequence, the population phylogenies generated can be used as master simulations. From these master simulations we can obtain individual-specific simulations for individual $i$, which satisfy the constraints represented by $\mathbf{U}_0^{(i)} = \left(n^{(i)}, a^{(i)}\right)$, by terminating the population phylogeny at the age $a^{(i)}$, and deleting terminal nodes to obtain a sample phylogeny on exactly $n^{(i)}$ single cell genomes. By the use of an appropriate sample of master simulations, it will usually be possible to achieve a total time cost having the remarkably low magnitude indicated by equation 48 (and equation 37), for a sequence of ABC samplers, for individual-specific posterior densities.

### 5 Estimation of donor-specific posterior predictive p-values from posteriors generated by single donors and by multiple donors

As in Section 4, suppose that we have observed data $\mathbf{Y}_0 = \left(\mathbf{Y}_0^{(1)}, \ldots, \mathbf{Y}_0^{(m)}\right)$ from multiple individuals ($i = 1, \ldots, m$). In order to asses the compatibility of the proposed model with the observed data $\mathbf{Y}_0^{(i)}$, from a specific individual $i$, we want to compute the posterior predictive p-values $P\left(\mathbf{Y}_0^{(i)}; \mathbf{U}_0^{(i)}\right)$, for individual $i$.

For each individual $i$, we can define an individual-specific posterior predictive p-value $P\left(\mathbf{Y}_0^{(i)}; \mathbf{U}_0^{(i)}\right)$, based on the individual-specific posterior density

$$\pi\left(\boldsymbol{\theta} \middle| \mathbf{Y}_0^{(i)}\right) = \frac{p\left(\mathbf{Y}_0^{(i)}; \boldsymbol{\theta}\right) \pi\left(\boldsymbol{\theta}\right)}{\pi\left(\mathbf{Y}_0^{(i)}\right)}. \tag{49}$$

This is the posterior density, conditional on the observed individual-specific data array $\mathbf{Y}_0^{(i)}$ from individual $i$ (only). Here $\pi\left(\mathbf{y}^{(i)}\right)$ is the marginal density of the individual-specific data array $\mathbf{y}^{(i)}$. (Compare equation 49 with equation 47 for the posterior density conditional on a vector of summary statistics, rather than the full data array for individual $i$.)

The individual-specific posterior predictive p-value $P\left(\mathbf{Y}_0^{(i)}; \mathbf{U}_0^{(i)}\right)$, based on this individual-specific posterior density, is

$$P\left(\mathbf{Y}_0^{(i)}; \mathbf{U}_0\right) = \int_{\mathcal{R}} \int_{\mathbf{y}^{(i)}: D\left(\mathbf{y}^{(i)}; \boldsymbol{\theta}\right) > D\left(\mathbf{Y}_0^{(i)}; \boldsymbol{\theta}\right)} \pi\left(\boldsymbol{\theta}, \mathbf{y}^{(i)} \middle| \mathbf{Y}_0^{(i)}, \boldsymbol{u}\left(\mathbf{y}^{(i)}\right) = \mathbf{U}_0^{(i)}\right) \boldsymbol{d}\mathbf{y}^{(i)} \boldsymbol{d\theta}, \quad (50)$$

where

$$\pi\left(\boldsymbol{\theta}, \mathbf{y}^{(i)} \middle| \mathbf{Y}_0^{(i)}, \boldsymbol{u}\left(\mathbf{y}^{(i)}\right) = \mathbf{U}_0^{(i)}\right) = p\left(\mathbf{y}^{(i)}; \boldsymbol{\theta}, \boldsymbol{u}\left(\mathbf{y}^{(i)}\right) = \mathbf{U}_0^{(i)}\right) \pi\left(\boldsymbol{\theta} \middle| \mathbf{Y}_0^{(i)}\right). \quad (51)$$

The density $\pi\left(\boldsymbol{\theta}, \mathbf{y}^{(i)} \middle| \mathbf{Y}_0^{(i)}, \boldsymbol{u}\left(\mathbf{y}^{(i)}\right) = \mathbf{U}_0^{(i)}\right)$ is what we refer to as the *joint posterior predictive density.* The density $p\left(\mathbf{y}^{(i)}; \boldsymbol{\theta}, \boldsymbol{u}\left(\mathbf{y}^{(i)}\right) = \mathbf{U}_0^{(i)}\right)$ is the individual-specific conditional sampling density, and $\pi\left(\boldsymbol{\theta} \middle| \mathbf{Y}_0^{(i)}\right)$ is the individual-specific posterior density.

Recall that, in order to fully specify a posterior predictive density, and the joint posterior predictive density, it is also necessary to know which functions of the data array are constrained to have the same values for the new unobserved data array as they have for the observed data array. These functions of the data array are what refer to as *auxiliary statistics* (following Gelman et al. (1996)). Let $\mathbf{U}_0^{(i)} = \boldsymbol{u}\left(\mathbf{U}_0^{(i)}\right)$ denote the vector of auxiliary statistics computed from the observed data array $\mathbf{Y}_0^{(i)}$ from individual $i$.

If we can sample observations $\boldsymbol{\Theta}_j$ ($j = 1, \ldots, N$), from the individual-specific posterior density $\pi\left(\boldsymbol{\theta} \middle| \mathbf{Y}_0^{(i)}\right)$ (equation 49), then we can compute a Monte Carlo estimate of the individual-specific posterior predictive p-value $P\left(\mathbf{Y}_0^{(i)}; \mathbf{U}_0^{(i)}\right)$ defined by equation 50. In fact it is possible to sample from (an approximation to) the individual-specific posterior density $\pi\left(\boldsymbol{\theta} \middle| \mathbf{Y}_0^{(i)}\right)$ using an ABC rejection method, or an ABC regression method.

Recall from Section 2, that the time cost of computing one of these Monte Carlo estimate of a posterior predictive p-value is given by equation 32, where $N$ is the number of observations sampled from the (approximate) posterior density (in this case, the the individual-specific posterior density in equation 49), $R$ is the number of simulations performed for each observation, and $N_{\text{parallel}}$ denotes an average of the number of simulations that can run in parallel on a cluster, at any one time. Therefore the total time cost of estimating a sequence of $m$ of these posterior predictive p-value is

$$C_{\text{PPC Sep}}(m) = m\, C_{\text{PPC}} = mNR\, \frac{c_{\text{sim}}}{N_{\text{parallel}}}. \tag{52}$$

Recall that the total time cost of running $m$ ABC samplers, to generate $m$ separate samples of $N$ observations, from each of the individual-specific posterior densities $\pi\left(\boldsymbol{\theta}\,\middle|\,\mathbf{T}_0^{(i)}\right)$ (for individuals $i = 1, 2, \ldots, m$), is only $C_{\text{ABC Sep}}(m)$ (given by equation 48), and remarkably, this cost does not increase with the number $m$ of individuals. (See Section 4 for details.) Therefore, the total time cost of running $m$ ABC samplers, and then estimating a sequence of $m$ posterior predictive p-values, is

$$C_{\text{ABC Sep}}(m) + C_{\text{PPC Sep}}(m) = (N_{\text{sim}} + mNR)\, \frac{c_{\text{sim}}}{N_{\text{parallel}}}. \tag{53}$$

In our analysis of HSC genomic data from 8 donor individuals, we ran $m = 8$ separate ABC samplers, to generate samples of $N = 1000$ observations, from each of the 8 individual-specific posterior densities. All of these ABC samplers were able to use the same sample of $N_{\text{sim}} = 100{,}000$ simulations. The ABC rejection sampling method was used ($P_\delta = N/N_{\text{sim}} = 0.01$). For each of the $m = 8$ individual-specific posterior densities, we then estimated the individual-specific posterior predictive p-value, using the sample of $N = 1000$ accepted observations generated by the ABC sampler, and $R = 1000$ re-simulations for each accepted observation, making a total of $NR = 1{,}000{,}000$ simulations (for each individual donor). Therefore, using the estimate from equation 53, the total time cost of running $m$ ABC samplers, and then estimating a sequence of $m$ individual-specific posterior predictive p-values (each one based on the corresponding individual-specific posterior density), is of order

$$
\begin{aligned}
C_{\text{ABC Sep}}(m) + C_{\text{PPC Sep}}(m) &= \left(10^5 + 8 \times 10^6\right) \frac{c_{\text{sim}}}{N_{\text{parallel}}} \\
&\approx 8 \times 10^6 \frac{c_{\text{sim}}}{N_{\text{parallel}}}. \qquad (54)
\end{aligned}
$$

For each individual $i$, we can also define an individual-specific posterior predictive p-value $P\left(\mathbf{Y}_0^{(i)}; \mathbf{U}_0^{(i)}\right)$, based on the multiple-individual posterior density

$$
\begin{aligned}
\pi^{*(s)}(\boldsymbol{\theta}) =& \pi\left(\boldsymbol{\theta} \middle| \mathbf{Y}_0^{(1)}, \ldots, \mathbf{Y}_0^{(s)}\right) \\
=& \frac{\left(\prod_{i=1}^{s} p\left(\mathbf{Y}_0^{(i)}; \boldsymbol{\theta}\right)\right) \pi(\boldsymbol{\theta})}{\pi\left(\mathbf{Y}_0^{(1)}, \ldots, \mathbf{Y}_0^{(s)}\right)}. \qquad (55)
\end{aligned}
$$

This is the multiple-individual posterior density, conditional on the sequence of data arrays $\mathbf{Y}_0^{(1)}$, ..., $\mathbf{Y}_0^{(s)}$. Here $\pi\left(\mathbf{y}^{(1)}, \ldots, \mathbf{y}^{(s)}\right)$ is the marginal density of the sequence of data arrays $\mathbf{y}^{(1)}$, ..., $\mathbf{y}^{(s)}$. (Compare equation 55 with equation 40 for the posterior density conditional on a vector of summary statistics, rather than the full data array for individual $i$.)

The individual-specific posterior predictive p-value $P\left(\mathbf{Y}_0^{(i)}; \mathbf{U}_0^{(i)}\right)$, based on this multiple-individual posterior density (equation 55), is

$$
\begin{aligned}
&P\left(\mathbf{Y}_0^{(i)}; \mathbf{U}_0^{(i)}\right) \\
&= \int_{\mathcal{R}} \int_{\mathbf{y}^{(i)}: D\left(\mathbf{y}^{(i)}; \boldsymbol{\theta}\right) > D\left(\mathbf{Y}_0^{(i)}; \boldsymbol{\theta}\right)} \pi\left(\boldsymbol{\theta}, \mathbf{y}^{(i)} \middle| \mathbf{Y}_0^{(1)}, \ldots, \mathbf{Y}_0^{(s)}, \boldsymbol{u}\left(\mathbf{y}^{(i)}\right) = \mathbf{U}_0^{(i)}\right) \boldsymbol{d}\mathbf{y}^{(i)} \boldsymbol{d\theta}, \qquad (56)
\end{aligned}
$$

where

$$\begin{aligned}&\pi\left(\boldsymbol{\theta},\mathbf{y}^{(i)}\middle|\mathbf{Y}_0^{(1)},\ldots,\mathbf{Y}_0^{(s)},\boldsymbol{u}\left(\mathbf{y}^{(i)}\right)=\mathbf{U}_0^{(i)}\right)\\&=p\left(\mathbf{y}^{(i)};\boldsymbol{\theta},\boldsymbol{u}\left(\mathbf{y}^{(i)}\right)=\mathbf{U}_0^{(i)}\right)\pi\left(\boldsymbol{\theta}\middle|\mathbf{Y}_0^{(1)},\ldots,\mathbf{Y}_0^{(s)}\right).\end{aligned}\tag{57}$$

The density $\pi\left(\boldsymbol{\theta},\mathbf{y}^{(i)}\middle|\mathbf{Y}_0^{(1)},\ldots,\mathbf{Y}_0^{(s)},\boldsymbol{u}\left(\mathbf{y}^{(i)}\right)=\mathbf{U}_0^{(i)}\right)$, defined in equation 57, is what we refer to as the *joint posterior predictive density.* The density $p\left(\mathbf{y}^{(i)};\boldsymbol{\theta},\boldsymbol{u}\left(\mathbf{y}^{(i)}\right)=\mathbf{U}_0^{(i)}\right)$ is the individual-specific conditional sampling density, and $\pi^{*(s)}\left(\boldsymbol{\theta}\right)=\pi\left(\boldsymbol{\theta}\middle|\mathbf{Y}_0^{(1)},\ldots,\mathbf{Y}_0^{(s)}\right)$ is the multiple-individual posterior density (as defined in equation 55).

It is possible to sample from (an approximation to) the multiple-individual posterior density $\pi^{*(s)}\left(\boldsymbol{\theta}\right)=\pi\left(\boldsymbol{\theta}\middle|\mathbf{Y}_0^{(1)},\ldots,\mathbf{Y}_0^{(s)}\right)$ using the sequential ABC regression method (Algorithm 2) described in Section 4. Recall that the total time cost of generating a sample of $N$ observations from the (approximate) posterior density $\pi^{*(m)}\left(\boldsymbol{\theta}\right)$, using the sequential ABC regression method, is $C_{\text{ABC Seq}}\left(m\right)$ (given by equation 46).

In order to compute a Monte Carlo estimate of an individual-specific posterior predictive p-value $P\left(\mathbf{Y}_0^{(i)};\mathbf{U}_0^{(i)}\right)$ (for individual $i$, given by equation 56), based on the multiple-individual posterior density $\pi^{*(s)}\left(\boldsymbol{\theta}\right)$, we need to perform $R$ simulations for each the $N$ observations in the sample from the multiple-individual posterior density $\pi^{*(s)}\left(\boldsymbol{\theta}\right)$.

Notice that for each for each individual $i$, the individual-specific posterior predictive p-value defined in equation 56, is obtained by averaging over the same multiple-individual posterior density $\pi^{*(s)}\left(\boldsymbol{\theta}\right)$ (equation 55). Therefore, we can re-use the same sample of $N$ observations from the multiple-individual posterior density $\pi^{*(s)}\left(\boldsymbol{\theta}\right)$ when we compute our Monte Carlo estimates of the individual-specific p-values, for individuals $i=1,\,2,\,\ldots,\,m$. In principle we can also re-use the same simulations, and the same simulations, and hence the same sample of $NR$ observations $\left(\boldsymbol{\Theta}_j,\mathbf{Y}_{j,k}\right)$ (for draws $j=1,\,\ldots,\,N$, and re-simulation $k=1,\,\ldots,\,R$) drawn from the joint posterior predictive density (equation 57). The insight is exploited in the Algorithm 3.

When the simulations are re-used in this way, the total time cost of computing a sequence of $m$ individual-specific posterior predictive p-value (defined in equation 56), is $C_{\text{PPC}}$ (given by equation 32), and does not increase with the number $m$ of individuals. That is

$$C_{\text{PPC Seq}}(m) = C_{\text{PPC}} = NR\,\frac{c_{\text{sim}}}{N_{\text{parallel}}}. \tag{58}$$

In practice, some care is required in order to achieve this remarkably low computational cost. As discussed in Section 4 (in the context of a sequence of ABC samplers for a sequence of individual-specific posterior densities), the difficulty arises from individual-specific data arrays having different individual-specific vectors of auxiliary statistics. In general, whenever we have a sequence of individual-specific data arrays, $\mathbf{Y}_0^{(1)}, \ldots, \mathbf{Y}_0^{(m)}$, the vectors of auxiliary statistics, $\mathbf{U}_0^{(1)}, \ldots, \mathbf{U}_0^{(m)}$, will be different for each individual $i$ in the sequence. As a consequence of these differences in the auxiliary statistics, the simulations used to estimate the individual-specific posterior predictive p-value $P\left(\mathbf{Y}_0^{(i)}; \mathbf{U}_0^{(i)}\right)$ for individual $i$ (simulations generated from the joint posterior predictive density $\pi\left(\boldsymbol{\theta}, \mathbf{y}^{(i)} \,\middle|\, \mathbf{Y}_0^{(1)}, \ldots, \mathbf{Y}_0^{(s)}, \boldsymbol{u}\left(\mathbf{y}^{(i)}\right) = \mathbf{U}_0^{(i)}\right)$, conditional on the individual-specific vector of auxiliary statistics $\mathbf{U}_0^{(i)}$), can not be re-used directly to estimate the individual-specific posterior predictive p-value for some other individual.

Fortunately, in most cases, we can easily get around this problem by generating a single sample of $NR$ *master simulations*. From each of these master simulations, we can obtain a simulated data set satisfying any of the constraints represented by the individual-specific vectors of auxiliary statistics, $\mathbf{U}_0^{(1)}, \ldots, \mathbf{U}_0^{(m)}$, simply by censoring the simulated data arrays from the master simulations.

Recall (from Section 4) that in our analysis of HSC whole genome sequence data from 8 donor individuals, the data array $\mathbf{Y}_0^{(i)}$ from individual $i$ is a representation of a *sample phylogeny* on a sample of single cell genomes (together with mutation assignments to branches). Each simulation (performed using the *rsimpop* package, Williams et al. (2020)) returns a *population phylogeny*. Provided that the simulator is run for the age of the oldest donor individual in the sequence, the population phylogenies generated can be used as master simulations. From these master simulations we can obtain individual-specific simulations for individual $i$, which satisfy the constraints represented by $\mathbf{U}_0^{(i)} = \left(n^{(i)}, a^{(i)}\right)$, by terminating the population phylogeny at the age $a^{(i)}$, and deleting terminal nodes to obtain a sample phylogeny on exactly $n^{(i)}$ single cell genomes.

Recall (from Section 4) that the total time cost of generating a samples of $N$ observations from the (approximate) multiple-individual posterior density $\pi^{*(s)}(\boldsymbol{\theta})$, by performing $s$ ABC regression steps in the sequential ABC regression algorithm (Algorithm 2), is $C_{\text{ABC Seq}}(s)$, which is given by equation 46. Therefore, the total time cost of of generating a samples of $N$ observations from the (approximate) multiple-individual posterior density $\pi^{*(s)}(\boldsymbol{\theta})$ (using Algorithm 2), and then estimating a sequence of $m$ individual-specific posterior predictive p-values (defined in equation 56) based on this multiple-individual posterior density, is

$$C_{\text{ABC Seq}}(s) + C_{\text{PPC Seq}}(m) = (sN_{\text{sim}} + NR)\ \frac{c_{\text{sim}}}{N_{\text{parallel}}}. \tag{59}$$

In our analysis of HSC whole genome sequence data from 8 donor individuals, we ran the sequential ABC regression algorithm (Algorithm 2) with $s = 4$ ABC regression steps, to generate a sample of $N = 2000$ observations from the multiple-individual posterior density (equation 55), conditional on the data from the 4 oldest donor individuals. At each ABC regression step, a sample of $N_{\text{sim}} = 100{,}000$ simulations was generated, of which $N = 2000$ observations were accepted ($P_\delta = N/N_{\text{sim}} = 0.02$). The regression adjustments were obtained by performing a ridge regression (using the R package *abc* (Csilléry et al., 2012)) on the re-scaled and logit-transformed parameter values in the sample of accepted observations.

For all $m = 8$ individuals, we then estimated the individual-specific posterior predictive p-value, using the sample of at least $N = 200$ observations (sampled without replacement from the 2000 accepted observations (drawn from the approximate

**Algorithm 3** PPC for sequential ABC

**for** $j$ in $1{:}N$ **do**
  $\boldsymbol{\Theta}_j = \boldsymbol{\Theta}_j^{(s)}$;
  **for** $k$ in $1{:}R$ **do**
    Simulate individual-specific data array $\mathbf{Y}_{j,k} \sim p\left(\mathbf{y}^{(1)}; \boldsymbol{\Theta}_j\right)$;
    **for** $i$ in $1{:}m$ **do**
      From the individual-specific data array $\mathbf{Y}_{j,k}$
      construct (by censoring) an individual-specific data array $\mathbf{Y}_{j,k}^{(i)}$
      which satisfies condition $\boldsymbol{u}\left(\mathbf{Y}_{j,k}^{(i)}\right) = \mathbf{U}_0^{(i)}$;
      Compute vector of summary statistics $\mathbf{T}_{j,k}^{(i)} = \boldsymbol{t}\left(\mathbf{Y}_{j,k}^{(i)}\right)$;
    **end for**
  **end for**
  **for** $i$ in $1{:}m$ **do**
    From the sample of vectors $\mathbf{T}_{j,k}^{(i)}$ $(k = 1, \ldots, R)$:
    Compute vector of estimated means $\hat{\boldsymbol{M}}^{(i)}\left(\boldsymbol{\Theta}_j\right)$ of summary statistics;
    Compute vector of estimated standard deviations $\hat{\boldsymbol{S}}^{(i)}\left(\boldsymbol{\Theta}_j\right)$
    of summary statistics;
    Compute the *observed* discrepancy $\hat{D}\left(\mathbf{Y}_0^{(i)}; \boldsymbol{\Theta}_j\right)$;
    **for** $k$ in $1{:}R$ **do**
      Compute the *simulated* discrepancy $\hat{D}\left(\mathbf{Y}_{j,k}^{(i)}; \boldsymbol{\Theta}_j\right)$;
    **end for**
  **end for**
**end for**
**for** $i$ in $1{:}m$ **do**
  Compute estimate of posterior predictive p-value estimate $P\left(\mathbf{Y}_0^{(i)}; \mathbf{U}_0^{(i)}\right)$
  using equation 31;
**end for**

multiple-individual posterior density), and at least $R = 500$ re-simulations for each accepted observation, making a total of at least $NR = 100{,}000$ simulations. Therefore, using the estimate from equation 59, the total time cost of running the sequential ABC regression algorithm (with $s = 4$), and then estimating a sequence of $m$ individual-specific posterior predictive p-values (each one based on the same multiple-individual posterior density), is of order

$$\begin{aligned} C_{\text{ABC Seq}}(s) + C_{\text{PPC Seq}}(m) &= \left(4 \times 10^5 + 10^5\right) \frac{c_{\text{sim}}}{N_{\text{parallel}}} \\ &= 5 \times 10^5 \frac{c_{\text{sim}}}{N_{\text{parallel}}}. \end{aligned} \tag{60}$$
